# Towards reliable detection of introgression in the presence of among-species rate variation

**DOI:** 10.1101/2023.05.21.541635

**Authors:** Thore Koppetsch, Milan Malinsky, Michael Matschiner

## Abstract

The role of interspecific hybridization has recently seen increasing attention, especially in the context of diversification dynamics. Genomic research has now made it abundantly clear that both hybridization and introgression – the exchange of genetic material through hybridization and backcrossing – are far more common than previously thought. Besides cases of ongoing or recent genetic exchange between taxa, an increasing number of studies report “ancient introgression” – referring to results of hybridization that took place in the distant past. However, it is not clear whether commonly used methods for the detection of introgression are applicable to such old systems, given that most of these methods were originally developed for analyses at the level of populations and recently diverged species, affected by recent or ongoing genetic exchange. In particular, the assumption of constant evolutionary rates, which is implicit in many commonly used approaches, is more likely to be violated as evolutionary divergence increases. To test the limitations of introgression detection methods when being applied to old systems, we simulated thousands of genomic datasets under a wide range of settings, with varying degrees of among-species rate variation and introgression. Using these simulated datasets, we showed that some commonly applied statistical methods, including the *D*-statistic and certain tests based on sets of local phylogenetic trees, can produce false-positive signals of introgression between divergent taxa that have different rates of evolution. These misleading signals are caused by the presence of homoplasies occurring at different rates in different lineages. To distinguish between the patterns caused by rate variation and genuine introgression, we developed a new test that is based on the expected clustering of introgressed sites along the genome, and implemented this test in the program Dsuite.

---

Recent research has demonstrated that hybridization – the production of viable offspring between distinct species – is far more common than previously thought (Mallet, 2005; Taylor and Larson, 2019). Hybridization seems to be particularly frequent in rapidly diversifying clades (Meier et al., 2017; Patton et al., 2020; Mitchell and Whitney, 2021) and has also been linked to the emergence of new species through so-called hybrid speciation (Rieseberg et al., 1995; Lamichhaney et al., 2018; Runemark et al., 2018). Hybridization therefore appears to promote diversification in certain situations (Seehausen, 2004; Abbott et al., 2013), contrary to the traditional view in which hybridization is seen as inhibiting speciation (Mayr, 1942).

Recent studies have also revealed that even highly divergent species are sometimes still able to hybridize and backcross. Apart from records of interspecific hybrids within a genus, such as crosses between fin whale *Balaenoptera physalus* and blue whale *B. musculus* (Pampoulie et al., 2021), also intergeneric hybrids are known, for example between colubrid snakes of the genera *Pituophis* and *Pantherophis* (LeClere et al., 2012). Various other hybridization events between deeply divergent lineages have been reported, as for example among coral reef fishes (Pomacanthidae) with over 10% mitochondrial divergence (Tea et al., 2020), even though the most extreme examples of hybridization between divergent lineages are known from captive specimens only (e.g., interfamilial hybrids between sturgeons, Acipenseridae, and paddlefishes, Polyodontidae; Káldy et al. 2020). While the examples listed above refer to recent hybridization events, often detected through the observation of F1-hybrids, the fact that hybridization is recorded among divergent groups today suggests that it has also taken place in the distant past, when they were still more closely related.

Introgression, the transfer of genetic material between species, can leave detectable traces in the genomes of extant taxa. Such traces are being reported from an increasing number of taxa, including highly divergent ones, and have been interpreted as evidence for “ancient introgression”. Such ancient introgression has for example been reported to have occurred between the Komodo dragon *Varanus komodoensis* and Australian monitor lizards (Varanidae) in the Late Miocene (11.6–5.3 million years ago; Ma) (Pavón-Vázquez et al., 2021), among North American darters (Percidae, e.g., the genus *Allohistium*) at least 20 Ma (MacGuigan and Near, 2019), or among sea turtles (Cheloniidae) (Vilaça et al., 2021) up to 46 Ma. In fungus gnats, germline-restricted genes were suggested to have introgressed between the ancestors of Sciaridae and Cecidomyiidae even as early as 114 Ma (Hodson et al., 2022). In plants, ancient introgression has been reported for several groups of angiosperms (Stull et al., 2023). For example, birch tree species within Coryloideae (Betulaceae) were reported to have exchanged genes between 17 and 33 Ma (Wang et al., 2022; Stull et al., 2023) and ancient hybridization has been reported during the early diversification of asterids over 100 Ma, between the order Ericales and the ancestor of Cornales or Gentianidae (Stull et al., 2020, 2023).

These reports raise the question whether methods for the detection of introgression from genomic data are still applicable to such old groups (Hibbins and Hahn, 2022), given that key methods were originally developed for analyses at the level of populations and recently diverged species. One of the most commonly used approaches for introgression detection is the *D*-statistic, which was first applied to assess genetic exchange between Neanderthals and the ancestors of modern humans (Green et al., 2010). The *D*-statistic detects introgression through the so-called ‘ABBA-BABA test’ (Green et al., 2010; Durand et al., 2011), based on an imbalance in the sharing of ancestral (‘A’) and derived (‘B’) alleles across the genomes of four populations or species. This test assumes that, in the absence of introgression but presence of incomplete lineage sorting (ILS), two sister species share an equal proportion of derived ‘B’ alleles with any third species. A statistically significant excess of allele sharing in either direction (an excess of ‘ABBA’ or ‘BABA’ sites) is then considered indicative of genetic exchange between non-sister taxa. Although misleading signals can under certain scenarios be created by population structure in ancestral species (Durand et al., 2011; Eriksson and Manica, 2012), the *D*-statistic is considered to be robust under a wide range of evolutionary scenarios when applied to genome-wide data (Zheng and Janke, 2018).

However, the violation of two assumptions that are implicit in the use of the *D*-statistic can lead to false positive results: First, each variable site is assumed to result from a single substitution, and thus homoplasies – caused by independent substitutions at the same site in different species – are assumed to be absent. Randomly occurring homoplasies would not produce a false signal of introgression, because they are equally likely to increase the numbers of ‘ABBA’ and ‘BABA’ sites. Thus, a substitution that occurs in an outgroup to two sister species is equally likely to also occur in one or the other of the two sisters. But when a second assumption – that of uniform substitution rates across all species – is violated, homoplasies are more likely to occur in the sister species with the higher rate. This could lead to significantly unequal numbers of ‘ABBA’ and ‘BABA’ sites and a *D*-statistic falsely supporting introgression (Pease and Hahn, 2015; Amos, 2020; Frankel and Ańe, 2023).

Both violations, homoplasies and substitution-rate variation, are more likely to occur in older groups of species. Homoplasies require that sites are substituted on two different branches of a phylogenetic tree, which occurs more often when these branches are longer. Substitution-rate variation, on the other hand, is influenced by factors such as metabolic rate, generation time, longevity, or temperature, that are all expected to be similar among closely related species but may vary with increasing phylogenetic distance (Wilson Sayres et al., 2011; Bromham, 2020; Hua and Bromham, 2017; Ivan et al., 2022; Hua et al., 2015). A misleading effect of substitution-rate variation on the *D*-statistic, generating false-positive signals of introgression, has been suspected repeatedly (Pease and Hahn, 2015; Zheng and Janke, 2018; Hibbins and Hahn, 2022) and was recently supported by simulations under the birth-death-hybridization process (Justison et al., 2023; Frankel and Ańe, 2023).

To avoid the effects of rate variation on introgression detection, a tree-based equivalent of the *D*-statistic has been used in several studies (Vanderpool et al., 2020; Ronco et al., 2021). In this approach, rooted phylogenetic trees are first built for a large number of loci (regions with hundreds to thousands of base pairs) across the genome, and the inferred set of trees is then analyzed for topological asymmetry in three-species subsets just like site patterns are in the *D*-statistic. Thus, the most frequent tree topology for a set of three species is assumed to represent their species tree, and the frequencies of the second- and third-most frequent topologies are compared to each other. A significant difference in these frequencies is then interpreted as evidence of introgression. The test statistic has been named *D*_tree_ in Ronco et al. (2021) (who were unaware that a non-normalized version of this statistic had already been called Δ by Huson et al. 2005). Frequencies of tree topologies have also been used to infer introgression in other studies (Schumer et al., 2016; Gante et al., 2016; Figueiró et al., 2017; Martin and Van Belleghem, 2017; Suvorov et al., 2022). One might expect that, as a tree-based alternative to the *D*-statistic, *D*_tree_ would be more robust to homoplasies, given that the occurrence of one or few homoplasies per locus should not have an effect on the tree topology (Hibbins and Hahn, 2022; Frankel and Ańe, 2023). On the other hand, homoplasies in combination with rate variation can lead to long-branch attraction (Felsenstein, 1978), which might bias tree-topology frequencies even if their effect on each individual tree is weak.

Here, we use simulations to test the robustness of introgression detection methods to the combined effects of homoplasies and rate variation, as expected to occur in older groups of species. We simulate genomic datasets under a wide range of settings, including varying population sizes, divergence times, recombination rates, mutation rates, introgression rates, and degrees of among-species rate variation. Besides the *D*-statistic and its tree-based equivalent *D*_tree_, we apply three further tree-based methods to detect introgression in complementary ways: the phylogenetic network approach implemented in SNaQ (Soĺıs-Lemus et al., 2017), the approach based on branch-length distributions implemented in QuIBL (Edelman et al., 2019), and a method based on divergence-time distributions in time-calibrated phylogenies. The latter method was presented by Meyer, Matschiner, and Salzburger (2017), and will henceforth be called “MMS17 method”. We hypothesized that all of these methods could produce false signals of introgression when among-species rate variation is present, and that these signals would become stronger with increasing age of the introgression event, mutation rates, and degree of rate variation. Our results confirm that the *D*-statistic, as well as some of the tested tree-based methods are affected by rate variation. To distinguish between true signals of introgression and the false signals resulting from rate variation, we developed a new test based on the distribution of ‘ABBA’ sites on the genome, and we implemented this test into the introgression analysis software Dsuite (Malinsky et al., 2021). We assess the performance of this new test with simulated and empirical datasets, and confirm its suitability across a broad range of parameters.

## Materials and Methods

### Simulations

To test the performance of commonly applied introgression detection methods, genomic data were simulated under diverse scenarios. All simulations were conducted in the Python version 3.8.6 environment using the program msprime v.1.0 (Baumdicker et al., 2021). A four-taxon phylogeny was defined for the species P1, P2, P3, and P4, in which P1 and P2 were sister species and P4 was the outgroup to all others. The divergence of species P1 and P2, *t*_P1,P2_, was set to occur either 10, 20, or 30 million generations ago, with species P3 and P4 in each case set to branch off 10 and 20 million generations earlier, respectively. Thus, the most recent common ancestor of the four species dated to between 30 and 50 million generations ago (Fig. 1), and the internode distances were in all cases identical, which implied that the expected degree of ILS remained identical. All simulated species had identical and constant effective population sizes (*N*_e_), set to either *N*_e_ = 10^4^ or *N*_e_ = 10^5^ in separate simulations. An effective population size of *N*_e_ = 10^6^ was used in exploratory simulations, but as these simulations were too computationally demanding and their results did not seem to differ from those based on smaller population sizes, final simulations were based on the two smaller population sizes. We conducted one set of simulations that did not include any genetic exchange between species while other simulations included introgression between species P2 and P3. In these cases, P2 and P3 exchanged migrants with a rate *m* of either *m* = 10*^-^*^9^ (“very weak”), *m* = 10*^-^*^8^ (“weak”), *m* = 10*^-^*^7^ (“strong”), or *m* = 10*^-^*^6^ (“very strong”) per individual per generation, which is equivalent to the exchange of one migrant on average every 10^2^ *-* 10^5^ generations when *N*_e_ = 10^4^ or every 10 *-* 10^4^ generations when *N*_e_ = 10^5^. In all simulations, migration between P2 and P3 occurred for the same period of time, beginning with the divergence of P1 and P2 and ending 2.5 million generations later (Fig. 1).

**Fig. 1.**
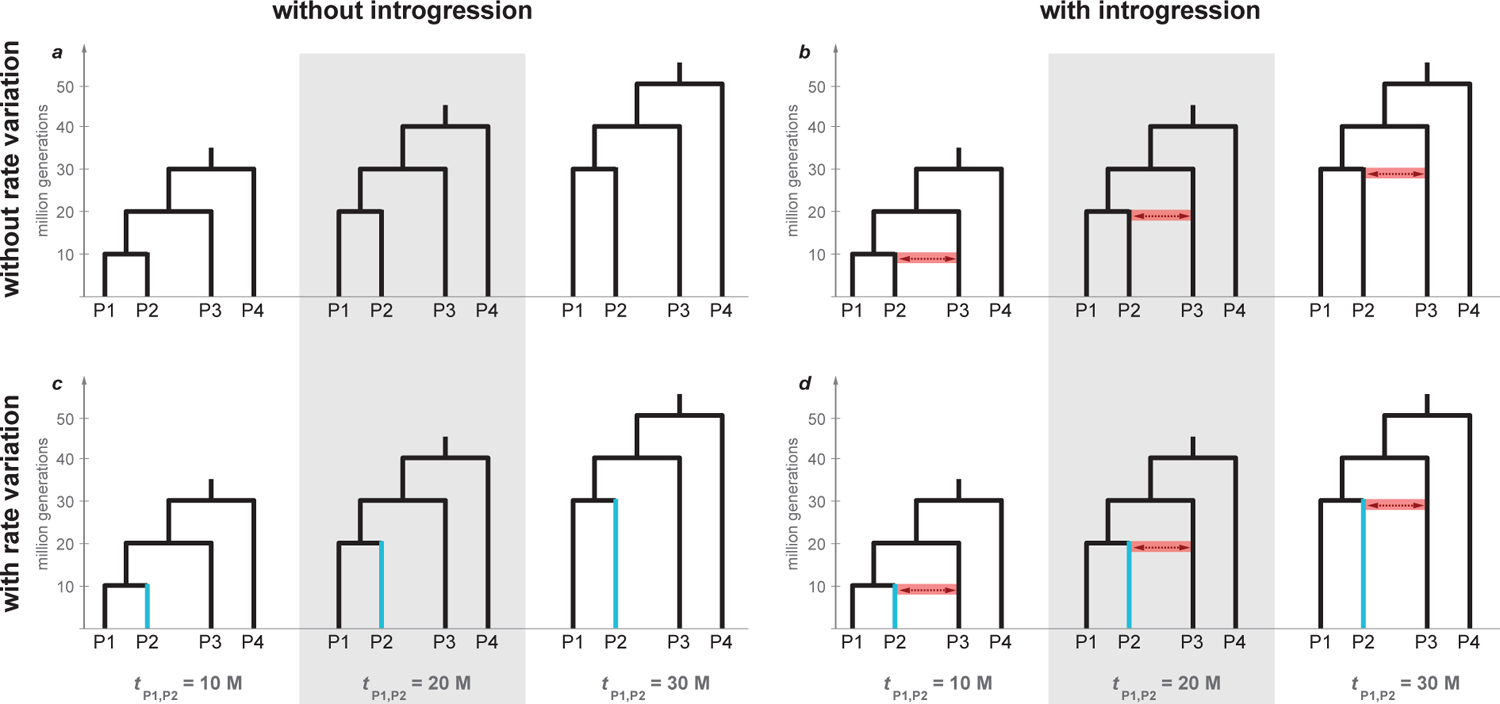
Four-taxon phylogenies used in simulations. Divergence times of species P1 and P2 (t_P1,P2_) are 10, 20, and 30 million generations in the past, with P3 and P4 branching off 10 and 20 million generations earlier, respectively. Species P2 evolved with a mutation rate that was either unchanged (scale factor s = 1; **a,b**) or slower (s = 0.25; illustrated in blue in **c,d**) than the mutation rate of all other species (besides s = 0.25, both a less extremely reduced rate and faster rates of species P2 were simulated with s = 0.5, s = 2, and s = 4, but are not shown here). In simulations that included introgression (**b,d**), this introgression occurred symmetrically between P2 and P3, beginning with the divergence of P1 and P2, and continuing for 2.5 million generations (illustrated in red in **b,d**). Any reliable method for introgression detection should identify a signal for **b** and **d** but not for **a** and **c**.

Based on this model of divergence and introgression, we simulated the evolution of the genomes of the four species, modeling these as a single chromosome with a length of 20 million basepairs (Mbp). The recombination rate *r* of this chromosome was set to *r* = 10*^-^*^8^ and the mutation rate *µ* was set to either *µ* = 10*^-^*^9^ or *µ* = 2 *⇥* 10*^-^*^9^ in separate simulations (both rates are given per site per generation). Mutations were simulated under the Hasegawa-Kishino-Yano (HKY) model (Hasegawa et al., 1985) with a transition-transversion rate ratio *k* = 2. Finally, we implemented among-species variation in the mutation rate to model a decreased, unchanged, or increased rate in species P2, with the rate change taking place immediately after its divergence from P1. Because msprime does not allow mutation-rate variation among species, we used a work-around with the same outcome, extending or shortening the branch leading to P2 with a scale factor *s*. We repeated the simulations using each of the five scale factors *s* = 0.25 (“very slow P2”), *s* = 0.5 (“slow P2”), *s* = 1 (“unchanged P2”), *s* = 2 (“fast P2”), and *s* = 4 (“very fast P2”) to model varying degrees of among-species rate variation. For each of the four simulated species, we sampled ten haploid chromosomes to form diploid genomes for five individuals per species. To implement the above-mentioned work-around for *s* = 0.25 and *s* = 0.5, we sampled individuals from species P2 at a time point in the past so that the length of its branch was effectively divided by 2 or 4. For *s* = 1, all samples were taken at the present. With scale factors *s* = 2 and *s* = 4, P2 was sampled at the present, but all divergences were shifted into the past by the amount of generations by which the P2 branch was extended, and P1, P3, and P4 were instead sampled in the past. In summary, we performed simulations with all possible combination of *t*_P1,P2_ *2 {*1 *⇥* 10^7^, 2 *⇥* 10^7^, 3 *⇥* 10^7^*}*, *N*_e_ *2 {*10^4^, 10^5^*}*, *m 2 {*0, 10*^-^*^9^, 10*^-^*^8^, 10*^-^*^7^, 10*^-^*^6^*}*, *r* = 10*^-^*^8^, *µ 2 {*1 *⇥* 10*^-^*^9^, 2 *⇥* 10*^-^*^9^*}*, and *s 2 {*0.25, 0.5, 1, 2, 4*}*; a total of 300 parameter combinations. For population size *N*_e_ = 10^5^, mutation rate *µ* = 2 *⇥* 10*^-^*^9^, introgression rate *m 2 {*0, 10*^-^*^8^, 10*^-^*^7^*}*, and a P2 branch rate *s 2 {*0.25, 1, 4*}*, 50 replicates (shown in Fig. 2–4) were simulated; for all the other parameter combinations, we performed ten replicate simulations (shown in Supplementary Figs. S1–S24), recording the resulting total genomic datasets in 4,080 files in the variant call format (VCF).

**Fig. 2.**
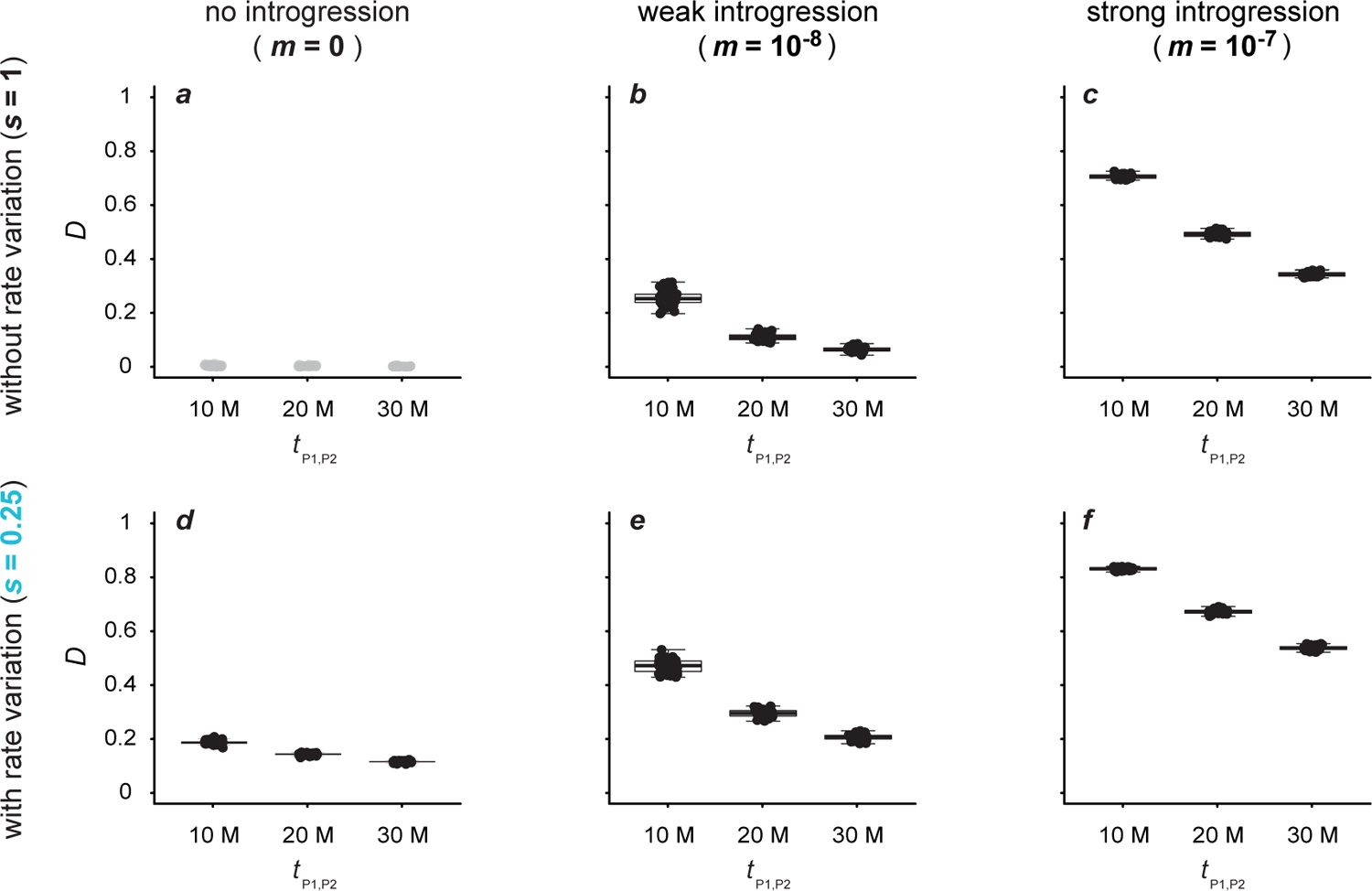
Patterson’s *D*-statistic for datasets simulated with a population size *N*e = 10^5^, a mutation rate *µ* = 2 *⇥* 10*^-^*^9^, either no (*m* = 0; **a,d**), weak (*m* = 10*^-^*^8^; **b,e**), or strong (*m* = 10*^-^*^7^; **c,f**) introgression, and either an unchanged (*s* = 1; **a**–**c**) or slow (*s* = 0.25; **d**–**f**) rate of branch P2. All results obtained with other settings are given in Supplementary Table S1 and illustrated in Supplementary Figures S1–S4. Per divergence time *t*P1,P2 *2 {*1 *⇥* 10^7^, 2 *⇥* 10^7^, 3 *⇥* 10^7^*}*, the *D*-statistic is shown for 50 replicate simulations. Circles in black indicate significant results (*p<* 0.05), and only these are summarized with box plots.

**Fig. 3.**
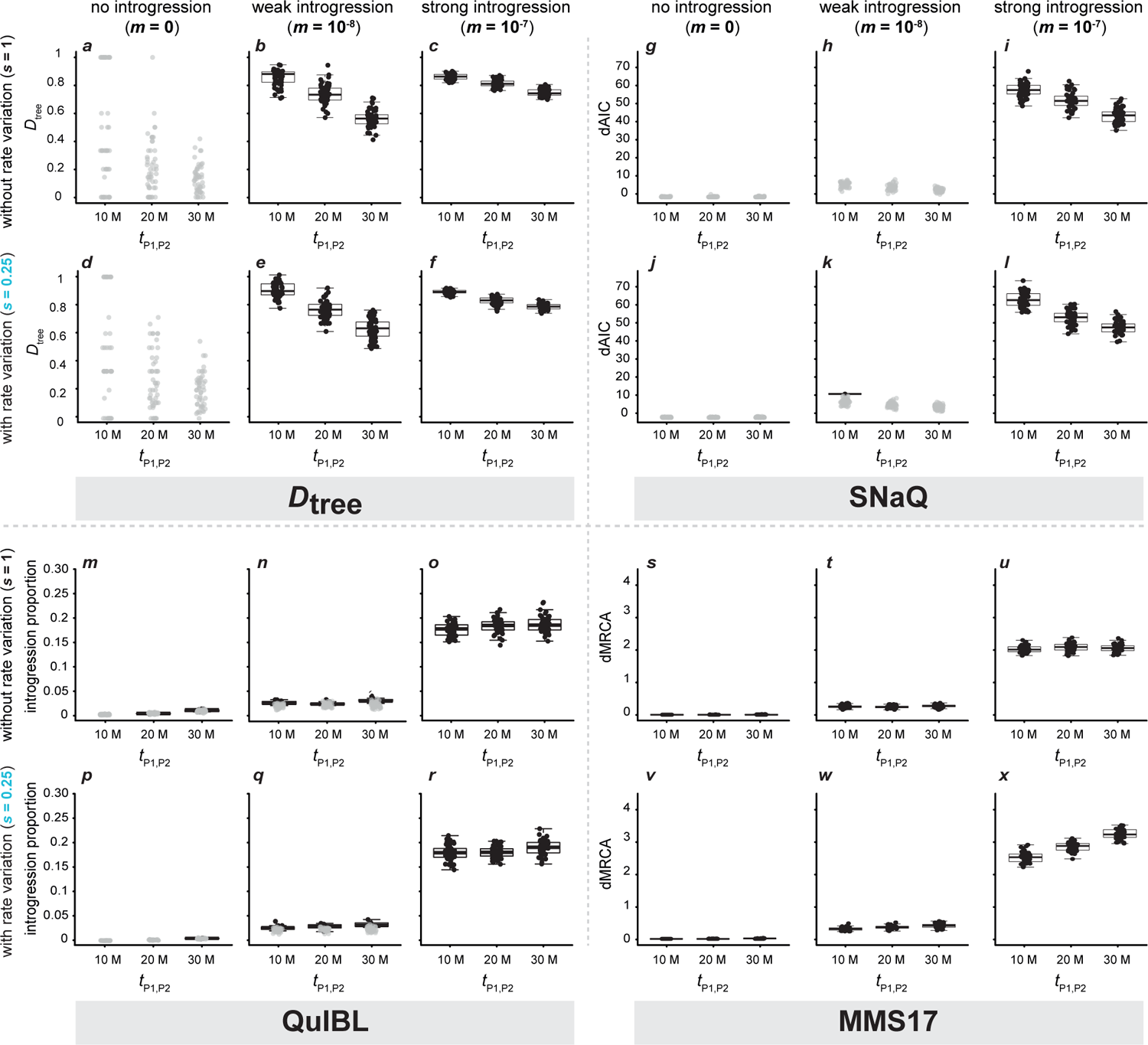
Signals of introgression detected with tree-based methods for datasets simulated with a population size *N*e = 10^5^, a mutation rate *µ* = 2*⇥*10*^-^*^9^, either no (*m* = 0), weak *m* = 10*^-^*^8^, or strong (*m* = 10*^-^*^7^) introgression, either an unchanged (*s* = 1) or very slow (*s* = 0.25) rate of branch P2, and an alignment length of 500 bp. All results obtained with other settings are shown in Supplementary Figures S5–S16. Per divergence time *t*P1,P2 *2 {*1*⇥*10^7^, 2*⇥*10^7^, 3*⇥*10^7^*}*, results are shown for 50 replicate simulations. Each result is based on 2,000 local trees. **a**–**f**) *D*tree; **g**–**l**) dAIC supporting introgression in networks produced with SNaQ; **m**–**r**) introgression proportion estimated with QuIBL; **s**–**x**) dMRCA estimated with the MMS17 method, in units of million generations. In **a**–**r**, circles in black indicate significant results (*p <* 0.05; before Bonferroni correction), and only these are summarized with box plots. As significance is not assessed with the MMS17 method, all values are shown in black in **s**–**x**.

**Fig. 4.**
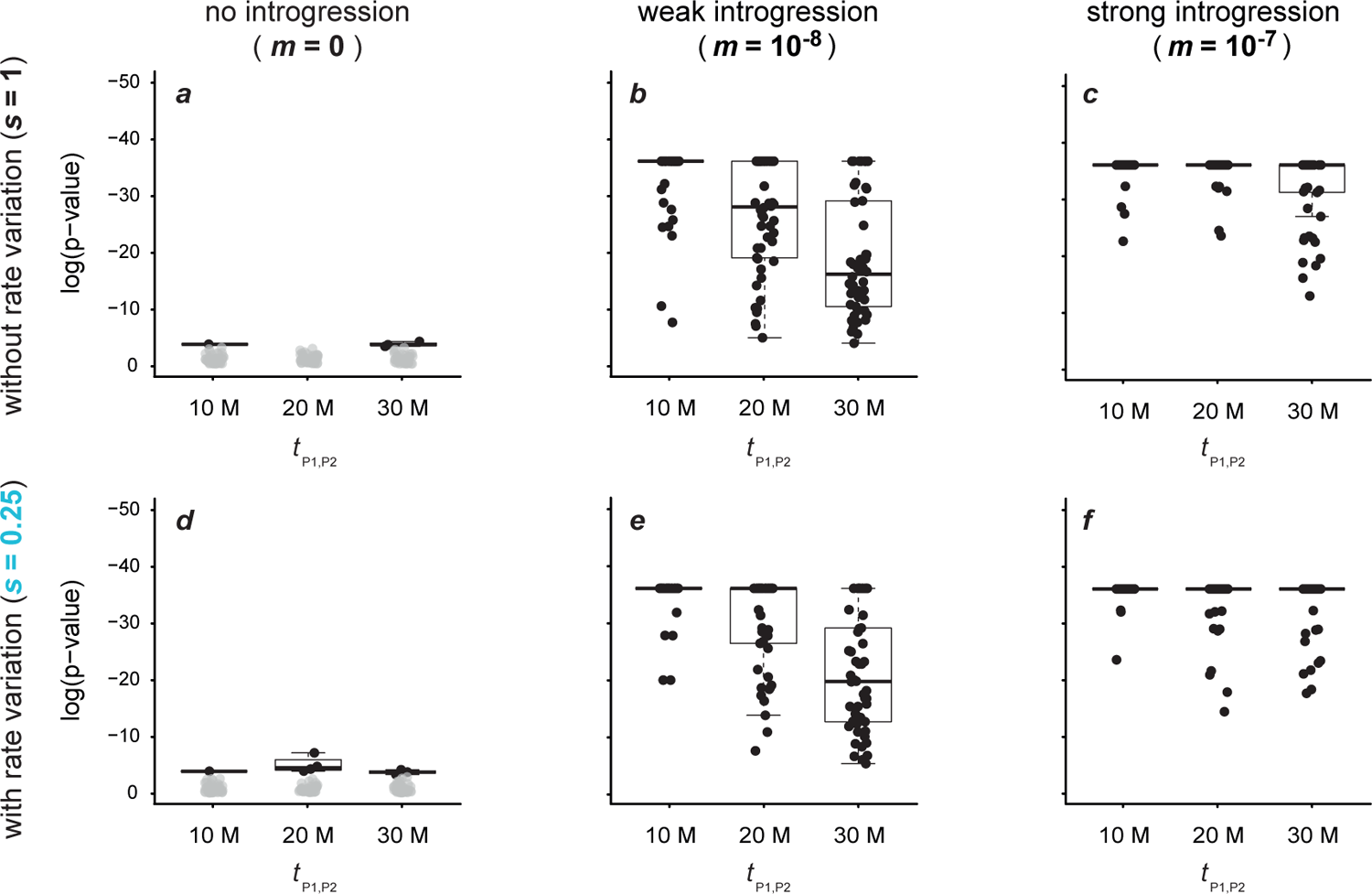
Signals of introgression detected with the “sensitive” version of the new ‘ABBA’-site clustering test for 50 replicate datasets simulated with a population size *N*e = 10^5^, a mutation rate *µ* = 2 *⇥* 10*^-^*^9^, either no (*m* = 0; **a,d**), weak (*m* = 10*^-^*^8^; **b,e**), or strong (*m* = 10*^-^*^7^; **c,f**) introgression, and either an unchanged (*s* = 1; **a**–**c**), or slow (*s* = 0.25; **d**–**f**) rate of branch P2. All results obtained with other settings are shown in Supplementary Figures S17–S21. Circles in black indicate significant results (*p <* 0.05), and only these are summarized with box plots. Significant results in **a** and **d** became non-significant after Bonferroni correction.

The range of parameters used in our simulations was selected to be comparable with some of the study systems for which ancient introgression has been reported. In terms of divergence time and mutation rate, our simulations are comparable to the example of North American darters (MacGuigan and Near, 2019): The divergence of the two genera *Allohistium* and *Simoperca*, for which signatures of ancient introgression have been reported, can be placed around 22 million generations ago, assuming a generation time of 1 year (Smith et al., 2011) and a divergence about 22 Ma (MacGuigan and Near, 2019). The mutation rates chosen for our simulations (*µ 2 {*10*^-^*^9^, 2 *⇥* 10*^-^*^9^*}*) also fall within estimates reported for darters as these range from around 6 *⇥* 10*^-^*^10^ to 9 *⇥* 10*^-^*^9^ per site and year (Smith et al., 2011).

### Patterson’s D-statistic

Patterson’s *D*-statistic (Green et al., 2010) measures signals of introgression in a species trio P1, P2, and P3 by counting the numbers of sites at which these species share alleles. Denoting ancestral alleles as ‘A’ and derived alleles as ‘B’, ‘ABBA’ sites are those at which P2 and P3 share the derived allele, while P1 and P3 share the derived allele at ‘BABA’ sites. By definition, the allele carried by the outgroup P4 is considered the ancestral allele ‘A’. The *D*-statistic is then calculated as the difference between the number of ‘ABBA’ sites *C*_ABBA_ and that of ‘BABA’ sites *C*_BABA_, normalized by the sum of these two numbers:

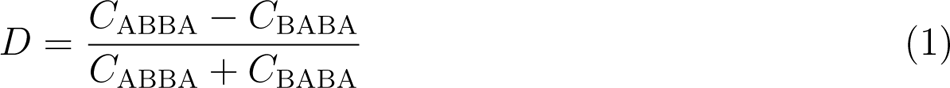

In the absence of introgression, *C*_ABBA_ and *C*_BABA_ are expected to be equal, in which case *D* = 0. However, this expectation is based on several assumptions, including that of equal rates for species P1 and P2, which we violated in part of our simulations. We therefore expected that the *D*-statistic would indicate signals of introgression (in the form of significant *p*-values) in these simulated datasets even when no introgression occurred.

We calculated the *D*-statistic for each of the 4,080 simulated genomic datasets with the program Dsuite v.r50 (Malinsky et al., 2021), using the program’s “Dtrios” module. By using the Dsuite implementation of the *D*-statistic, we were able to account for within-species variation in the calculation of *C*_ABBA_ and *C*_BABA_. When multiple individuals are sampled per species, Dsuite calculates *C*_ABBA_ and *C*_BABA_ based on the frequencies of the ancestral and derived alleles within the species. With the frequency of the derived allele ‘B’ at site *i* in the genome of species *j* denoted as *f*_B*,j,i*_ and a total number of sites *n*,

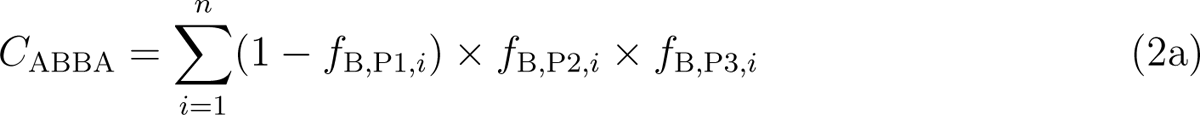

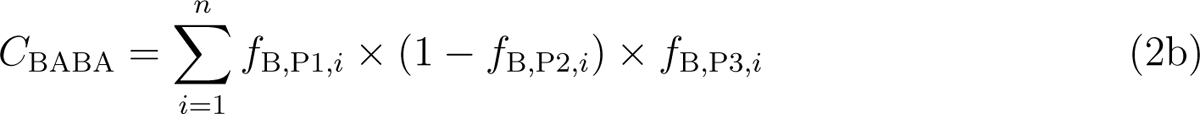

For both parts of Equation 2, Dsuite defines the derived allele as the one occurring at lower frequency in the outgroup P4 and multi-allelic sites are ignored. The significance of *D* was assessed with block jackknife tests, based on 20 equally sized subsets of each genomic dataset. In our interpretation of these results, we applied the Bonferroni correction (Bonferroni, 1935) to account for the large number of tests that we performed.

### Tree-based Introgression Detection Methods

Besides the *D*-statistic, we applied four tree-based introgression detection methods to all datasets simulated with a population size of *N*_e_ = 10^5^ and a mutation rate of *µ* = 2 *⇥* 10*^-^*^9^, a total of 1,830 datasets. To infer local trees as input for these methods, variants were extracted from equally spaced windows across the simulated chromosome. We separately extracted 5,000 windows of 200 bp, 2,000 windows of 500 bp, and 1,000 windows of 1,000 bp from each of the 1,830 datasets. These window sizes were chosen as a compromise between too little phylogenetic information in shorter windows and the occurrence of within-window recombination in larger windows, which could bias any phylogenetic inference (Bryant and Hahn, 2020). With these selected window sizes and numbers, only 1 Mbp out of the 20 Mbp of the simulated chromosomes was used for phylogenetic analyses. Additionally, per species and variable site, only the first allele of the first individual was extracted. All invariable sites within windows were replaced with randomly selected nucleotides A, C, G, and T, thus forming a sequence alignment for each window. By using only one allele of one individual per species – instead of both alleles of the five simulated individuals – we again reduced the amount of data by a factor of ten.

Consequently, for any given simulation, only 0.5% of the data used to calculate the *D*-statistic were also used for tree-based analyses. This data reduction was required due to the computational demands of our phylogenetic analyses: Because we used 1,830 genomic datasets in total and extracted 8,000 (5,000 + 2,000 + 1,000) windows from each of these, 14.64 million alignments were produced. As each alignment was used for phylogenetic analyses with both maximum-likelihood and Bayesian inference (see below), a total of 29.28 million such analyses were required.

*D*_tree_ *—* Conceptually similar to Patterson’s *D*-statistic, *D*_tree_ aims to detect introgression by comparing the counts of alternative rooted tree topologies for a given species trio, in a large set of local trees sampled across the genome. For any such trio P1, P2, and P3, three different rooted tree topologies can be found: One in which P1 and P2 are sister species, one in which P1 appears next to P3, and one in which P2 and P3 are sisters. Like for the *D*-statistic, the assumptions of no introgression and no among-species rate variation predict that, if the most frequent of these tree topologies represents the species-tree, the other two should occur in equal frequencies due to ILS. Any significant difference in the frequencies of the latter two topologies, assessed for example with a one-sided binomial test, can therefore be seen as support for introgression.

In its unconstrained version (Ronco et al. 2021; also see Vanderpool et al. 2020), *D*_tree_ is calculated from the counts of the second- and third-most frequent rooted topologies for the species trio, *C*_2nd_ and *C*_3rd_, as

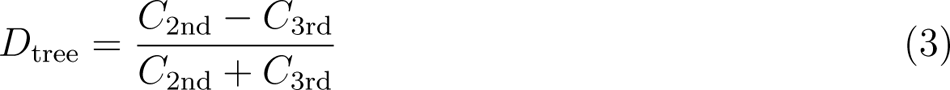

However, the use of this unconstrained version of *D*_tree_ may underestimate high levels of introgression when the most frequent tree topology of the three species does not reflect the species tree (due to very high levels of genetic exchange and/or very short internal branches). Therefore, we here applied a constrained version of *D*_tree_ to test explicitly for introgression between P2 and P3:

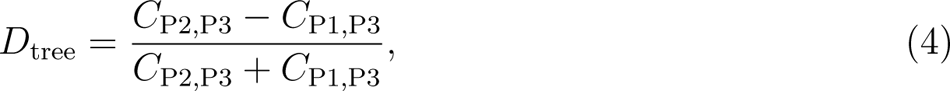

where *C*_P2,P3_ is the count of trees in which P2 and P3 are sisters, and *C*_P1,P3_ is the count of trees that place P1 and P3 next to each other.

To generate these counts, we inferred maximum-likelihood phylogenies from each window alignment of the 1,830 simulated genomic datasets using IQ-TREE v.2.1.2 (Minh et al., 2020), specifying the HKY substitution model (Hasegawa et al., 1985) and P4 as outgroup. The resulting tree sets were filtered by excluding trees with an internal branch shorter than 0.001 substitutions per site. We then obtained *C*_P1,P3_ and *C*_P2,P3_ by counting how often P1 and P3, or P2 and P3, respectively, were sister species in a set of trees. We did so separately for the sets of trees corresponding to each simulated genomic dataset and window size, by applying the Ruby script analyze tree asymmetry.rb (Ronco et al., 2021). Finally, a one-sided binomial test was used to identify whether *C*_P2,P3_ was significantly larger than *C*_P1,P3_ and thus supporting introgression between P2 and P3.

*SNaQ —* The SNaQ (Species Networks applying Quartets) method, implemented in PhyloNetworks (Soĺıs-Lemus and Ańe, 2016; Soĺıs-Lemus et al., 2017), is representative of a class of methods based on the multi-species coalescent model with hybridization (Meng and Kubatko, 2009). This class also includes approaches implemented in PhyloNet (Yu et al., 2014; Than et al., 2008; Yu and Nakhleh, 2015) or SpeciesNetwork (Zhang et al., 2018). From a set of local trees, SNaQ quantifies concordance factors for unrooted species quartets (either all possible quartets or a random sample) and calculates the likelihood for each of these quartets under the multi-species coalescent model with hybridization. By multiplying these likelihoods over all quartets, SNaQ derives the pseudolikelihood for a given species network. A heuristic search then allows SNaQ to estimate the network that optimizes the pseudolikelihood for a given maximum number of hybridization events. Thus, by repeating the SNaQ analysis with a maximum number of 0 and 1 such events, support for hybridization can be evaluated from the difference of the resulting pseudolikelihoods.

The multi-species coalescent model with hybridization considers hybridization events on the species level that instantaneously copy part of the genome from one species to another. Thus, this model is violated by our simulations in a way in which it may also be violated by most empirical cases of hybridizing species: Our simulations model hybridization between individuals that over long time scales (2.5 million generations) of ongoing introgression and subsequent drift, recombination, and occasional fixation has a gradual effect on the genomes of the recipient species. We expected that – barring other model violations – fitting such a period of hybridization and introgression to the multi-species coalescent model with hybridization would lead to the inference of a single hybridization event between species. Nevertheless, we expected that the support for this inferred hybridization event would correlate with the truth – i.e., the presence and the rate of introgression used in our simulations. We quantified this support as the difference in the Akaike information criterion (dAIC) (Akaike, 1974) for models that did or did not include a hybridization event, and considered dAIC values above 10 as significant. It has been pointed out that criteria like the Akaike information criterion are not generally suitable for the pseudolikelihoods estimated by SNaQ (Hibbins and Hahn, 2022). However this criterion is applicable in our case, because with no more than four species (i.e., a single quartet), SNaQ estimates the actual likelihood and not the pseudolikelihood (Soĺıs-Lemus and Ańe, 2016). We calculated the dAIC supporting introgression separately for each simulated genomic dataset and each of the three window sizes (200, 500, and 1,000 bp), based on the maximum-likelihood tree sets inferred for these windows with IQ-TREE, again excluding trees in which the internal branch was short (*<* 0.001 substitutions per site). We used PhyloNetworks v.0.14.2 for these analyses, providing the correct species tree as starting tree and specifying P4 as the outgroup when calling SNaQ.

*QuIBL —* QuIBL (Quantifying Introgression via Branch Lengths) is an approach to estimate proportions of introgressed loci based on the distribution of branch lengths in a species trio (Edelman et al., 2019). By using branch lengths as a source of information, QuIBL is complementary to SNaQ, as the latter is informed exclusively by the topologies of a set of local trees. All species trios in a given dataset are used by QuIBL and examined independently of each other. Per trio, QuIBL sorts the set of local trees into three subsets, one for each of the three possible rooted topologies of the triplet. For each of the three subsets, QuIBL then determines the distribution of the lengths of the internal branch (in numbers of substitutions per site), across all of the local trees within the subset. Applying an expectation-maximization algorithm in combination with the Bayesian information criterion (BIC) (Schwarz, 1978), it uses the shape of these distributions to determine whether they result from a single process (i.e., ILS) or additionally from a second process. This second process is interpreted either as lineage sorting within a common ancestor or introgression, depending on the relationships of the three species in a predetermined species tree. In the latter case, the number of local trees in the respective subset, multiplied by the proportion of them assigned to introgression rather than ILS, quantifies the overall introgression proportion. Like the multi-species coalescent model with hybridization implemented in SNaQ, the assumptions behind QuIBL also include a single pulse of hybridization instead of continuous introgression over a period of time (Edelman et al., 2019).

We applied QuIBL to the filtered sets of local trees generated using IQ-TREE for the 1,830 genomic datasets and each of the three window sizes. The QuIBL parameters included a likelihood precision treshold (“likelihoodthresh”) of 0.01, a limit of 50 steps for the expectation-maximization algorithm (“numsteps”), and a scale factor of 0.5 to reduce the step size when an ascent step fails (“gradascentscalar”), as recommended by the authors. We further specified P4 as the outgroup to the trio formed by P1, P2, and P3.

The results of QuIBL analyses were processed with the quiblR library (https://github.com/nbedelman/quiblR). Following Edelman et al. (2019), we considered support for introgression significant when the difference in BIC values (dBIC) was greater than 10.

*MMS17 method —* A fourth class of tree-based introgression detection methods uses distributions of divergence times in a set of ultrametric, time-calibrated local trees (Marcussen et al., 2014; Fontaine et al., 2015; Meyer et al., 2017). Of this class, we here apply the method developed by Meyer, Matschiner, and Salzburger (2017) (“MMS17 method”). This method compares the mean divergence times for all three possible pairs of species within a species trio, repeating this comparison for all possible species trios of a given dataset. For the species trio P1, P2, and P3, the mean ages of the most recent common ancestor (MRCA) of the pairs P1–P2, P1–P3, and P2–P3 are calculated over all local trees. If P1 and P2 are sister species and no introgression occurred with P3, the P1–P3 and P2–P3 mean MRCA age estimates are expected to be similar in the absence of introgression. In contrast, any introgression occurring between non-sister species should reduce one of these two mean MRCA ages (while increasing the P1–P2 mean MRCA age). The difference between these pairwise mean MRCA (dMRCA) ages is therefore informative about past introgression within the species trio – the larger dMRCA, the stronger the evidence for introgression (Meyer et al., 2017). On the other hand, the MMS17 method does not include a formal statistical test allowing one to reject the null hypothesis of no introgression. It has therefore been designed and used only to identify hypotheses of introgression that can then be tested with other methods (Meyer et al., 2017).

We used the Bayesian program BEAST2 v.2.6.4 (Bouckaert et al., 2019) to infer sets of time-calibrated local trees using the three alignment window sizes (200, 500, and 1,000 bp) for each of the 1,830 simulated genomic datasets. Per alignment, an input file for BEAST2 was produced with the babette library (Bilderbeek and Etienne, 2018), specifying the birth-death tree process as a tree prior (Gernhard, 2008) and the HKY substitution model (Hasegawa et al., 1985). Each tree was time-calibrated with a strict-clock model and an age constraint on the root. This constraint was defined as a log-normal prior distribution with a mean according to the true root age used in the simulation of the respective dataset (assuming a generation time of one year) and a narrow standard deviation of 0.001. Each BEAST2 analysis was performed with 5 million Markov-chain Monte Carlo iterations. Upon completion of each BEAST2 analysis, the resulting posterior tree distribution was summarized in the form of a maximum-clade-credibility tree with TreeAnnotator v.2.6.4 (Heled and Bouckaert, 2013). For each of the 1,830 genomic datasets and each window size, we used all produced summary trees jointly as input for the MMS17 method, as described above.

### ‘ABBA’-Site Clustering

In this manuscript, we propose a new test to discriminate between spurious and genuine signals of introgression based on clustering of ‘ABBA’ sites. This test aims to distinguish between homoplasies and introgressed sites, exploiting the fact that introgression typically leaves behind haplotypes with clusters of multiple linked variable sites that show the introgression pattern. On the other hand, homoplasies are expected to occur individually one by one. Our new “ABBA-site clustering” test therefore examines whether the ‘ABBA’ sites that are used for the *D*-statistic cluster among variable sites along chromosomes – which would support introgression – or whether they are distributed homogeneously as expected of homoplasies (although homoplasies can show limited clustering as a result of mutation-rate variation along the genome; see below).

As a first step, we identify “strong ABBA sites” for which most of the individuals in the dataset support the ‘ABBA’ pattern. Formally, these are sites for which where *f*_B,P1_, *f*_B,P2_, *f*_B,P3_, and *f*_B,P4_ are the frequencies of the derived allele ‘B’ in populations P1, P2, P3, and the outgroup (see Equation 4 a in Malinsky et al. 2021). We then test for clustering of these sites along chromosomes in two ways, the first of which is more sensitive, while the other one is robust to mutation-rate variation along the genome.

For the “sensitive” version of the test, we let *∼g* be a vector of all polymorphic sites on a chromosome or scaffold. We then define another vector *^∼^i*, where we record the indices of “strong ABBA sites” within *∼g*. For data from multiple chromosomes or scaffolds, vectors *∼g_c_* and *^∼^i_c_* are first calculated for each such unit *c* and then concatenated to form *∼g* and *^∼^i*.

We divide the values in *^∼^i* by the length of *∼g* (the number of polymorphic sites in the dataset), obtaining a normalized vector *^∼^i_n_* on the interval [0,1]. To test for clustering of “strong ABBA sites” we compare this normalized *^∼^i_n_* to the standard uniform distribution using a one-sample Kolmogorov-Smirnov test (Kolmogorov, 1933; Simard and L’Ecuyer, 2011). A significant test statistic supports clustering of “strong ABBA sites” among polymorphic sites along chromosomes and thus provides additional support for interpreting any signal of introgression as being genuine.

Under certain circumstances (see Results), the “sensitive” test version can show a clustering of “strong ABBA sites” arising purely from variation in the mutation rate along the chromosome. Therefore, we developed a second, “robust” version of the ‘ABBA’-site-clustering test, in which we replace vector *∼g* with a vector *^∼^h* that includes not all polymorphic sites, but only “strong ABBA sites” and the analogously identified “strong BABA sites”. This test version is robust because local mutation-rate variation increases the frequencies of “strong ABBA sites” and “strong BABA sites” equally in mutation hotspots. On the other hand, this version of the test is less sensitive than the first version, especially in cases where there are few strong ‘BABA’ sites; thus, for example, this test version might not detect strong introgression in the absence of ILS.

We implemented both versions of this test in the software Dsuite, where they can be called jointly with the function “--ABBAclustering” of the Dtrio module (Malinsky et al., 2021). We then assessed the power and reliability of both test versions by applying them to all simulated genomic datasets.

As a further evaluation of the performance of the ‘ABBA’-site-clustering test, we also applied it to an empirical dataset that we expected to be free from introgression but characterized by ILS. Specifically, we used a subset of the single-nucleotide polymorphism (SNP) data of Ronco et al. (2021), based on Illumina sequencing for all *⇠* 250 cichlid fish species of Lake Tanganyika and mapping to the Nile tilapia reference assembly (Conte et al., 2017). As the investigation by Ronco et al. (2021) had shown, introgression has occurred frequently among cichlid species within the taxonomic tribes of the Lake Tanganyika cichlid radiation, but only rarely among species of different tribes. We therefore reduced the SNP dataset of Ronco et al. (2021) to individuals of up to five randomly selected species from each of the four tribes Boulengerochromini (monotypic, including only *Boulengerochromis microlepis*), Lamprologini, Cyprichromini, and Tropheini, considering only quartets with one species per tribe. To the best of our knowledge, introgression between these tribes has not previously been reported and appeared absent in the study of Ronco et al. (2021). After subsetting the SNP dataset to include only the selected species, newly monomorphic sites were removed with BCFtools v.1.17 (Li, 2011). Both versions of the ‘ABBA’-site clustering test were then applied to the resulting SNP data subsets with Dsuite’s Dtrio module (placing *B. microlepis* as outgroup), while also calculating the *D*-statistic and its significance.

## Results

### Simulations

The numbers of variable sites in simulated datasets ranged from 1.46–8.20 million (7.3–41.0%), depending primarily on the mutation rate *µ* and the divergence time *t*_P1,P2_ (Table 1). Between 35,000 and 1.3 million (0.175–6.5%) of these sites were multi-allelic. The alignments of lengths 200, 500, and 1,000 bp had mean numbers of variables sites between 14.6 and 410.2 (Table 1). Pairwise genetic distances between species (*d*_xy_) for datasets with a population size *N*_e_ = 10^5^, a mutation rate *µ* = 2 *⇥* 10*^-^*^9^, and a recombination rate *r* = 10*^-^*^8^ ranged from 0.03 to 0.08 among P1 and P2 (*d*_xy_(P1,P2)) with a very slow P2 rate (*s*=0.25), and from 0.09 to 0.25 with a very fast P2 rate (*s*=4) (see Supplementary Table S4).

**Table 1.** Numbers of variable sites in simulated genomic datasets and alignments. Alignments with lengths of 200, 500, and 1,000 bp were extracted from the genomic datasets and used for tree-based inference methods. *N*e: effective population size; *µ*: mutation rate; *t*P1,P2:divergence time. The specified minimum and maximum values represent mean values obtained for a specific combination of all simulation parameters, across all simulation replicates for this parameter combination (see Supplementary Table S1 for a comprehensive overview of the numbers of variable, biallelic, and multiallelic sites per simulated dataset, as well as the numbers of variable and parsimony-informative sites per alignment lengths of 200, 500, and 1,000 bp).

| $N_e$ | $\mu$ | $t_{P1,P2}$ | Variable sites<br>($\times 10^6$) | Multi-allelic sites<br>($\times 10^3$) | Variable sites in alignments | | |
| --- | --- | --- | --- | --- | --- | --- | --- |
|  |  |  |  |  | 200 bp | 500 bp | 1,000 bp |
| $10^4$ | $1 \times 10^{-9}$ | $1 \times 10^7$ | 1.46 – 2.27 | 35 – 85 | 14.6 – 22.7 | 36.5 – 56.8 | 73.2 – 113.5 |
| $10^4$ | $1 \times 10^{-9}$ | $2 \times 10^7$ | 2.06 – 3.47 | 69 – 205 | 20.6 – 34.7 | 51.4 – 86.8 | 102.9 – 173.5 |
| $10^4$ | $1 \times 10^{-9}$ | $3 \times 10^7$ | 2.63 – 4.59 | 115 – 368 | 26.3 – 45.9 | 65.7 – 114.6 | 131.4 – 229.5 |
| $10^4$ | $2 \times 10^{-9}$ | $1 \times 10^7$ | 2.82 – 4.28 | 133 – 318 | 28.1 – 42.8 | 70.4 – 107.1 | 140.5 – 214.0 |
| $10^4$ | $2 \times 10^{-9}$ | $2 \times 10^7$ | 3.90 – 6.33 | 261 – 737 | 39.0 – 63.3 | 97.4 – 158.4 | 194.9 – 316.8 |
| $10^4$ | $2 \times 10^{-9}$ | $3 \times 10^7$ | 4.92 – 8.12 | 428 – 1,281 | 49.2 – 81.2 | 123.0 – 203.1 | 246.0 – 405.9 |
| $10^5$ | $1 \times 10^{-9}$ | $1 \times 10^7$ | 1.54 – 2.33 | 39 – 90 | 15.4 – 23.3 | 38.5 – 58.3 | 77.0 – 116.6 |
| $10^5$ | $1 \times 10^{-9}$ | $2 \times 10^7$ | 2.12 – 3.53 | 74 – 211 | 21.2 – 35.3 | 53.2 – 88.2 | 106.1 – 176.4 |
| $10^5$ | $1 \times 10^{-9}$ | $3 \times 10^7$ | 2.70 – 4.64 | 121 – 377 | 27.0 – 46.4 | 67.4 – 116.1 | 134.8 – 232.1 |
| $10^5$ | $2 \times 10^{-9}$ | $1 \times 10^7$ | 2.96 – 4.39 | 147 – 335 | 29.6 – 43.9 | 73.9 – 109.7 | 147.8 – 219.5 |
| $10^5$ | $2 \times 10^{-9}$ | $2 \times 10^7$ | 4.02 – 6.43 | 279 – 762 | 40.2 – 64.3 | 100.4 – 160.7 | 201.0 – 321.4 |
| $10^5$ | $2 \times 10^{-9}$ | $3 \times 10^7$ | 5.03 – 8.20 | 448 – 1,311 | 50.3 – 82.0 | 125.6 – 205.1 | 251.5 – 410.2 |

The simulated data based on the divergence model in Figure 1 had very little or no ILS. While the mean lengths of chromosomal regions unbroken by recombination – termed “c-genes” by Doyle (1995) – were between 18 and 20 bp, the lengths of chromosomal regions sharing the same species topology (“single-topology tracts”) were far longer.

Without introgression (*m* = 0), almost all simulated chromosomes (58 out of 60 for which we made this assessment) had the same topology – that of the species tree – from beginning to end. The two exceptions were datasets simulated with the population size *N*_e_ = 10^5^ that included three and two single-topology tracts, respectively.

With introgression rates increasing from *m* = 10*^-^*^9^ to *m* = 10*^-^*^7^, the mean lengths of single-topology tracts decreased from a minimum of 64,516 bp (and a maximum of the chromosome length) to a range between 7,132 and 10,010 bp with a population size of *N*_e_ = 10^4^, and from 22,026–217,391 bp to 422–789 bp with *N*_e_ = 10^5^. However, with the highest simulated rate of introgression *m* = 10*^-^*^6^, the lengths of single-topology tracts mostly increased again, to 2,246–95,238 bp with *N*_e_ = 10^4^ and to 86–1,277 bp with *N*_e_ = 10^5^ (Supplementary Table S1). The reason for this was a dominance of regions with introgression in these chromosomes, causing them to form single-topology tracts.

While only 0–3.7% of the chromosome were affected by introgression with *m* = 10*^-^*^9^, these proportions grew to 1.4–10.0%, 30.6–46.5%, and 95.7–99.5% with *m* = 10*^-^*^8^, 10*^-^*^7^, and 10*^-^*^6^, respectively. Because of these extreme differences, we focus on the scenarios of weak (*m* = 10*^-^*^8^) and strong introgression (*m* = 10*^-^*^7^), besides the scenario without introgression (*m* = 0), in the remainder of the Results section. We present all results, including those obtained with very weak (*m* = 10*^-^*^9^) and very strong introgression (*m* = 10*^-^*^6^) in the Supplementary Material.

### Patterson’s D-statistic

As expected, Patterson’s *D*-statistic reliably indicated introgression when it was present and rate variation was absent (*s* = 1). With a population size *N*_e_ = 10^5^ and mutation rate *µ* = 2 *⇥* 10*^-^*^9^ (Fig. 2), the *D*-statistic was below 0.015 and insignificant (*p* > 0.05) for all replicate datasets when introgression was absent (*m* = 0), regardless of the divergence time *t*_P1,P2_ (Fig. 2a). With weak (*m* = 10*^-^*^8^) or strong introgression (*m* = 10*^-^*^7^), on the other hand, the *D*-statistic was in the ranges of 0.04–0.31 and 0.33–0.73, respectively, and in all cases highly significant (*p<* 10*^-^*^10^) (Fig. 2b,c). The *D*-statistic was lower (0–0.05) and in some cases not statistically significant in settings with very weak (*m* = 10*^-^*^9^) introgression, and higher (0.59–0.87) and always significant (*p<* 10*^-^*^16^) in settings with very strong (*m* = 10*^-^*^6^) introgression (Supplementary Fig. S2). In all cases, the *D*-statistic decreased with increasing age of the phylogeny (i.e. with *t*_P1,P2_), suggesting that both false and true signals of introgression would be even stronger in groups with younger divergences. This decrease with age was caused by homoplasies and reversals accumulating on the longer branches of the older phylogenies, augmenting both *C*_ABBA_ and *C*_BABA_ (Supplementary Note 1). Simulations with a lower population size (*N*_e_ = 10^4^) or a lower mutation rate (*µ* = 1 *⇥* 10*^-^*^9^) produced the same patterns (Supplementary Figs. S1, S3, and S4).

In contrast to the results obtained without rate variation, the *D*-statistic was not a reliable indicator of introgression when rate variation was present. While the *D*-statistic was significant for nearly all datasets simulated with introgression (Supplementary Figs. S1–S4), it was also significant for all datasets simulated without introgression (*m* = 0) whenever rate variation was present. In these cases, the *D*-statistic ranged from 0.05 to 0.21 (*p<* 4.4 *⇥* 10*^-^*^10^) (Fig. 2d; Supplementary Figs. S1–S4).

Like the decrease of the *D*-statistic with increasing age of the phylogeny, the false-positive signals of introgression were caused by homoplasies and reversals. This can be explained focusing on the results obtained with a very fast rate of the P2 branch (*s* = 4) on the youngest phylogeny (*t*_P1,P2_ = 10 million generations), shown in Supplementary Figure S2. The high *D*-statistic of 0.19–0.20 for these simulated datasets resulted from a *C*_ABBA_ in the range of 42,366–43,403 and a *C*_BABA_ around 28,756–29,355. Perhaps contrary to expectations, this *D*-statistic does not support introgression between P2 and P3, but instead between P1 and P3 (Dsuite automatically rotates P1 and P2 so that *D* > 0). A detailed analysis of one replicate simulation output explains this result: As expected, the faster rate of evolution of P2 led to more homoplasies shared between P2 and P3 (11,583; considering only bi-allelic sites) than between P1 and P3 (3,314). However, as the outgroup P4 had a longer branch than P3, this difference was more than compensated for by a greater number of homoplasies between P2 and P4 (16,856) compared to P1 and P4 (4,690). Additionally, far more reversals of substitutions in the common ancestor of P1, P2, and P3 occurred on the branch leading to P2 (5,990) than on that leading to P1 (1,697), further increasing allele sharing between P1 and P3. The remaining difference between *C*_ABBA_ and *C*_BABA_ may be explained by multi-allelic sites, of which there were 21,123.

The *D*-statistic was similarly high, in the range of 0.18–0.20, in datasets produced without introgression (*m* = 0) and a very slow rate of the P2 branch (*s* = 0.25), but it supported introgression between P2 and P3, not between P1 and P3, in these instances (Fig. 2d). As in the cases with an increased P2 rate, the imbalance between a *C*_ABBA_ of 11,565–12,098 and a *C*_BABA_ of 7,877–8,211 is explained by homoplasies and reversals: While P1 and P3 shared more homoplasies (3,282) than P2 and P3 (1,046), P1 also shared even more homoplasies with P4 (4,679; compared to 1,493 homoplasies shared between P2 and P4). Additionally, more reversals of substitutions in the common ancestor of P1, P2, and P3 occurred on the branch leading to P1 (1,637) compared to P2 (509), resulting in more allele sharing between P2 and P3 and thus the imbalance between *C*_ABBA_ and *C*_BABA_.

The false signals of introgression were not exclusive to the datasets simulated with extreme rate variation (*s* = 0.25 and *s* = 4), but also affected the datasets with more modest rate variation (*s* = 0.5 and *s* = 2). While the *D*-statistic was lower in these cases (0.05–0.14), it remained highly significant for all these datasets (*p<* 10*^-^*^9^) (Supplementary Figs. S1–S4).

### Tree-based Introgression Detection Methods

*D*_tree_ *—* Sets of maximum-likelihood trees, generated for the 1,830 simulated datasets, produced high *D*_tree_ values up to around 1, even when no introgression was present (Fig. 3a,d). This pattern did not seem to be affected by rate variation (Fig. 3d), and was found with the alignment lengths 200, 500, and 1,000 bp (Fig. 3a,d; Supplementary Figs. S5–S7). The applied binomial test did not support a significant difference (*p>* 0.05) between *C*_P2,P3_ and *C*_P1,P3_ in all cases without introgression. These high but non-significant *D*_tree_ values resulted from stochastic variation in the small numbers of trees that are not concordant with the species phylogeny. For example, with the youngest phylogeny (*t*_P1,P2_ = 10 million generations) and an alignment length of 500 bp, no more than 14 out of 2,000 trees were discordant for any of the 50 replicates with P2 branch rate *s 2 {*0.25, 1, 4*}*. With older phylogenies and the same alignment length, these numbers of discordant trees remained in the ranges of 4–46 and 17–139, for *t*_P1,P2_ = 20 and *t*_P1,P2_ = 30 million generations, respectively.

The lack of significance could indicate that unlike Patterson’s *D*-statistic, *D*_tree_ might be robust to rate variation. Alternatively, however, it could also result from the reduced amount of data used in tree-based analyses (covering only 1 Mbp of the 20 Mbp-chromosome). If rate variation combined with homoplasies would influence the ratio of *C*_P2,P3_ and *C*_P1,P3_, it is conceivable, that this becomes apparent only with larger numbers of discordant trees. To test whether the small numbers of discordant trees may hide a weak influence of rate variation, we compared the mean values for *C*_P2,P3_ and *C*_P1,P3_ across all replicates for settings with *s<* 1 and *s>* 1 (Supplementary Table S2). We expected that if rate variation affected *D*_tree_ in the same direction as Patterson’s *D*, the mean values of *C*_P2,P3_ should generally be larger than those for *C*_P1,P3_ when *s<* 1, and vice versa. Focusing only on those settings for which we had simulated 50 replicate datasets, this was in fact the case for 10 out of 12 settings (the two mean values being small and equal in the remaining two settings) (Supplementary Table S2). Thus, homoplasies and rate variation appear to influence topologies in the same direction as they influence site patterns. However their effect on tree topologies appears minimal, so that it can only be noticed when assessing a large number of replicate analyses jointly. Moreover, the influence of homoplasies and rate variation on *D*_tree_ was clearly far weaker than the effect of true introgression. When introgression was included in the simulations, its presence was reliably detected for migration rates *m* > 10*^-^*^8^, regardless of divergence time *t*_P1,P2_ or alignment length (Fig. 3; Supplementary Figs. S5–S7). Like Patterson’s *D*-statistic, *D*_tree_ values were decreasing with increasing divergence times (e.g., Fig. 3b). This was apparently caused by added stochasticity in tree topologies resulting from homoplasious substitutions, as both types of discordant trees became more frequent with older divergence times (Supplementary Table S3).

*SNaQ —* The maximum-likelihood values reported by SNaQ were in all cases equally good or better for the model that included a hybridization event, compared to the hybridization-free model (Supplementary Table S1). No effect of rate variation was recorded, and SNaQ correctly favored the model without hybridization when analyzing data simulated without introgression (Fig. 3g,j; Supplementary Figs. S8–S10). However, SNaQ had a low power to detect weak introgression (*m* 6 10*^-^*^8^) (Fig. 3h,k; Supplementary Figs. S8–S10). Only with a strong introgression rate in the simulations (*m* > 10*^-^*^7^) did SNaQ detect significant signals of it (e.g., Fig. 3i,l). The dAIC values ranged from 0.60 to 10.36 (with a single significant dAIC value *>* 10; Fig. 3k) when weak introgression (*m* = 10*^-^*^8^) was present (Fig. 3h,k), but increased to significant values between 39.43 and 73.06 with strong introgression (*m* = 10*^-^*^7^) (Fig. 3i,l). As with Patterson’s *D*-statistic or *D*_tree_, signals of introgression became weaker with increasing divergence times (Fig. 3i,l), probably because of the generally higher numbers of discordant trees inferred in those cases. The patterns described above were equally found with all tested alignment lengths (Supplementary Figs. S8–S10), and therefore seemed to be unaffected by it.

*QuIBL —* QuIBL produced signals of introgression even when neither rate variation nor introgression were present (*s* = 1, *m* = 0). Analyzing sets of trees generated for alignments of 500 bp, four out of 50 simulation replicates with *t*_P1,P2_ = 20 million generations and ten replicates with *t*_P1,P2_ = 30 million generations produced significant results (Fig. 3m). This changed dramatically for different alignment lengths. With trees produced under these settings (*s* = 1, *m* = 0) for alignments of 200 bp, QuIBL reported significant results for all ten replicates, regardless of phylogeny age (Supplementary Fig. S11). On the other hand, with alignments of 1,000 bp, none of the results were significant (Supplementary Fig. S13). Like the level of significance, the introgression proportion estimated by QuIBL was higher with shorter alignments; ranging from 0.01 to 0.05 with alignments of 200 bp, from 0 to 0.01 with alignments of 500 bp, and remaining around 0 when alignments of 1,000 bp were used (Supplementary Figs. S11–S13).

Adding rate variation while still excluding introgression (*m* = 0) led to fewer significant results with decreased rates of the P2 branch (*s<* 1); however, even more significant results were found for faster rates (*s>* 1) (Fig. 3m,p; Supplementary Fig. S12). With a very slow rate (*s* = 0.25) of the P2 branch, 12 of the 50 replicate tree sets for alignments of 500 bp produced significant results, though only those with *t*_P1,P2_ = 30 million generations (Fig. 3p). On the other hand, an increased rate of branch P2 (*s* = 4) led to even more significant false-positive signals of introgression, particularly for older phylogenies (2, 18, and 49 significant results out of 50 for *t*_P1,P2_ = 10, 20, and 30 million generations, respectively) (Supplementary Fig. S12). As before without rate variation, this pattern was affected by the length of the alignments used to produce the tree sets. With alignments of 200 bp, almost all analyses produced significant results, while alignments of 1,000 bp led to results that were in most cases non-significant (Supplementary Figs. S11 and S13).

When introgression was simulated with *m* > 10*^-^*^7^, QuIBL detected it reliably, but failed to detect it in most cases (478 out of 810 datasets) when *m* = 10*^-^*^8^. The introgression proportion was estimated between 0.01–0.08 with *m* = 10*^-^*^8^ and between 0.13 and 0.24 with *m* = 10*^-^*^7^, which was influenced only to a minor degree by rate variation (*s*), phylogeny age (*t*_P1,P2_), and alignment length (Fig. 3n,o,q,r; Supplementary Figs. S11–S13).

*MMS17 method —* The MMS17 method performed as expected when neither rate variation nor introgression were present (Fig. 3s; Supplementary Figs. S14–S16), with a difference between the two oldest pairwise mean MRCA ages (dMRCA) close to 0 (between 0 and 0.07 million generations). With increasing levels of introgression, dMRCA was continuously growing, to 0.14–0.42 million generations when *m* = 10*^-^*^8^ and 1.58–2.47 million generations when *m* = 10*^-^*^7^. Phylogeny age (*t*_P1,P2_) had no noticeable influence on dMRCA in these cases, but dMRCA was slightly higher with shorter alignments compared to longer ones (Pearson’s product-moment correlation, *p<* 0.001; Supplementary Figs. S14–S16).

However, when rate variation was simulated, the MMS17 method became rather unreliable, particularly with faster rates (*s* > 1) of the P2 branch (Supplementary Figs. S14–S16). With the very fast P2 rate (*s* = 4) and the youngest phylogeny (*t*_P1,P2_ = 10 million generations), dMRCA increased to values between 1.45 and 4.10 myr, again depending on alignment length (Pearson’s product-moment correlation, *p<* 0.001). These strong signals were the result of local trees in which P2 was incorrectly placed as the sister to a clade combining P1 and P3. As this placement allowed an extension of the P2 branch length, the inferred rate variation across the phylogeny was lowered, improving the prior probability of the tree in the strict-clock model. With the older phylogenies (*t*_P1,P2_ > 20 million generations) and the very fast rate for the P2 branch (*s* = 4), the two oldest mean pairwise MRCA ages were no longer those between P1 and P3 and between P2 and P3, leading to erroneous signals (Supplementary Figs. S14–S16).

In contrast, a slower rate (*s* 6 1) of the P2 branch did not have a strong influence on dMRCA (Fig. v). An increasing false signal of introgression with increasing age of the phylogeny could nevertheless be observed when the tree set was based on short alignments of 200 bp (Supplementary Fig. S14). In these cases, dMRCA ranged between 0.23 to 0.32 million generations.

### ‘ABBA’-Site Clustering

Across the tested parameter space, our new method based on ‘ABBA’-site clustering proved to be reliable in distinguishing false positives from genuine introgression signals (Figs. 4 and 5; Supplementary Figs. S17–S24). Applied to the datasets simulated without introgression (*m* = 0) and without rate variation (*s* = 1), the “robust” version of the test did not produce a single significant result (Fig. 5a; Supplementary Figs. S21–S24).

**Fig. 5.**
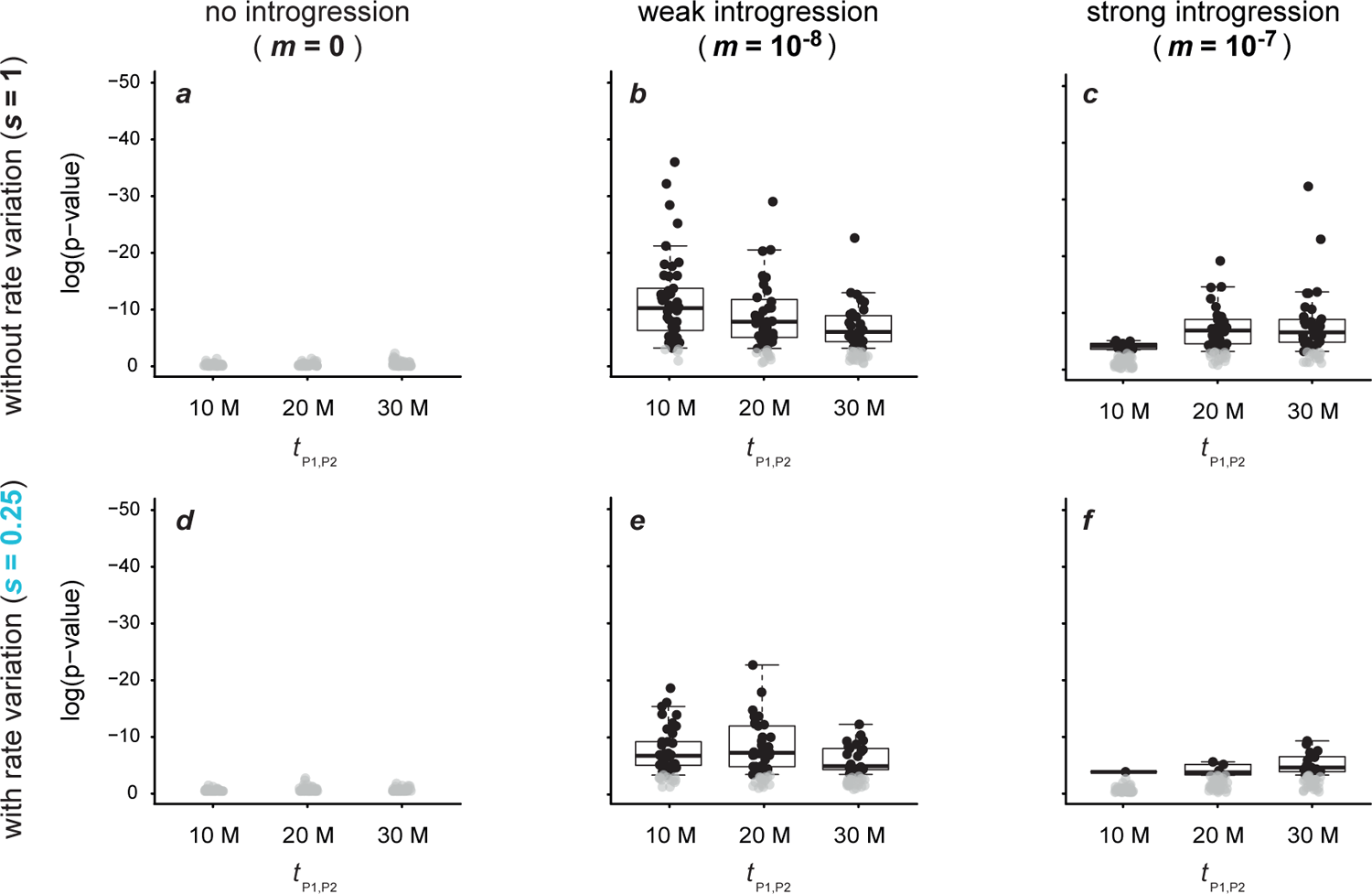
Signals of introgression detected with the robust version of the new ‘ABBA’-site clustering test for datasets simulated for 50 replicates with a population size *N*e = 10^5^, a mutation rate *µ* = 2 *⇥* 10*^-^*^9^, either no (*m* = 0; **a,d**), weak (*m* = 10*^-^*^8^; **b,e**), or strong (*m* = 10*^-^*^7^; **c,f**) introgression, and either an unchanged (*s* = 1; **a**–**c**), or slow (*s* = 0.25; **d**–**f**) rate of branch P2. All results obtained with other settings are shown in Supplementary Figures S21–S24. Circles in black indicate significant results (*p<* 0.05), and only these are summarized with box plots.

While the “sensitive” version returned for the same parameters weakly significant false-positive signals for up to 7 out of 240 datasets (*p>* 0.0017; Fig. 4a; Supplementary Figs. S17–S20), all of these became non-significant after Bonferroni correction.

Importantly, adding branch rate variation (*s 2 {*0.25, 0.5, 2, 4*}*) did not lead to false positives. There were weakly significant (*p>* 0.0002) signals for 37 out of 720 datasets with the “sensitive” test, all of which became non-significant after Bonferroni correction (Fig. 4d; Supplementary Figs. S17–S20). The “robust” test version again did not return a single false-positive (Fig. 5d; Supplementary Figs. S21–S24).

Similar results were obtained with a variable recombination rate, where three out of 90 datasets produced significant results with the “sensitive” test version (*p>* 0.01; all non-significant after Bonferroni correction; Supplementary Note S2, Supplementary Fig. S25) and none were significant with the “robust” test version (Supplementary Fig. S26). For increased levels of ILS (Supplementary Note S3; Supplementary Figs. S27–S32), 16 out of 360 significant values were recorded with the “sensitive” test version (*p>* 0.002) (Supplementary Figs. S27–S29), while a single significant value was recorded with the “robust” test version (*p* = 0.02) (Supplementary Figs. S30–S32). Again, all of these became non-significant after Bonferroni correction.

Next, we assessed whether mutation-rate variation along the chromosome could lead to clustering of ‘ABBA’ sites and thus to false-positive signals in our new test. To this end, we performed an additional set of simulations (Supplementary Note S4) with among-site mutation-rate variation, and applied both versions of the ‘ABBA’-site-clustering test to these additional datasets. The presence of among-site mutation-rate variation led to some false positives in the “sensitive” version of the test (Supplementary Fig. S33). Of 30 datasets simulated with neither introgression (*m* = 0) nor among-species rate variation (*s* = 1), 10 yielded significant signals of ‘ABBA’-site clustering (*p* > 0.00008), and one of these remained significant after Bonferroni correction. Adding among-species rate variation (*s 2 {*0.25, 4*}*) led to similar results (Supplementary Fig. S33). Of the 60 datasets simulated with these settings, 15 produced significant results (*p* > 3 *⇥* 10*^-^*^7^) and two of these remained significant after Bonferroni correction. In contrast, the among-site mutation rate variation did not influence the “robust” version of the test, producing not a single significant results when introgression was excluded (Supplementary Fig. S34).

The presence of introgression led to significant ‘ABBA’-site clustering for a large majority of simulated datasets (Figs. 4,5; Supplementary Figs. S17–S24). The “sensitive” version of the test was always significant for strong (*m* > 10*^-^*^7^) and very strong rates (*m* > 10*^-^*^6^) of introgression (Fig. 4f; Supplementary Figs. S17–S20). All false negatives – cases that did not lead to significant clustering despite the presence of introgression – were limited to settings where the P2 rate was increased (*s>* 1) and introgression was weak (*m* = 10*^-^*^8^) or very weak (*m* = 10*^-^*^9^) (305 out of 600; for *N*_e_ *2 {*10^4^,10^5^*}*) (Supplementary Figs. S17–S20). In contrast to the “sensitive” version, the “robust” version of the ‘ABBA’-site-clustering test produced more false-negative results in the presence of introgression (Fig. 5e,f; Supplementary Figs. S21–S24). While fewer false-negative results were found with weak introgression (*m* = 10*^-^*^8^; 247 out of 660), particularly cases with very weak (*m* = 10*^-^*^9^; 261 out of 300), strong (*m* = 10*^-^*^7^; 394 out of 660), and very strong introgression rates (*m* = 10*^-^*^6^; 285 out of 300) did not lead to significant ‘ABBA’-site clustering when the population size was large (*N*_e_ = 10^5^) (Fig. 5e,f; Supplementary Figs. S21–S22). For a lower population size (*N*_e_ = 10^4^) fewer false-negative results were found: While cases with strong (*m* = 10*^-^*^7^; 61 out of 300), very strong (*m* = 10*^-^*^6^; 209 out of 300), and very weak introgression rates (*m* = 10*^-^*^9^; 182 out of 300) produced moderate numbers of false-negative results, only very few (3 out of 300) non-significant results were present with a weak rate of introgression (*m* = 10*^-^*^8^) (Supplementary Figs. S23–S24).

Applying the ‘ABBA’-site-clustering test to the presumably introgression-free empirical dataset for four tribes of Lake Tanganyika cichlids led to the surprising result of highly significant clustering, regardless of whether the “sensitive” or “robust” version of the test were considered and which combinations of species were selected from the four tribes (*p<* 0.0002 in all cases). We investigated these results further by focusing on a randomly selected species quartet, comprising *Tropheus polli* (Tropheini), *Cyprichromis pavo* (Cyprichromini), *Neolamprologus savoryi* (Lamprologini), and *Boulengerochromis microlepis* (Boulengerochromini, placed as outgroup; Ronco et al. 2021). For this species quartet, the “sensitive” and “robust” versions of the ‘ABBA’-site-clustering test strongly supported clustering with *p* = 2.3 *⇥* 10*^-^*^16^ (the smallest value handled by Dsuite) and *p* = 7.8 *⇥* 10*^-^*^9^, respectively. In stark contrast, Dsuite reported a low and non-significant *D*-statistic of *D* = 0.01 for this quartet (with *Cyprichromis pavo* and *Tropheus polli* placed in positions P1 and P2, respectively, based on the number of shared alleles with P3) (Supplementary Table S5).

However, repeating the analysis separately for each of the 23 linkage groups (LG) of the Nile tilapia reference assembly (Conte et al., 2017) revealed that the only linkage group for which both versions of the ‘ABBA’-site-clustering test reported significant clustering was LG2 (*p* = 2.3 *⇥* 10*^-^*^16^ for both test versions), where we also found a high and significant *D*-statistic (*D* = 0.285; *p* = 1.8 *⇥* 10*^-^*^5^). Clustering of “strong ABBA sites” was not detected on any of the other linkage groups with the “robust” version of the test; however, the “sensitive” test version supported clustering on 18 other linkage groups (with 4.7 *⇥* 10*^-^*^6^ 6 *p* 6 0.04), suggesting perhaps an effect of mutation rate variation along the chromosomes. Plotting the positions of “strong ABBA sites” relative to all polymorphic sites (vector *∼g* of the “sensitive” test version) or relative to all “strong ABBA sites” and “strong BABA sites” (vector *^∼^h* of the “robust” test version) clearly illustrates the clustering on LG2 (Fig. 6 for LGs 1–3; Supplementary Figs. S35 and S36 for all LGs).

**Fig. 6.**
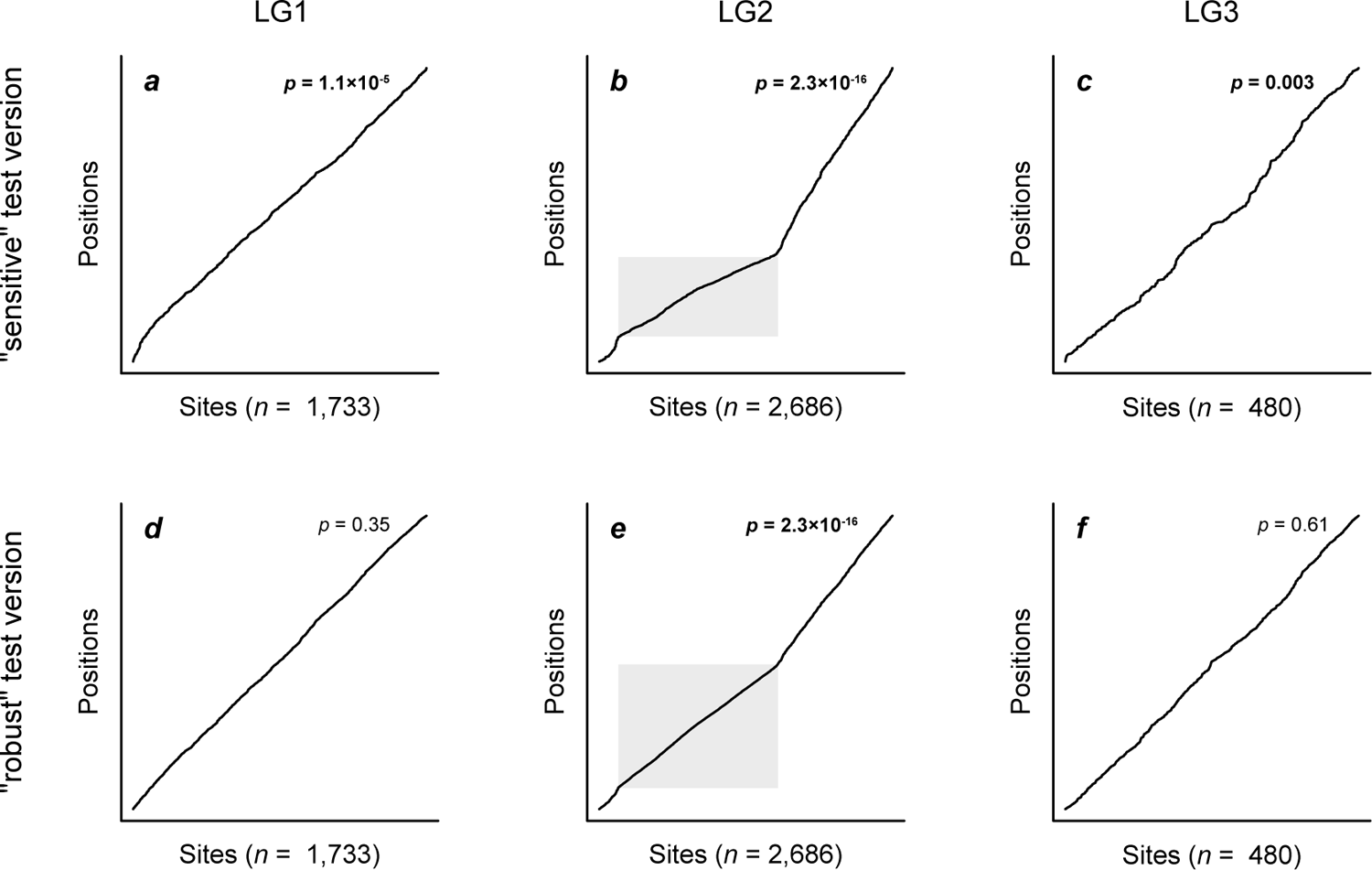
Clustering of ‘ABBA’ sites in empirical data for Lake Tanganyika cichlid fishes. Results are shown for the first three linkage groups (LG) of the Nile tilapia reference assembly; results for all linkage groups are presented in Supplementary Figures S35 and S36. With sorted “strong ABBA sites” on the horizontal axis, the black line indicates their position within a vector of polymorphic sites on the vertical axis. A straight, diagonal line therefore illustrates a homogeneous distribution of these sites within this vector, while changes in the gradient illustrate clustering. significant *p*-values are marked in bold. The gray area indicates a region with increased frequency of “strong ABBA sites” in the first half of LG2.

Repeating this analysis for other quartets of species from the four tribes revealed that the pattern of strong clustering on LG2 was shared by all of them.

## Discussion

As genome-wide data are becoming available for more and more species across the tree of life, these give us the opportunity to investigate the extent of between-species hybridization and introgression in unprecedented detail (Taylor and Larson, 2019). These data have already revealed an unexpected frequency of ongoing or recent introgression, and are beginning to uncover their occurrence also in the deep past (MacGuigan and Near 2019 [Percidae]; Pavón-Vázquez et al. 2021 [Varanidae]; Hodson et al. 2022 [Sciaridae, Cecidomyiidae]). However, the results of studies on ancient introgression must be critically evaluated when they are based on introgression detection methods that were originally developed for recently diverged species or populations (Pease and Hahn, 2015; Hibbins and Hahn, 2022; Zheng and Janke, 2018).

Our results confirm a recent report that demonstrated a sensitivity of Patterson’s *D*-statistic and the related *D*_3_ (Hahn and Hibbins, 2019) and HyDe (Blischak et al., 2018) tests to among-species rate variation (Frankel and Ańe, 2023). We extended these previous results to data simulated with a more diverse range of parameters, including different phylogeny ages, population sizes, mutation rates, and both homogeneous and variable recombination rates, corroborating that the *D*-statistic is generally sensitive to rate variation.

To distinguish between false signals caused by rate variation and genuine indicators of introgression, we propose a test for clustering of ‘ABBA’ sites along the chromosome, a pattern which arises when several polymorphisms are derived from the same introgressed haplotype. Our analyses demonstrated that this test is robust to among-species rate variation, with no false positives after multiple testing correction, and few false negatives across a wide range of datasets. False negatives for the “sensitive” test version were limited mainly to cases of weak introgression in combination with an elevated substitution rate in P2. The “robust” test version, on the other hand, performed most reliably when introgression rates were intermediate, with only a minor or no influence of among-species rate variation. The reason why the “robust” test version returned many false negatives with very strong introgression was that most of the simulated chromosomes carried introgressed sequences in very long continuous blocks. However, such cases of very strong introgression could always be identified reliably by their very high and significant *D*-statistic, along with a highly significant result of the “sensitive” version of the ‘ABBA’-site-clustering test. In their combination, the ‘ABBA’-site-clustering test and *D*-statistics thus form a powerful set of tools to detect introgression across a wide range of settings.

In addition to introgression, clustering of ‘ABBA’ sites could in principle be expected to arise from ILS, as ILS-derived tracts can contain multiple genetic variants. Nevertheless, we showed here that the ‘ABBA’-site clustering test is robust to ILS. In the absence of introgression but presence of ILS, we did not observe any false-positives after multiple-testing correction. This difference in sensitivity to introgression vs. ILS cannot be explained by the lengths of the tracts produced by these two processes, as these were comparable across the simulations. Instead, the explanation likely lies in the difference in numbers of ‘ABBA’ sites within introgressed vs. ILS tracts. Haplotypes introduced through introgression may often have had a long time, at least ten million generations in our simulations, to accumulate the mutations that produce ‘ABBA’ sites following introgression between P3 and P2. On the other hand, haplotypes introduced through ILS had much less time for the accumulation of mutations that would ultimately produce ‘ABBA’ or ‘BABA’ sites – on average, one coalescent unit (2*N*_e_ generations) – in the common ancestor of P1, P2, and P3. Thus, the tracts produced by ILS carry fewer ‘ABBA’ sites than those produced by introgression, which, as a consequence, renders the ‘ABBA’-site-clustering test robust to ILS.

Besides introgression and ILS, mutation-rate variation along the chromosome, for example driven by an elevated rate in GC-rich mutation hotspots (Śegurel et al., 2014; Nesta et al., 2021), can also cause clustering of ‘ABBA’ sites. Both versions of the ‘ABBA’-site-clustering test are designed to account for this variation to some degree. In the “sensitive” test version, clustering of “strong ABBA sites” is considered relative to all polymorphic sites, while the “robust” version of the test assesses clustering relative to “strong BABA sites”. All of these increase in frequency along with ‘ABBA’ sites in mutation hotspots. However, our simulations revealed that, at least for some parameter combinations, the frequency of ‘ABBA’-pattern homoplasies among all polymorphic sites is higher in mutation-rate hotspots, leading to their clustering and some false positives for the “sensitive” test. However, the relative probabilities of ‘ABBA’ and ‘BABA’ homoplasies both scale equally with the local mutation rate. This is why the version of the test that focuses only on these two types of sites is robust to variation in the mutation rate along the chromosome.

The application of our ‘ABBA’-site-clustering test to a presumably introgression-free empirical dataset led to the surprise identification of a single linkage group – LG2 – on which not just our test produced a strong signal of introgression, but where this signal was also confirmed by a high and clearly significant *D*-statistic. For other linkage groups, in contrast, significant clustering was detected only with the “sensitive” version, but not the “robust” version of the ‘ABBA’-site-clustering test, suggesting that this clustering is in fact derived from mutation-rate variation and not from introgression.

For LG2, the signal detected by our test as well as the *D*-statistic stemmed from a high frequency of ‘ABBA’ sites on the first half of the linkage group. Due to the clear localization of the signal to a specific region of the chromosome and its consistency across many different species quartets, we suspect that it may be contained within a large region of low (or no) recombination, possibly facilitated by a chromosomal inversion. Two scenarios could then explain the localized clustering of ‘ABBA’ sites: The region could have been transferred between species due to introgression that otherwise left little to no signal in the genome, or it could result from ILS. Further comparative analyses of species quartets could help to discriminate between these two options, and might reveal interesting insights into the evolution of Lake Tanganyika cichlids in a future study. Here, however, we limit our conclusion for this analysis to the performance of the ‘ABBA’-site-clustering test: First, we conclude that the “robust” version of the test did not produce any false positives. And second, we note that the test is able to identify large regions, that potentially derived from introgression, even more clearly than the *D*-statistic.

Our implementation of the ‘ABBA’-site-clustering test in the program Dsuite is easy to use and comes with negligible added cost to Dsuite analyses. Given that Dsuite is among the fastest tools available for the calculation of *D*-statistics (Malinsky et al., 2021), the additional application of the ‘ABBA’-site clustering test should be computationally feasible for all users.

Our analyses of simulated datasets revealed that tree-based methods can be useful for the detection of introgression when rate variation is present, and identified the conditions under which each approach performs reliably. While we observed an effect of long-branch attraction affecting the *D*_tree_ statistic, this effect was weak and only noticeable when all results were considered in aggregate. In fact, none of the datasets simulated without introgression produced a false-positive, significant *D*_tree_ statistic, even when *D*_tree_ itself reached the maximum value of 1.0 (Fig. 3a). On the other hand, *D*_tree_ was consistently large and significant even with weak introgression (*m* = 10*^-^*^8^; Fig. 3b,e,h), suggesting that *D*_tree_ is a powerful detector of introgression.

Besides *D*_tree_, SNaQ appeared to be robust to rate variation across our simulated datasets (Fig. 3j,p). Given that SNaQ analyzes tree topologies, however, we caution that the same weak bias that affected *D*_tree_ might also be relevant for SNaQ. Like for *D*_tree_, we thus advise that weaker signals reported by SNaQ might better be ignored. Furthermore, it has been pointed out that the use of AIC is inappropriate for the comparison of SNaQ results, due to the pseudolikelihood framework employed by SNaQ (Hibbins and Hahn, 2022). To avoid this issue, users of SNaQ may want to focus – like we did – on sets of four taxa with putative introgression, in which case SNaQ calculates and reports actual likelihood values (Soĺıs-Lemus and Ańe, 2016).

Finally, we found that the performance of QuIBL depended strongly on the length of the alignments used to generate the input tree set. Given that QuIBL produced many false-positive signals of introgression regardless of rate variation when the alignments were short (Supplementary Fig. S11), the use of longer alignments, with lengths of at least 1,000 bp may be recommendable. With such alignments as input, QuIBL performed rather reliably (Supplementary Fig. S13) and detected most cases of stronger introgression.

In practice, the inference of ancient introgression between divergent species may often be hampered by the requirements of detection methods. Site-pattern-based methods (such as the *D*-statistic and the ‘ABBA’-site clustering test) require SNP datasets that are typically obtained through read mapping towards a reference genome assembly. When investigating divergent taxa, however, it may no longer be possible to map all of them reliably to the same reference genome. As a result, SNP datasets produced for such taxa may be limited and prone to reference bias particularly in taxa with lower read coverage, which can generate misleading signals of introgression (Günther and Nettelblad, 2019). To minimize the chance of reference bias while also reducing the numbers of homoplasies, outgroup species should be chosen that are as closely related to the ingroup as possible. As an alternative that does not depend on a reference, multi-marker sets of alignments have traditionally been produced through ortholog-identification approaches focusing on genes or ultra-conserved elements. While these approaches may be more suited for divergent taxa than read mapping, they are generally limited to certain regions of the genome, corresponding to a set of input query sequences.

Fortunately, two recent developments promise to overcome these limitations, rendering larger regions of the genome accessible for the detection of ancient introgression: First, methods for whole-genome alignment have finally matured to the degree that they can be applied to hundreds of genome assemblies of highly diverged taxa (Armstrong et al., 2020). By using assemblies instead of mapped reads, these whole-genome alignments are immune to reference bias, and allow the extraction of massive numbers of SNPs for site-pattern-based methods, or of alignment blocks for tree-based methods. Second, more and more genome assemblies are now highly contiguous, chromosome-resolved or nearly so (Rhie et al., 2021; Formenti et al., 2022). This is relevant for the completeness of whole-genome alignments, and reduces their fragmentation. Both will contribute to the utility of the new ‘ABBA’-site clustering test, given that this test requires contiguous genomic blocks within which clustering can be observed.

In their combination, these new developments are now allowing us to push the limits of reliable introgression detection, enabling the inference of introgression even among species that have diverged many tens of millions of years ago. We are thus coming closer to being able to assess the true extent of hybridization and introgression across the tree of life.

## Code Availability

Code for all our computational analyses is available on https://github.com/thorekop/ABBA-Site-Clustering

## Acknowledgements

We are grateful to Fabrizia Ronco and Kjetil Lysne Voje for valuable help and support with R scripts. We thank Miriam Miyagi and Nathaniel Edelman for advice on how to interpret the QuIBL output when using trees with polytomies. All computations were performed on resources provided by Sigma2 – the National Infrastructure for High Performance Computing and Data Storage in Norway.

## Supplementary Material

Data available from the Dryad Digital Repository: http://dx.doi.org/10.5061/dryad.sf7m0cgbs.

