## Supplementary Material for "Towards reliable detection of introgression in the presence of among-species rate variation"

**Koppetsch et al.**

### Supplementary Notes

**Supplementary Note S1:** The decrease in the $D$-statistic with increasing age of the phylogeny (i.e. with $t_{\mathrm{P1,P2}}$) was caused by homoplasies and reversals accumulating on the longer branches of the older phylogenies, augmenting both $C_{\mathrm{ABBA}}$ and $C_{\mathrm{BABA}}$. Using the results with strong introgression shown in Figure 2f as an example, $C_{\mathrm{ABBA}}$ and $C_{\mathrm{BABA}}$ were in the ranges 93,608–101,459 and 16,458–16,961, respectively, for the youngest phylogeny ($t_{\mathrm{P1,P2}} = 10$ million generation), producing a $D$-statistic between 0.70 and 0.72. For the oldest phylogeny ($t_{\mathrm{P1,P2}} = 30$ million generation), however, $C_{\mathrm{ABBA}}$ and $C_{\mathrm{BABA}}$ were around 131,143–139,571 and 65,467–66,364, respectively, and the corresponding $D$-values were between 0.33 and 0.36. Thus, both $C_{\mathrm{ABBA}}$ and $C_{\mathrm{BABA}}$ increased by a number on the order of 40,000–50,000. More detailed analysis of the output of one randomly selected simulation replicate confirmed that this was due to homoplasies and reversals. Counting only bi-allelic sites, the older phylogeny had 12,922 homoplasies more between P1 and P3 than the younger phylogeny, increasing $C_{\mathrm{BABA}}$. A greater effect, however, was produced by homoplasies occurring between P2 and P4, which also contribute to $C_{\mathrm{BABA}}$: Of these, the older phylogeny had 15,677 more than the younger phylogeny. The older phylogeny also had 3,372 more reversals in P2 that further increased $C_{\mathrm{BABA}}$. In addition, the older phylogeny had 43,377 more multi-allelic sites that likely explain the remainder of the increase in $C_{\mathrm{BABA}}$ (Table 1).

**Supplementary Note S2:** Simulations with among-site recombination-rate variation.
To assess whether recombination-rate variation could affect the reliability of our 'ABBA'-site-clustering test, we repeated a number of simulations with a recombination map instead of a homogeneous recombination rate. As a well-resolved recombination map for a chromosome with a relatively small size, we selected the HapMap Phase II genetic map (Genome Reference Consortium Human Build 37) for human chromosome 20 (The International HapMap Consortium, 2007). We truncated its length from 62.9 Mbp to 20 Mbp to match that of the chromosomes produced in all other simulations. The simulations with this recombination map were restricted to the larger population size ($N_{\mathrm{e}} = 10^5$), the faster mutation rate ($\mu = 2 \times 10^{-9}$), and three introgression rates ($m \in \{0, 10^{-8}, 10^{-7}\}$). These use of a variable recombination rate, however, did not influence the performance of the 'ABBA'-site-clustering test (Supplementary Figs. S25 and S26).

**Supplementary Note S3:** Simulations with increased ILS.
We further assessed the performance of both the "sensitive" and the "robust" version of the 'ABBA'-site-clustering test by repeating a subset of the simulations with higher levels of ILS. The degree of ILS was increased by shortening the internal branch corresponding to the common ancestor of P1 and P2, from an original length of 10 million generations to reduced lengths of 1 million generations, 100,000 generations, and 10,000 generations. We call these alternative scenarios "moderate ILS", "high ILS", and "very high ILS". The terminal branches leading to species P3 and P4 were shortened accordingly, and as a result, the most recent common ancestor of the trio P1, P2, and P3, as well as the root node became younger by the same amount. Focusing on potential false positive signals of introgression, we performed these simulations without introgression ($m = 0$) and also excluded

rate variation ($s = 1$), testing the three divergence times ($t_{\mathrm{P1,P2}} \in \{1 \times 10^7, 2 \times 10^7, 3 \times 10^7\}$), both population sizes ($N_\mathrm{e} \in \{10^4, 10^5\}$), and the two mutation rates ($\mu \in \{1 \times 10^{-9}, 2 \times 10^{-9}\}$), with ten replicate datasets for each parameter combination.

In the "moderate ILS" setting, between 42,174 and 47,652 bp (0.21 – 0.24%) of the 20 Mbp-chromosomes simulated with a population size $N_\mathrm{e} = 10^5$ had a topology different from the species tree as the result of ILS. With the "high ILS" setting, the simulations with $N_\mathrm{e} = 10^4$ also led to ILS, affecting between 39,191 and 62,778 bp (0.20 – 0.31%) of the 20-Mbp chromosomes. With $N_\mathrm{e} = 10^5$, as much as $3,972,624 - 4,058,703$ bp (19.9 – 20.3%) carried ILS. The mean lengths of ILS-derived tracts ranged between 71.8 and 837.0 bp. These tract lengths are comparable for example to those introduced by weak introgression in combination with the larger population size ($m = 10^{-8}$, $N_\mathrm{e} = 10^5$), which ranged from 476.8 to 502.4 bp and led to highly significant results with the "sensitive" version of the 'ABBA'-site clustering test (Fig. 4b).

For the "very high ILS" setting, the simulations with $N_\mathrm{e} = 10^4$ also led to ILS, affecting between 7,818,986 and 8,282,532 bp (39.1 – 41.4%) of the 20-Mbp chromosomes. With $N_\mathrm{e} = 10^5$, as much as $12,593,496 - 12,698,566$ bp (63.0 – 63.5%) carried ILS. The mean lengths of ILS-derived tracts ranged from 270.4 to 2184.2 bp. Crucially, these substantial levels of ILS did not lead to false-positive signals of introgression both in the "sensitive" and the "robust" version of the 'ABBA'-site clustering test. For the "sensitive" version of the 'ABBA'-site clustering test, only three out of the 120 simulated datasets under the "moderate ILS" scenario produced significant $p$-values and none remained significant after Bonferroni correction. With the "high ILS" scenario, seven tests were just about statistically significant but, again, none remained significant after Bonferroni correction. With the "robust" version of the 'ABBA'-site clustering test, no significant $p$-values could be found for both the "moderate ILS" and "high ILS" scenario. The "high ILS" scenario produced a single significant $p$-value, being also not significant after Bonferroni correction.

**Supplementary Note S4:** Simulations with among-site mutation-rate variation.
Finally, we assess the robustness of both versions of the 'ABBA'-site-clustering test to mutation-rate variation across the genome, by repeating a number of simulations with a mutation map instead of applying the same mutation rate to all sites of the simulated chromosome. These simulations were restricted to the larger population size ($N_\mathrm{e} = 10^5$), the faster mutation rate ($\mu = 2 \times 10^{-9}$), and three introgression rates ($m \in \{0, 10^{-8}, 10^{-7}\}$). We generated the mutation map by drawing, for each window of 500 bp on the simulated 20-Mbp chromosome, a rate at random from an exponential distribution with a mean set to $\overline{\mu} = 2 \times 10^{-9}$ per site per generation. As alternatives, we also used gamma-distributed rate variation (with a shape parameter of 2 and a scale parameter of $10^{-9}$; thus again $\overline{\mu} = 2 \times 10^{-9}$) or a larger window size of 1,000 bp; however, as these alternative settings led to very similar results of the 'ABBA'-site-clustering test, we here focus on those results obtained with the exponential distribution and a window size of 500 bp. These results, shown in Supplementary Figures S33 and S34, suggest that the "sensitive" version of the 'ABBA'-site-clustering test produces false signals of introgression when the mutation rate varies across the genome (Supplementary Fig. S33). The "robust" version of the test, however, produces no such false signals.

### Supplementary Figures

**Supplementary Figure S1:** Signals of introgression detected with the $D$-statistic for datasets simulated with a population size $N_e = 10^5$, a mutation rate $\mu = 1 \times 10^{-9}$, an introgression rate $m \in \{0, 10^{-9}, 10^{-8}, 10^{-7}, 10^{-6}\}$, a P2 branch rate $s \in \{0.25, 0.5, 1, 2, 4\}$, and a divergence time $t_{\mathrm{P1,P2}} \in \{1 \times 10^7, 2 \times 10^7, 3 \times 10^7\}$. Note that for combinations of increased P2 branch rates and low or no introgression (either $s = 2$ and $m \leq 10^{-9}$ or $s = 4$ and $m \leq 10^{-8}$), P3 always shared more alleles with P1, not P2 (the $D$-statistic is nevertheless positive because Dsuite in this case exchanges the two taxa in its output).

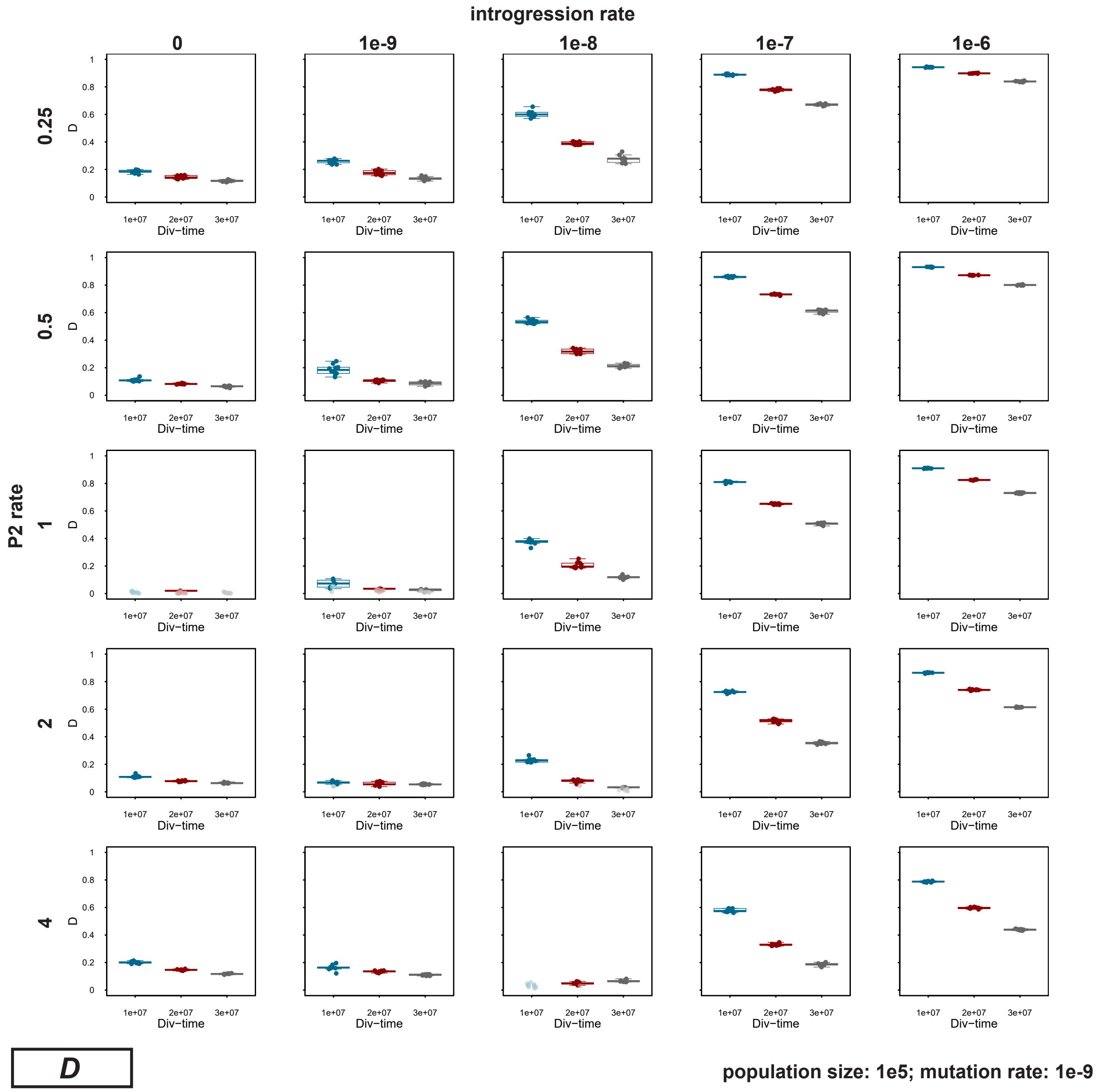

**Supplementary Figure S2:** Signals of introgression detected with the $D$-statistic for datasets simulated with a population size $N_e = 10^5$, a mutation rate $\mu = 2 \times 10^{-9}$, an introgression rate $m \in \{0, 10^{-9}, 10^{-8}, 10^{-7}, 10^{-6}\}$, a P2 branch rate $s \in \{0.25, 0.5, 1, 2, 4\}$, and a divergence time $t_{\mathrm{P1,P2}} \in \{1 \times 10^7, 2 \times 10^7, 3 \times 10^7\}$. For introgression rate $m \in \{0, 10^{-8}, 10^{-7}\}$ and P2 branch rate $s \in \{0.25, 1, 4\}$, 50 replicates were simulated; for all other parameter combinations, we performed ten replicate simulations.

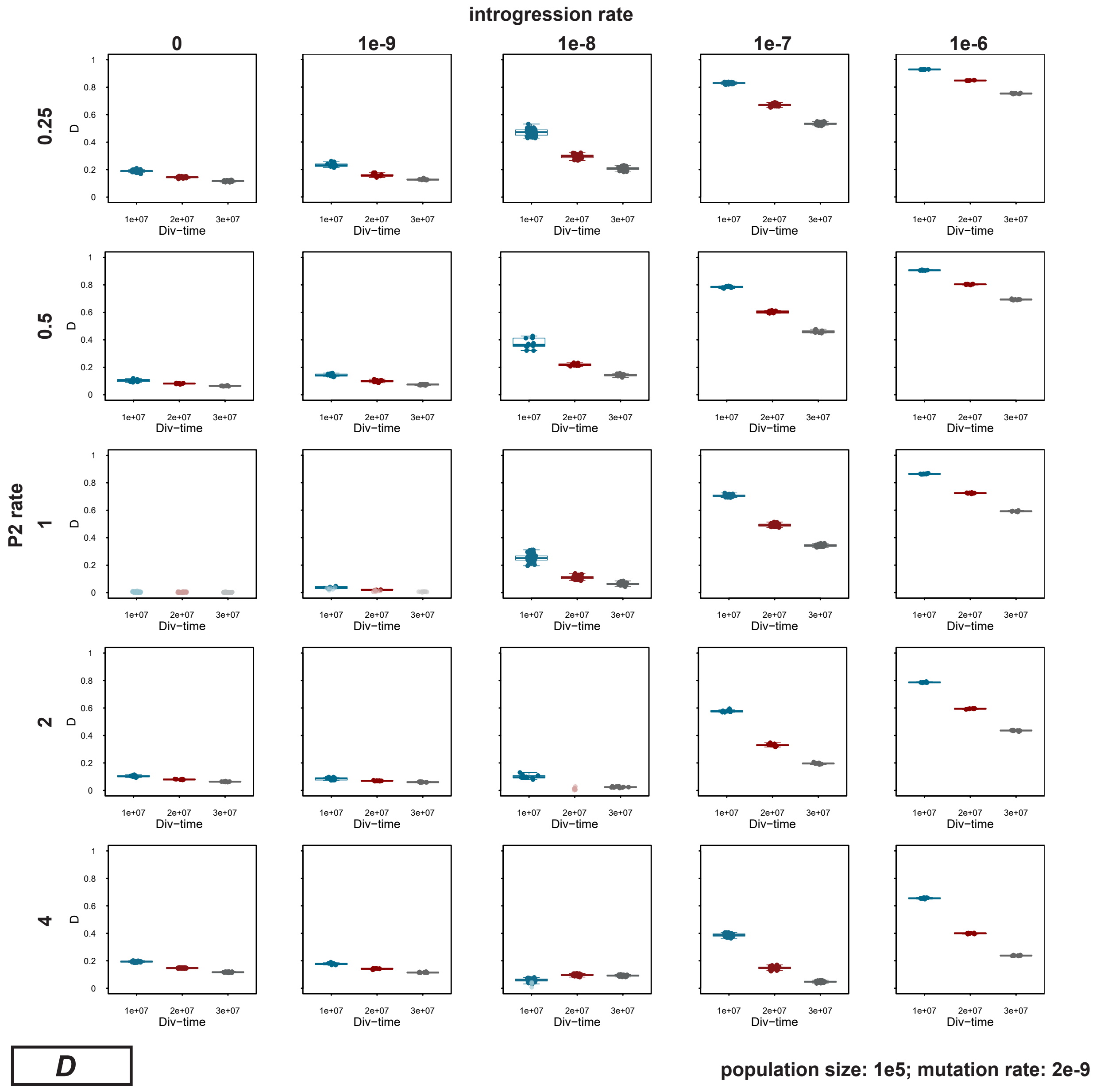

**Supplementary Figure S3:** Signals of introgression detected with the $D$-statistic for datasets simulated with a population size $N_e = 10^4$, a mutation rate $\mu = 1 \times 10^{-9}$, an introgression rate $m \in \{0, 10^{-9}, 10^{-8}, 10^{-7}, 10^{-6}\}$, a P2 branch rate $s \in \{0.25, 0.5, 1, 2, 4\}$, and a divergence time $t_{\mathrm{P1,P2}} \in \{1 \times 10^7, 2 \times 10^7, 3 \times 10^7\}$.

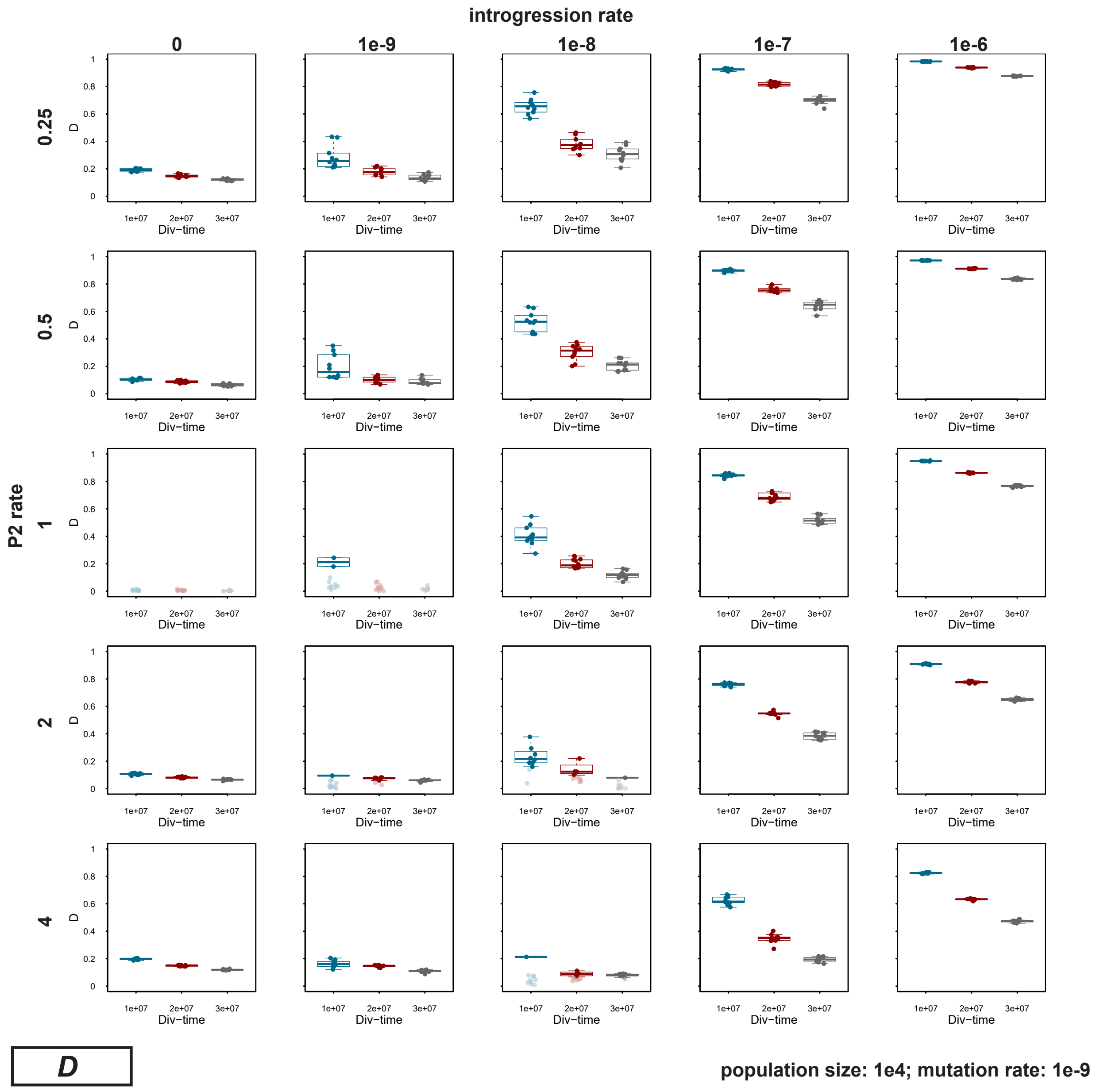

**Supplementary Figure S4:** Signals of introgression detected with the $D$-statistic for datasets simulated with a population size $N_\mathrm{e} = 10^4$, a mutation rate $\mu = 2 \times 10^{-9}$, an introgression rate $m \in \{0, 10^{-9}, 10^{-8}, 10^{-7}, 10^{-6}\}$, a P2 branch rate $s \in \{0.25, 0.5, 1, 2, 4\}$, and a divergence time $t_{\mathrm{P1,P2}} \in \{1 \times 10^7, 2 \times 10^7, 3 \times 10^7\}$.

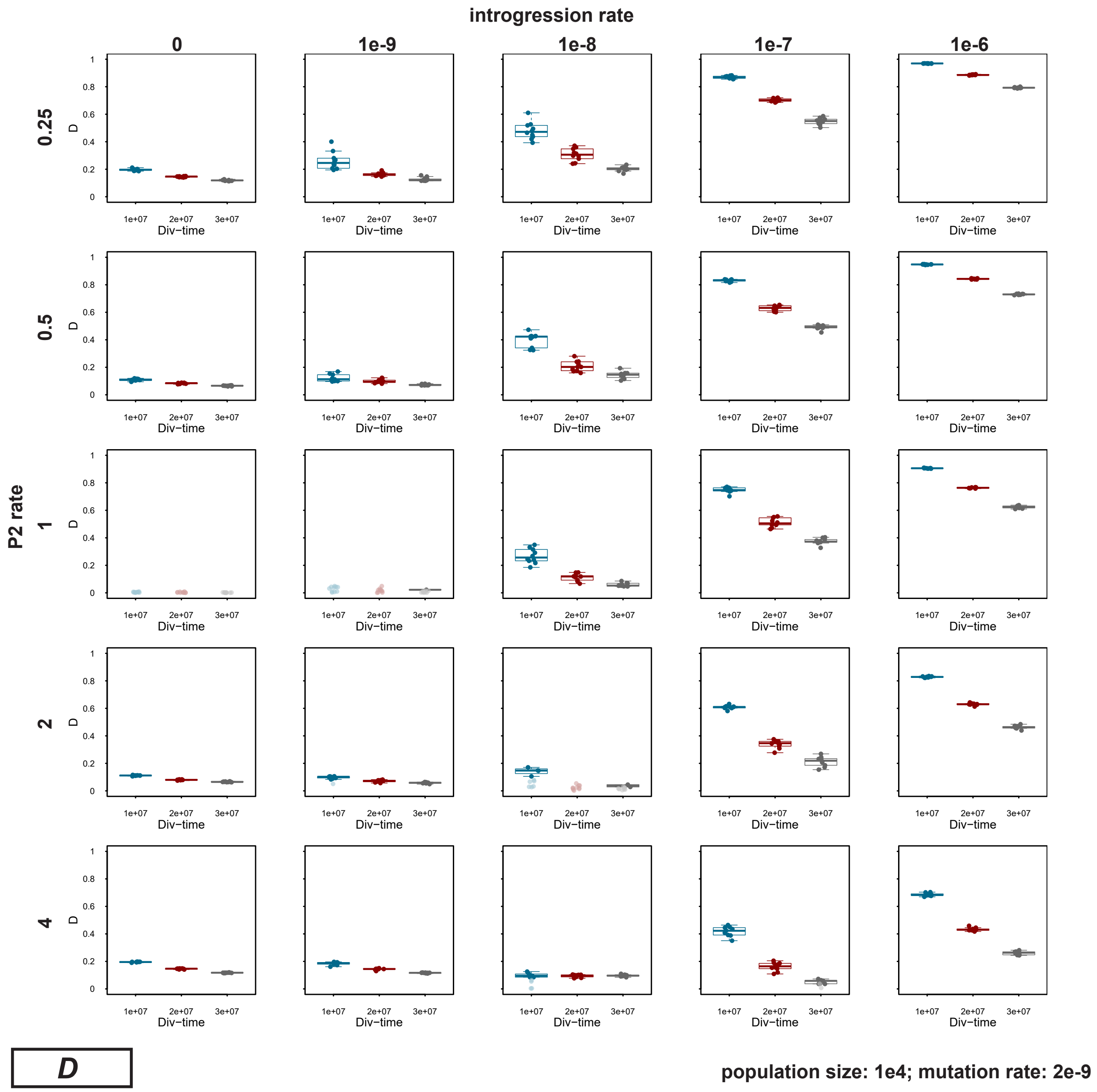

**Supplementary Figure S5:** Signals of introgression detected with the $D_{\text{tree}}$ statistic for datasets simulated with a population size $N_{\text{e}} = 10^5$, a mutation rate $\mu = 2 \times 10^{-9}$, an introgression rate $m \in \{0, 10^{-9}, 10^{-8}, 10^{-7}, 10^{-6}\}$, a P2 branch rate $s \in \{0.25, 0.5, 1, 2, 4\}$, a divergence time $t_{\text{P1,P2}} \in \{1 \times 10^7, 2 \times 10^7, 3 \times 10^7\}$, and an alignment length of 200 bp.

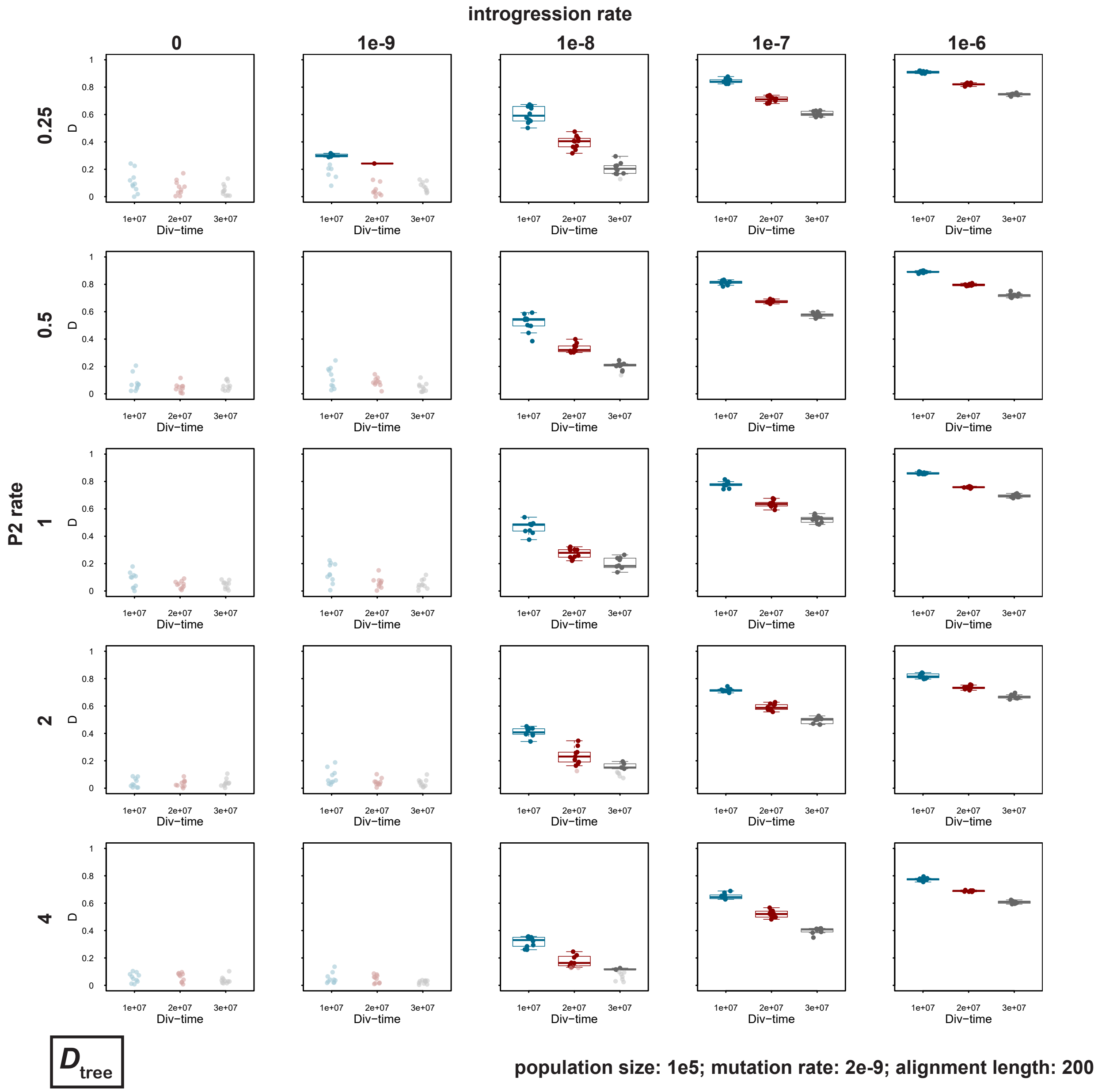

**Supplementary Figure S6:** Signals of introgression detected with the $D_{\text{tree}}$ statistic for datasets simulated with a population size $N_e = 10^5$, a mutation rate $\mu = 2 \times 10^{-9}$, an introgression rate $m \in \{0, 10^{-9}, 10^{-8}, 10^{-7}, 10^{-6}\}$, a P2 branch rate $s \in \{0.25, 0.5, 1, 2, 4\}$, a divergence time $t_{\text{P1,P2}} \in \{1 \times 10^7, 2 \times 10^7, 3 \times 10^7\}$, and an alignment length of 500 bp. For introgression rate $m \in \{0, 10^{-8}, 10^{-7}\}$ and P2 branch rate $s \in \{0.25, 1, 4\}$, 50 replicates were simulated; for all other parameter combinations, we performed ten replicate simulations.

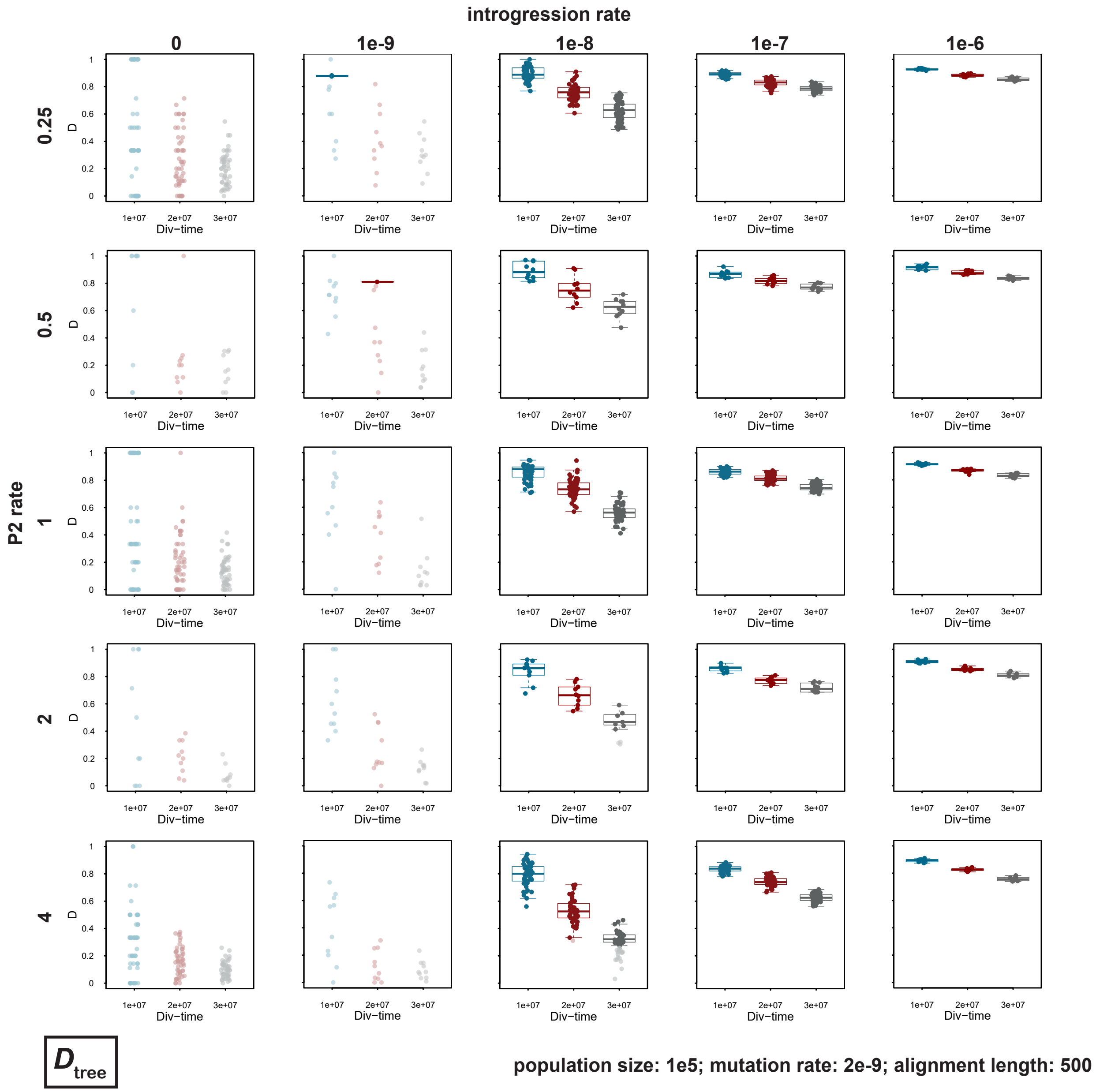

**Supplementary Figure S7:** Signals of introgression detected with the $D_{\text{tree}}$ statistic for datasets simulated with a population size $N_e = 10^5$, a mutation rate $\mu = 2 \times 10^{-9}$, an introgression rate $m \in \{0, 10^{-9}, 10^{-8}, 10^{-7}, 10^{-6}\}$, a P2 branch rate $s \in \{0.25, 0.5, 1, 2, 4\}$, a divergence time $t_{\text{P1,P2}} \in \{1 \times 10^7, 2 \times 10^7, 3 \times 10^7\}$, and an alignment length of 1,000 bp.

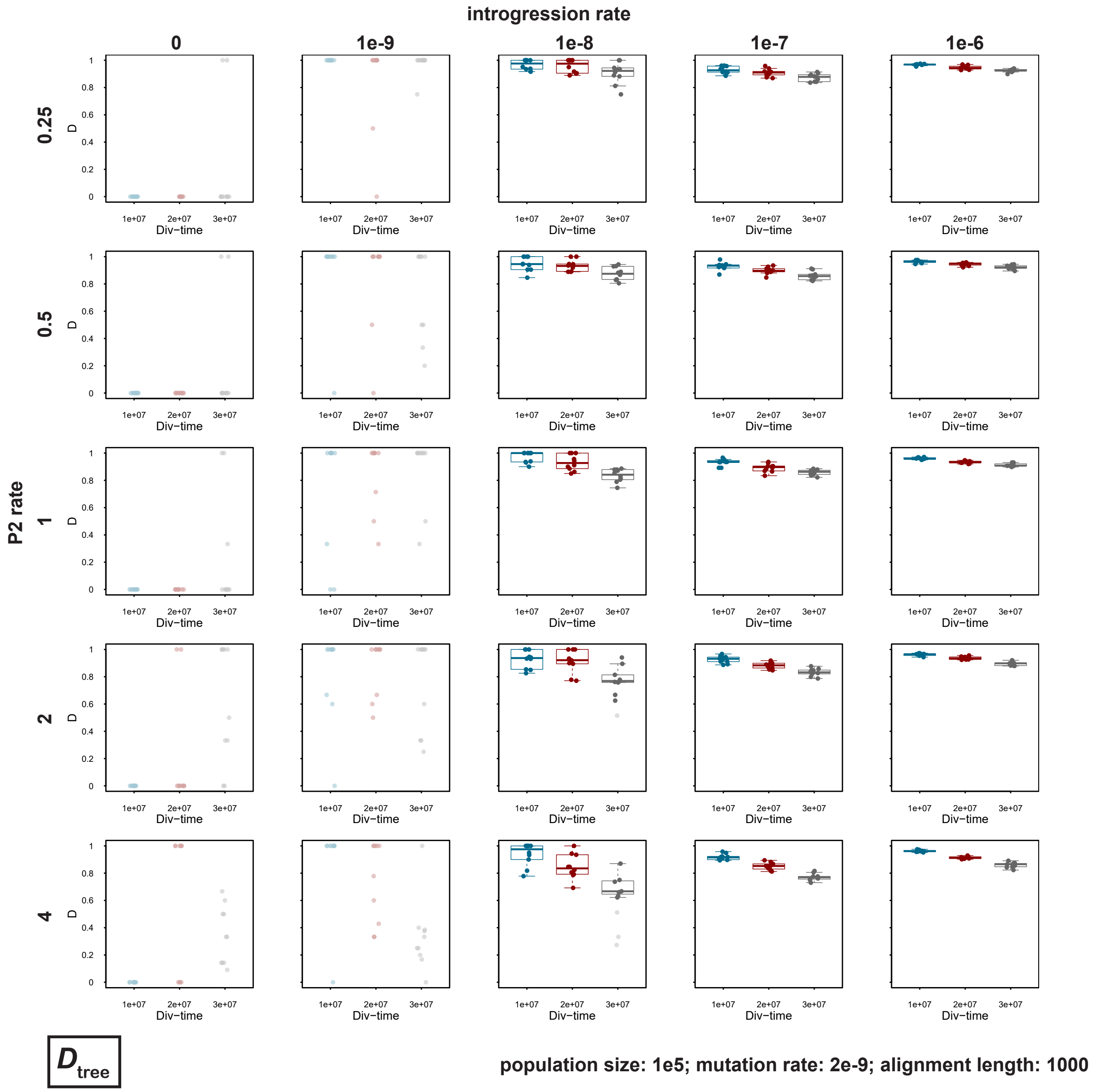

**Supplementary Figure S8:** Signals of introgression detected with SNaQ for datasets simulated with a population size $N_\mathrm{e} = 10^5$, a mutation rate $\mu = 2 \times 10^{-9}$, an introgression rate $m \in \{0, 10^{-9}, 10^{-8}, 10^{-7}, 10^{-6}\}$, a P2 branch rate $s \in \{0.25, 0.5, 1, 2, 4\}$, a divergence time $t_{\mathrm{P1,P2}} \in \{1 \times 10^7, 2 \times 10^7, 3 \times 10^7\}$, and an alignment length of 200 bp. Gray crosshairs indicate results based on incorrectly inferred topologies.

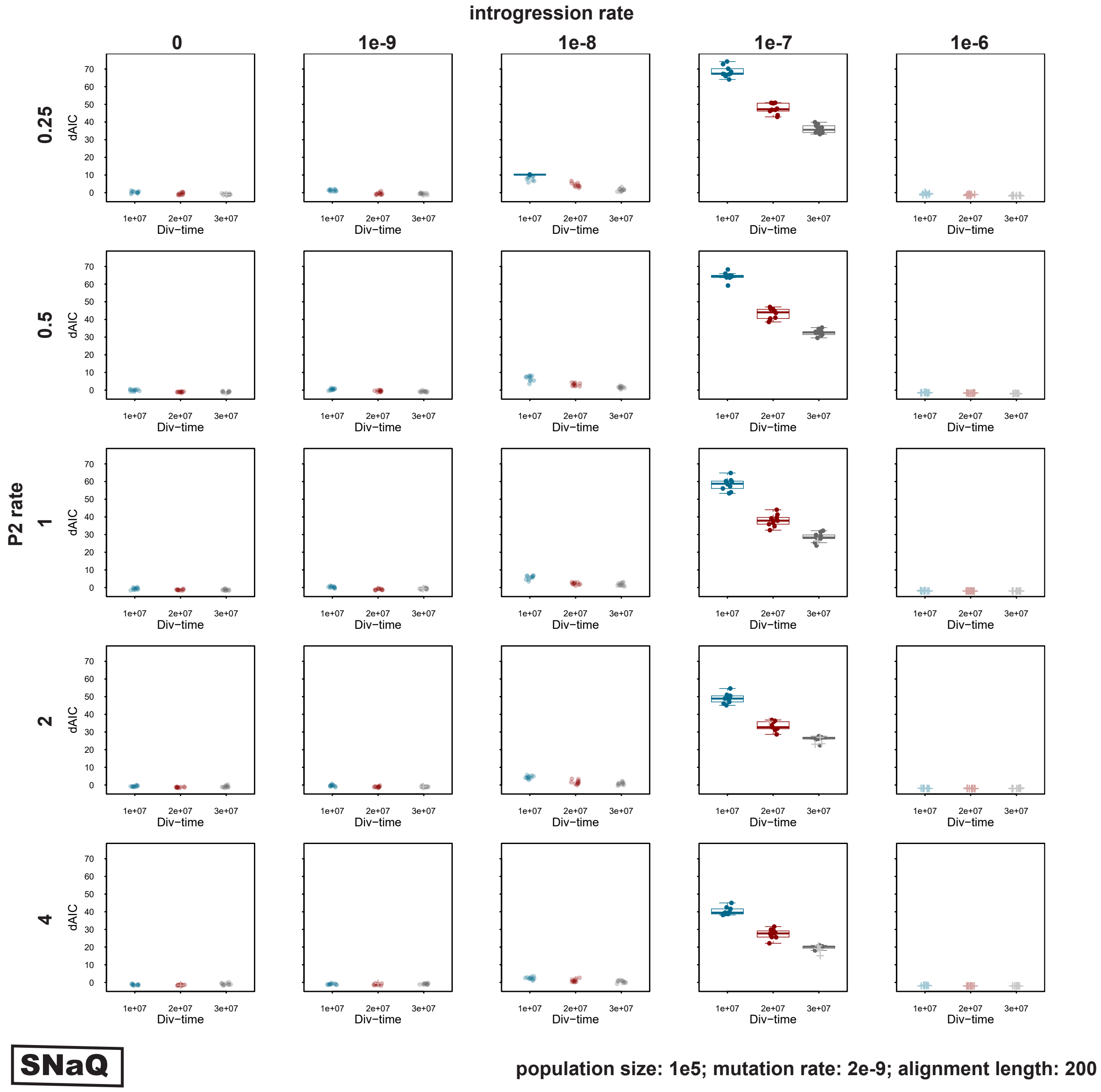

**Supplementary Figure S9:** Signals of introgression detected with SNaQ for datasets simulated with a population size $N_\mathrm{e} = 10^5$, a mutation rate $\mu = 2 \times 10^{-9}$, an introgression rate $m \in \{0, 10^{-9}, 10^{-8}, 10^{-7}, 10^{-6}\}$, a P2 branch rate $s \in \{0.25, 0.5, 1, 2, 4\}$, a divergence time $t_{\mathrm{P1,P2}} \in \{1 \times 10^7, 2 \times 10^7, 3 \times 10^7\}$, and an alignment length of 500 bp. Gray crosshairs indicate results based on incorrectly inferred topologies. For introgression rate $m \in \{0, 10^{-8}, 10^{-7}\}$ and a P2 branch rate $s \in \{0.25, 1, 4\}$, 50 replicates were simulated; for all other parameter combinations, we performed ten replicate simulations.

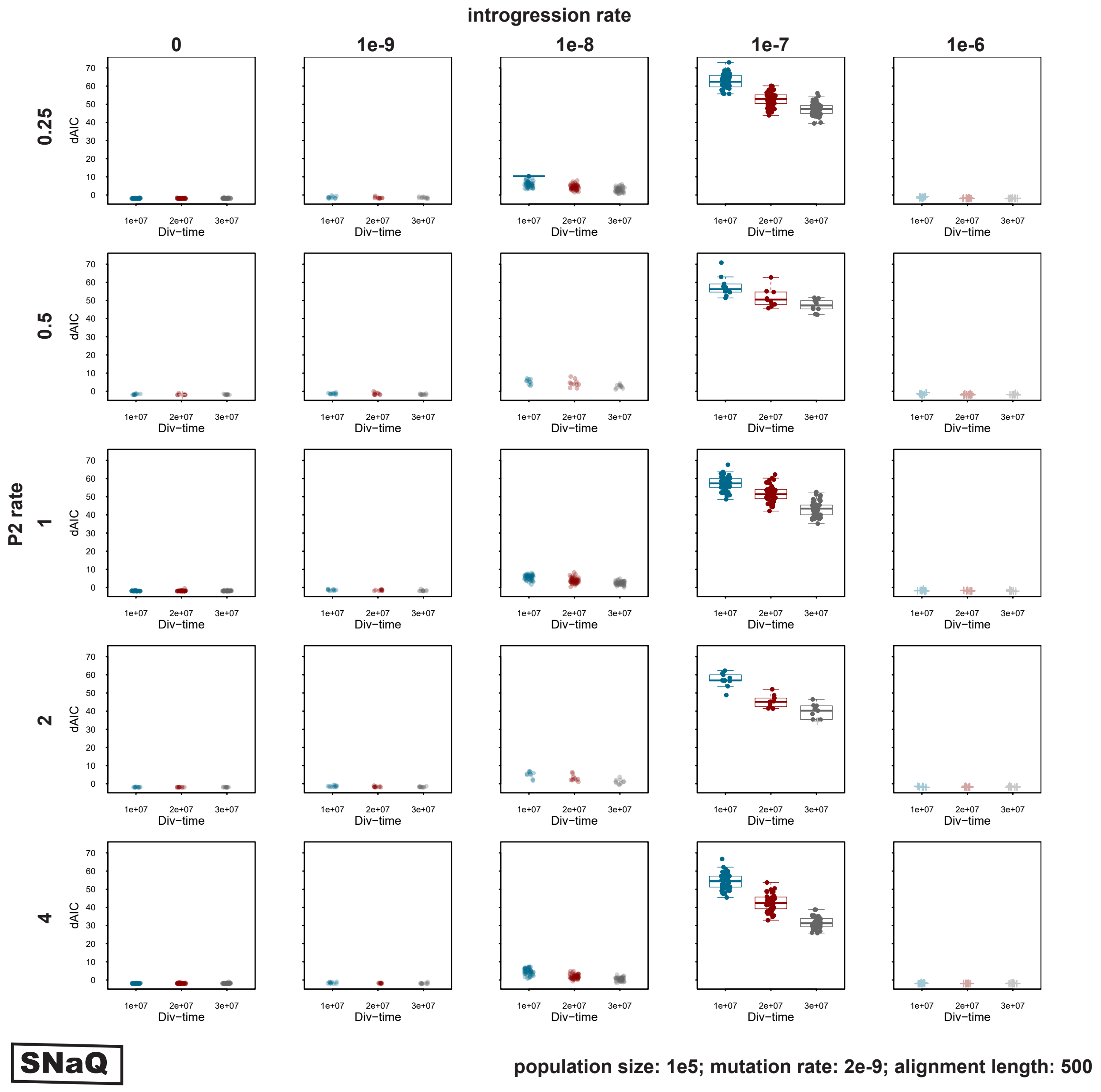

**Supplementary Figure S10:** Signals of introgression detected with SNaQ for datasets simulated with a population size $N_e = 10^5$, a mutation rate $\mu = 2 \times 10^{-9}$, an introgression rate $m \in \{0, 10^{-9}, 10^{-8}, 10^{-7}, 10^{-6}\}$, a P2 branch rate $s \in \{0.25, 0.5, 1, 2, 4\}$, a divergence time $t_{\mathrm{P1,P2}} \in \{1 \times 10^7, 2 \times 10^7, 3 \times 10^7\}$, and an alignment length of 1,000 bp. Gray crosshairs indicate results based on incorrectly inferred topologies.

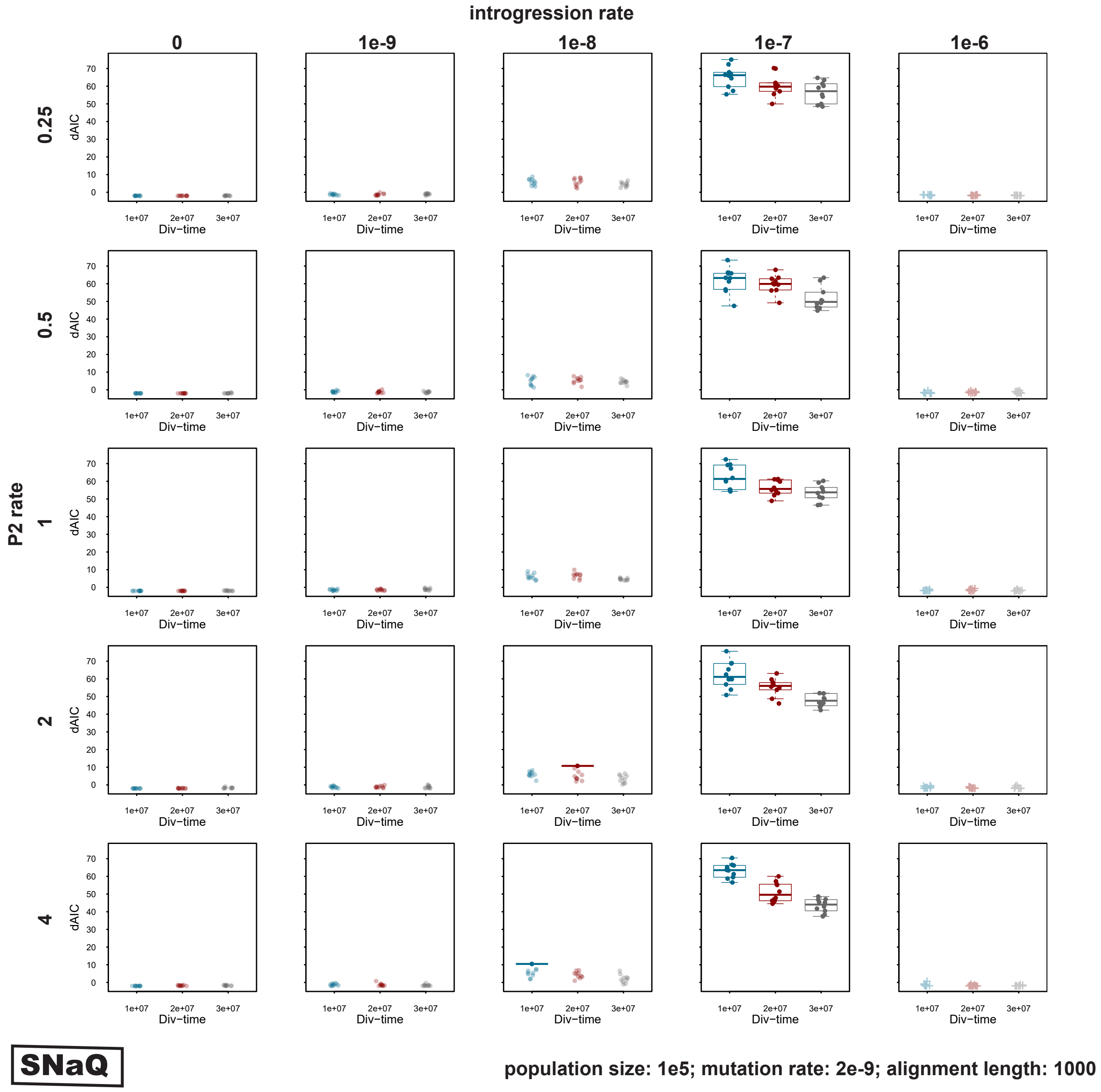

**Supplementary Figure S11:** Signals of introgression detected with QuIBL for datasets simulated with a population size $N_e = 10^5$, a mutation rate $\mu = 2 \times 10^{-9}$, an introgression rate $m \in \{0, 10^{-9}, 10^{-8}, 10^{-7}, 10^{-6}\}$, a P2 branch rate $s \in \{0.25, 0.5, 1, 2, 4\}$, a divergence time $t_{\mathrm{P1,P2}} \in \{1 \times 10^7, 2 \times 10^7, 3 \times 10^7\}$, and an alignment length of 200 bp.

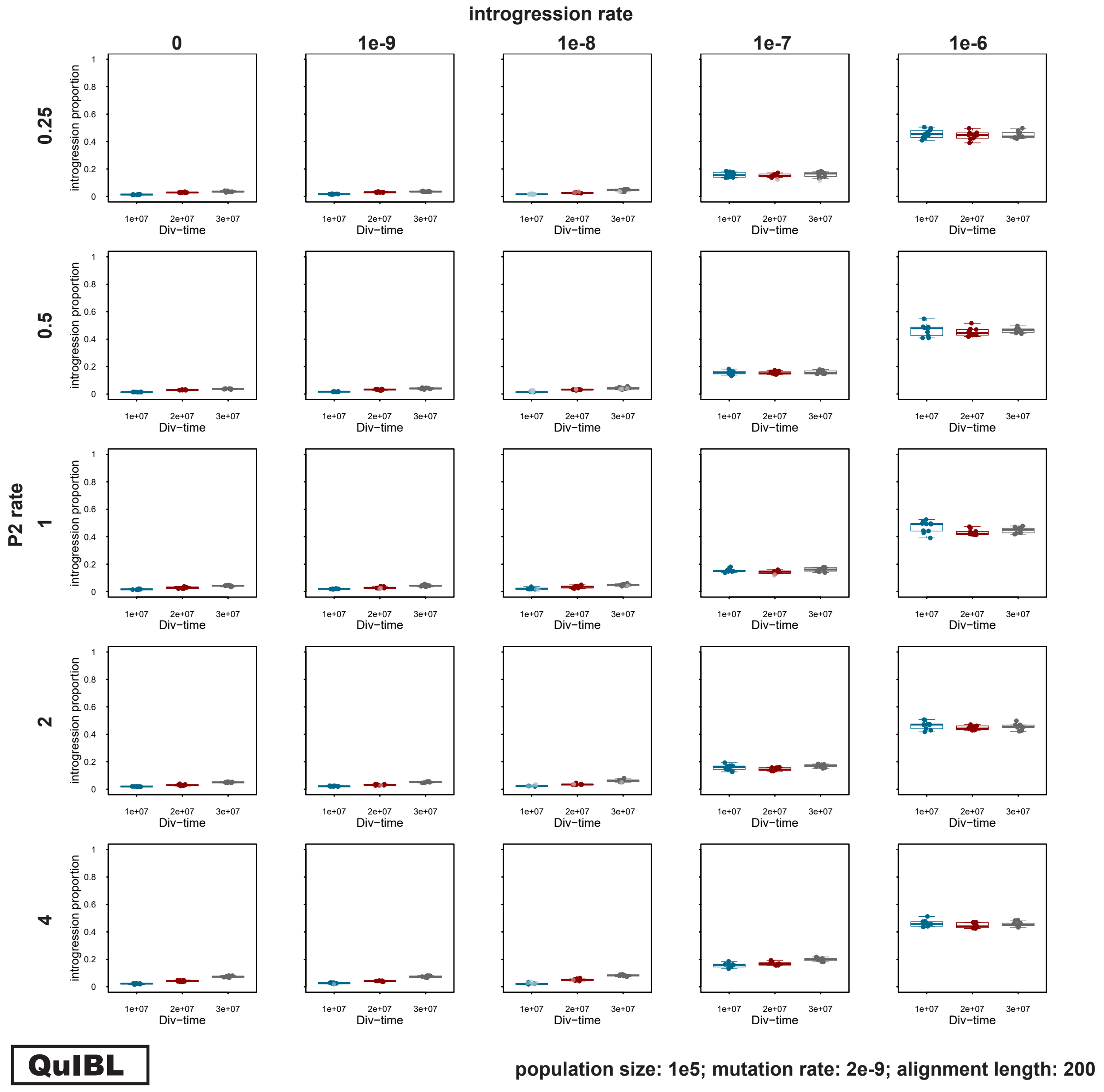

**Supplementary Figure S12:** Signals of introgression detected with QuIBL for datasets simulated with a population size $N_e = 10^5$, a mutation rate $\mu = 2 \times 10^{-9}$, an introgression rate $m \in \{0, 10^{-9}, 10^{-8}, 10^{-7}, 10^{-6}\}$, a P2 branch rate $s \in \{0.25, 0.5, 1, 2, 4\}$, a divergence time $t_{\text{P1,P2}} \in \{1 \times 10^7, 2 \times 10^7, 3 \times 10^7\}$, and an alignment length of 500 bp. For introgression rate $m \in \{0, 10^{-8}, 10^{-7}\}$ and a P2 branch rate $s \in \{0.25, 1, 4\}$, 50 replicates were simulated; for all other parameter combinations, we performed ten replicate simulations.

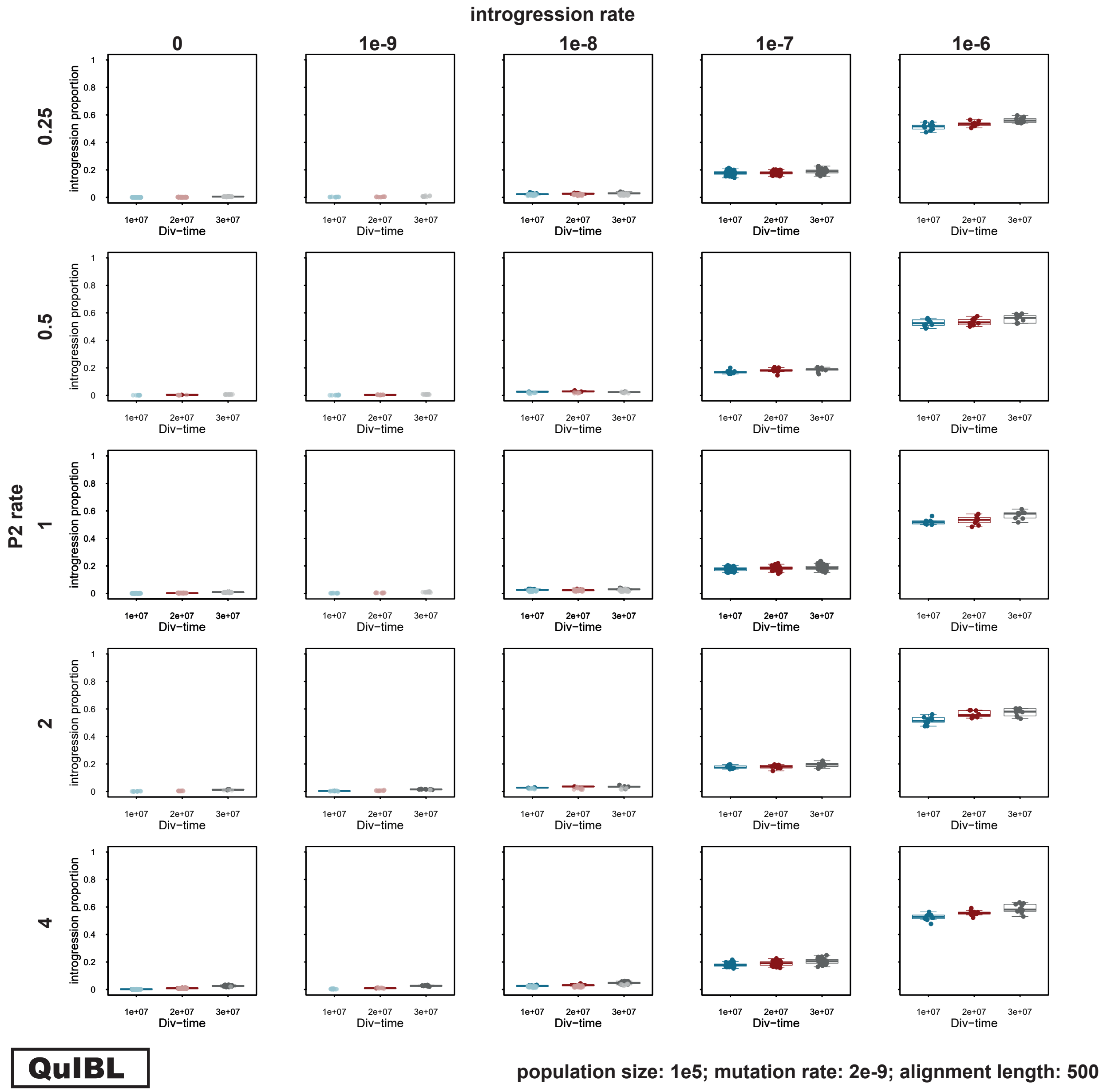

**Supplementary Figure S13:** Signals of introgression detected with QuIBL for datasets simulated with a population size $N_e = 10^5$, a mutation rate $\mu = 2 \times 10^{-9}$, an introgression rate $m \in \{0, 10^{-9}, 10^{-8}, 10^{-7}, 10^{-6}\}$, a P2 branch rate $s \in \{0.25, 0.5, 1, 2, 4\}$, a divergence time $t_{\text{P1,P2}} \in \{1 \times 10^7, 2 \times 10^7, 3 \times 10^7\}$, and an alignment length of 1,000 bp.

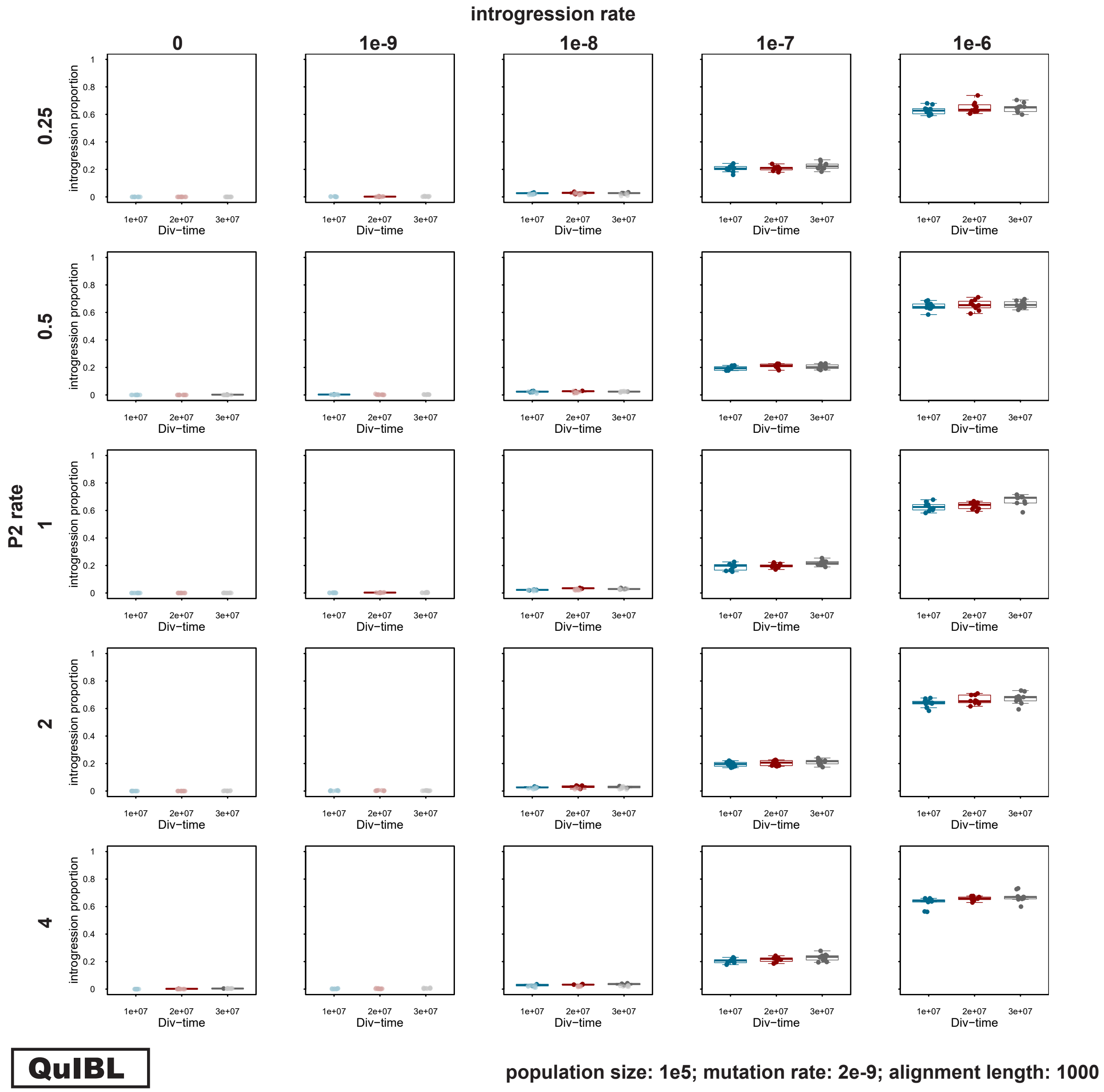

**Supplementary Figure S14:** Signals of introgression detected with the "MMS17 method" for datasets simulated with a population size $N_\mathrm{e} = 10^5$, a mutation rate $\mu = 2 \times 10^{-9}$, an introgression rate $m \in \{0, 10^{-9}, 10^{-8}, 10^{-7}, 10^{-6}\}$, a P2 branch rate $s \in \{0.25, 0.5, 1, 2, 4\}$, a divergence time $t_{\mathrm{P1,P2}} \in \{1 \times 10^7, 2 \times 10^7, 3 \times 10^7\}$, and an alignment length of 200 bp. Gray crosshairs indicate results based on incorrectly inferred topologies.

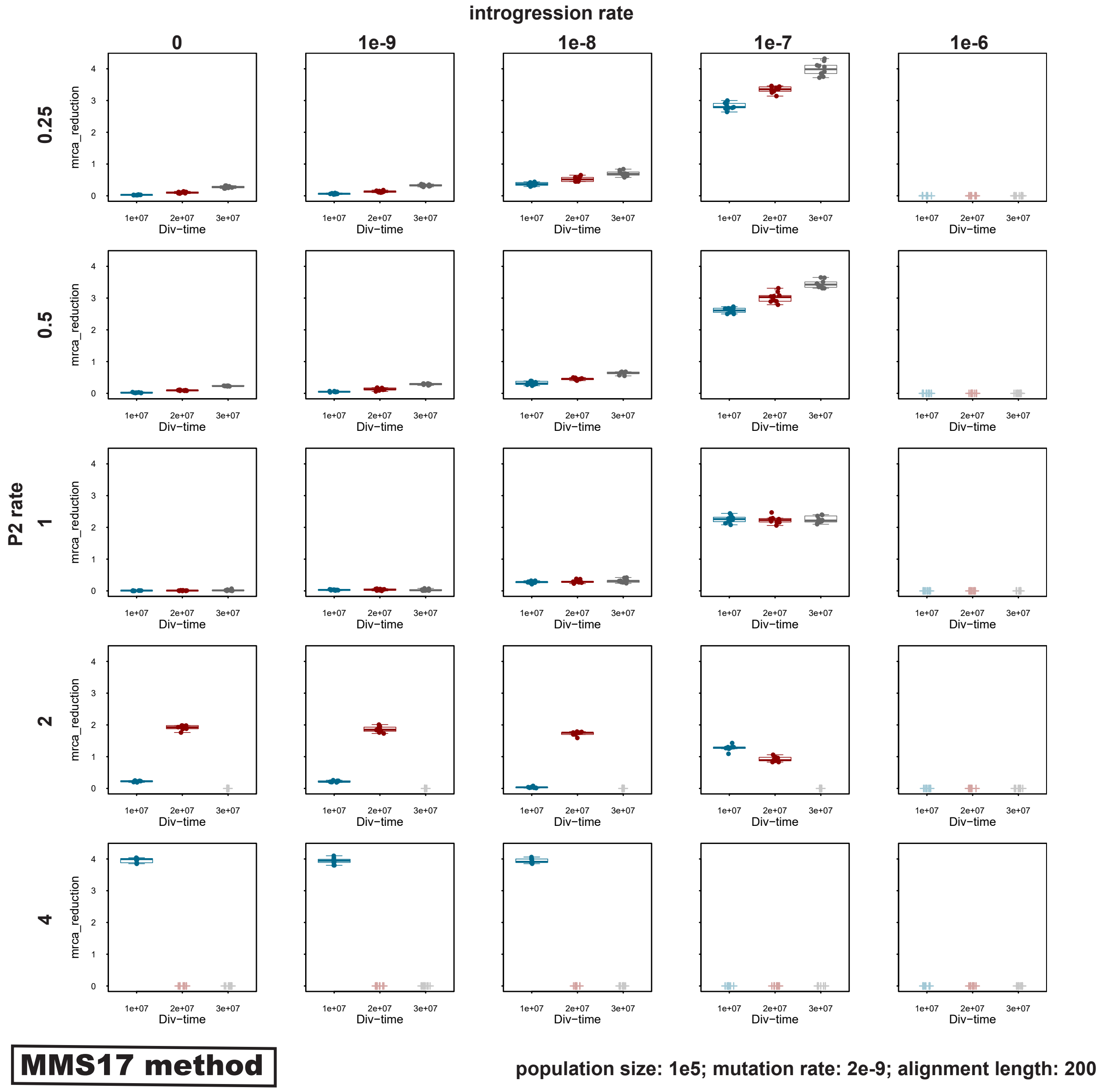

**Supplementary Figure S15:** Signals of introgression detected with the "MMS17 method" for datasets simulated with a population size $N_e = 10^5$, a mutation rate $\mu = 2 \times 10^{-9}$, an introgression rate $m \in \{0, 10^{-9}, 10^{-8}, 10^{-7}, 10^{-6}\}$, a P2 branch rate $s \in \{0.25, 0.5, 1, 2, 4\}$, a divergence time $t_{\text{P1,P2}} \in \{1 \times 10^7, 2 \times 10^7, 3 \times 10^7\}$, and an alignment length of 500 bp. For introgression rate $m \in \{0, 10^{-8}, 10^{-7}\}$, and a P2 branch rate $s \in \{0.25, 1, 4\}$ 50 replicates were simulated, for all the other parameter combinations we performed ten replicate simulations. Gray crosshairs indicate results based on incorrectly inferred topologies.

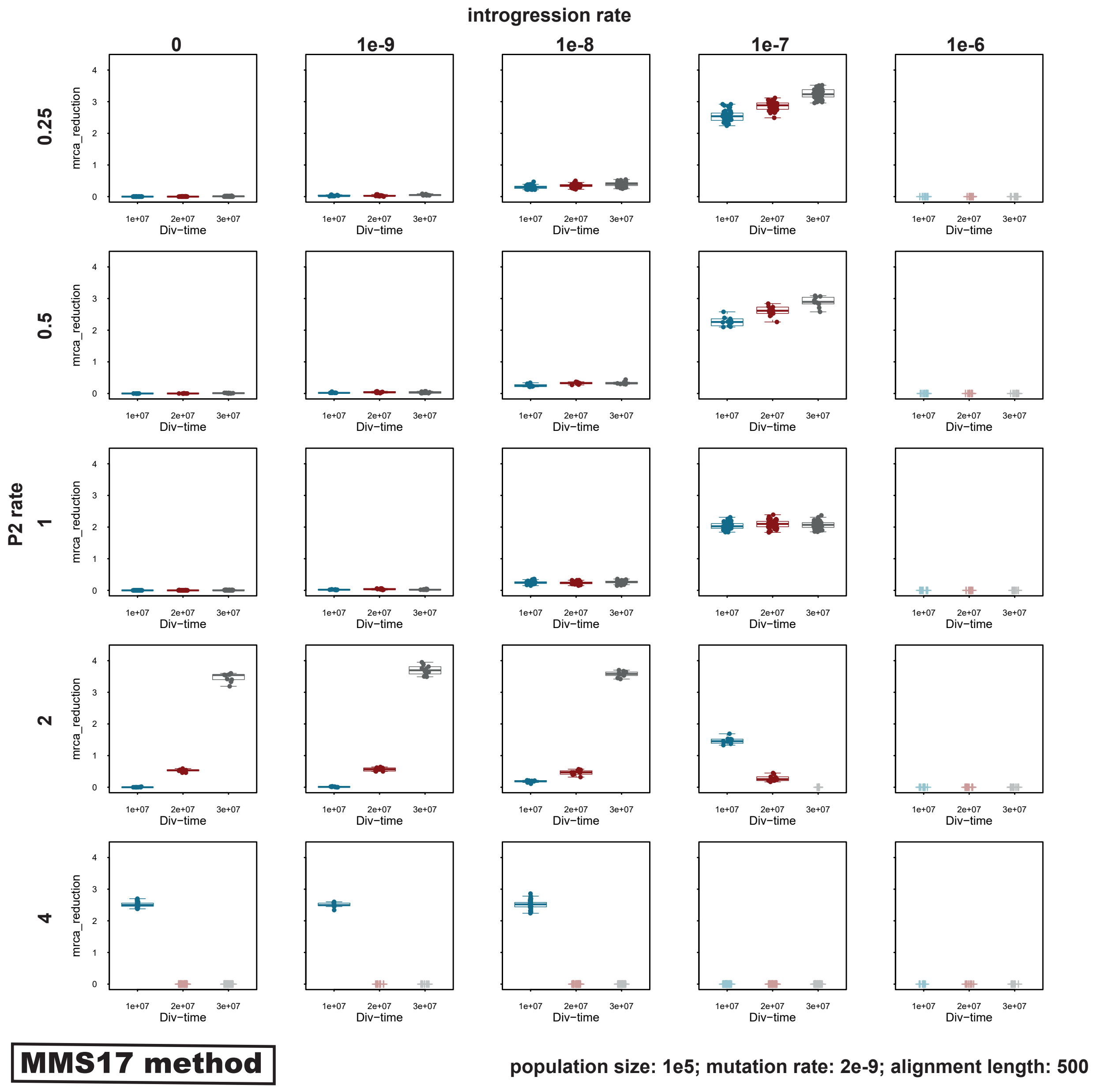

**Supplementary Figure S16:** Signals of introgression detected with the "MMS17 method" for datasets simulated with a population size $N_e = 10^5$, a mutation rate $\mu = 2 \times 10^{-9}$, an introgression rate $m \in \{0, 10^{-9}, 10^{-8}, 10^{-7}, 10^{-6}\}$, a P2 branch rate $s \in \{0.25, 0.5, 1, 2, 4\}$, a divergence time $t_{\mathrm{P1,P2}} \in \{1 \times 10^7, 2 \times 10^7, 3 \times 10^7\}$, and an alignment length of 1,000 bp. Gray crosshairs indicate results based on incorrectly inferred topologies.

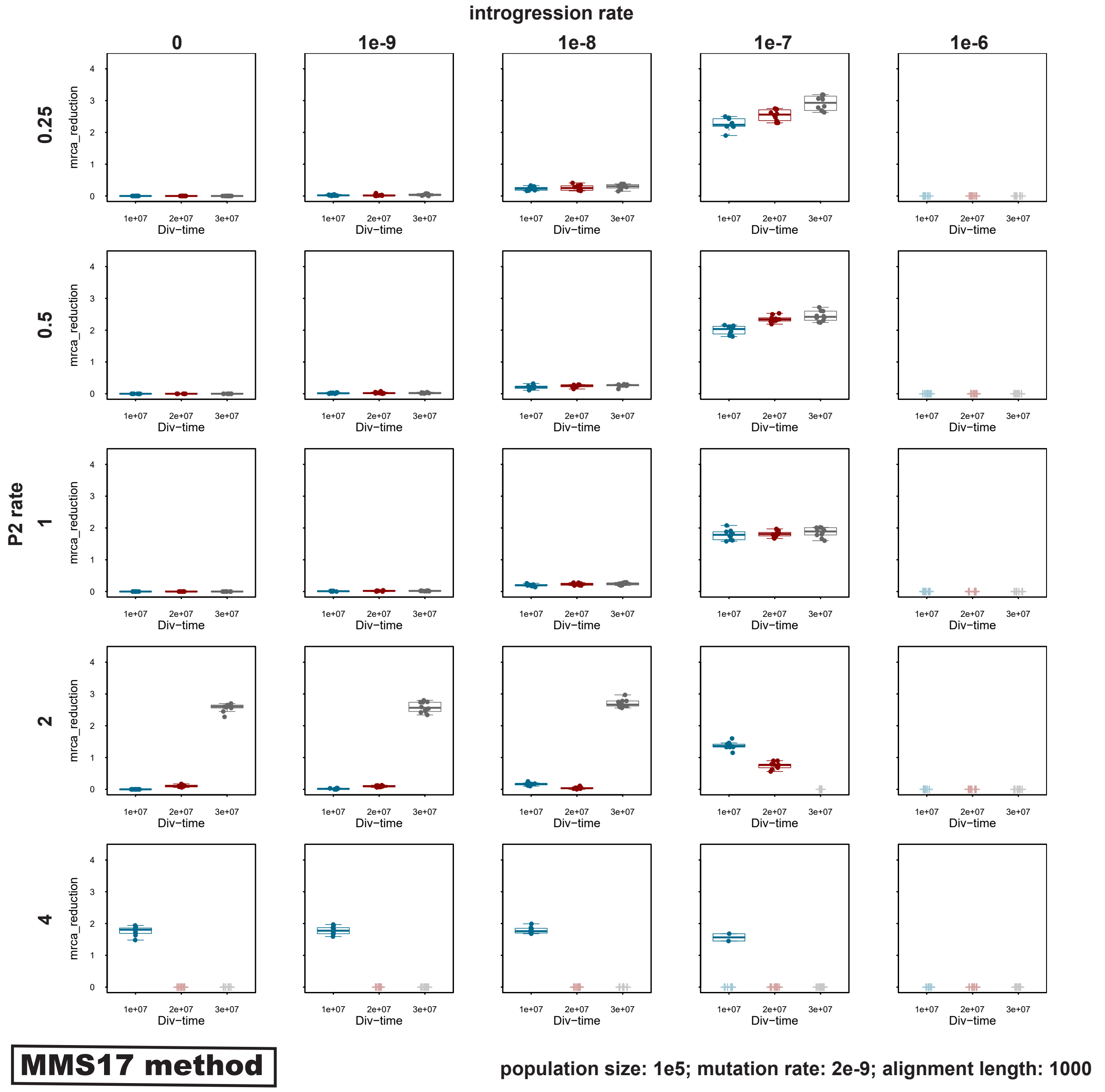

**Supplementary Figure S17:** Signals of introgression detected with the "sensitive" version of the 'ABBA'-site clustering test for datasets simulated with a population size $N_{\mathrm{e}} = 10^5$, a mutation rate $\mu = 1 \times 10^{-9}$, an introgression rate $m \in \{0, 10^{-9}, 10^{-8}, 10^{-7}, 10^{-6}\}$, a P2 branch rate $s \in \{0.25, 0.5, 1, 2, 4\}$ and a divergence time $t_{\mathrm{P1,P2}} \in \{1 \times 10^7, 2 \times 10^7, 3 \times 10^7\}$.

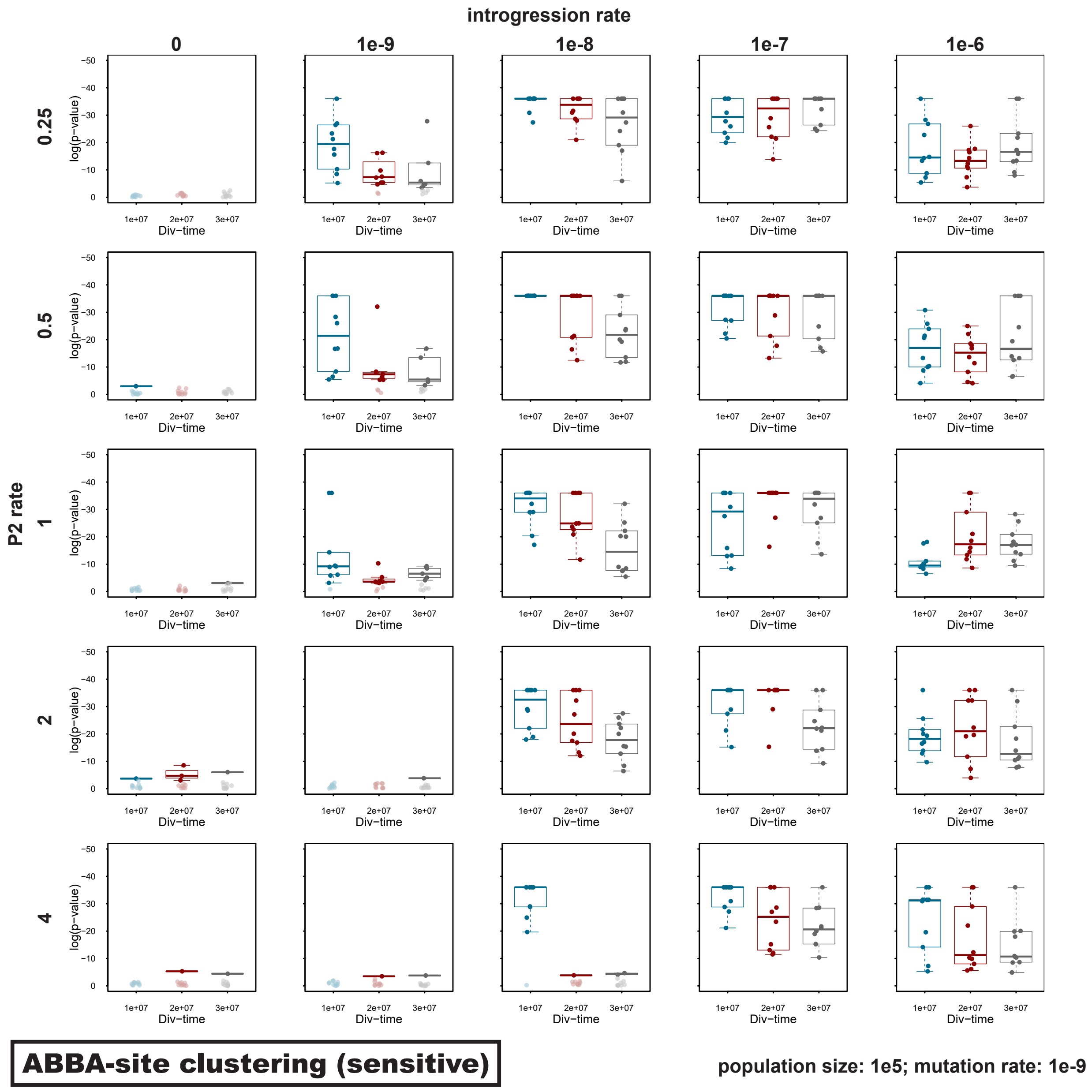

**Supplementary Figure S18:** Signals of introgression detected with the "sensitive" version of the 'ABBA'-site clustering test for datasets simulated with a population size $N_e = 10^5$, a mutation rate $\mu = 2 \times 10^{-9}$, an introgression rate $m \in \{0, 10^{-9}, 10^{-8}, 10^{-7}, 10^{-6}\}$, a P2 branch rate $s \in \{0.25, 0.5, 1, 2, 4\}$ and a divergence time $t_{P1,P2} \in \{1 \times 10^7, 2 \times 10^7, 3 \times 10^7\}$. For introgression rate $m \in \{0, 10^{-8}, 10^{-7}\}$ and a P2 branch rate $s \in \{0.25, 1, 4\}$, 50 replicates were simulated; for all other parameter combinations, we performed ten replicate simulations.

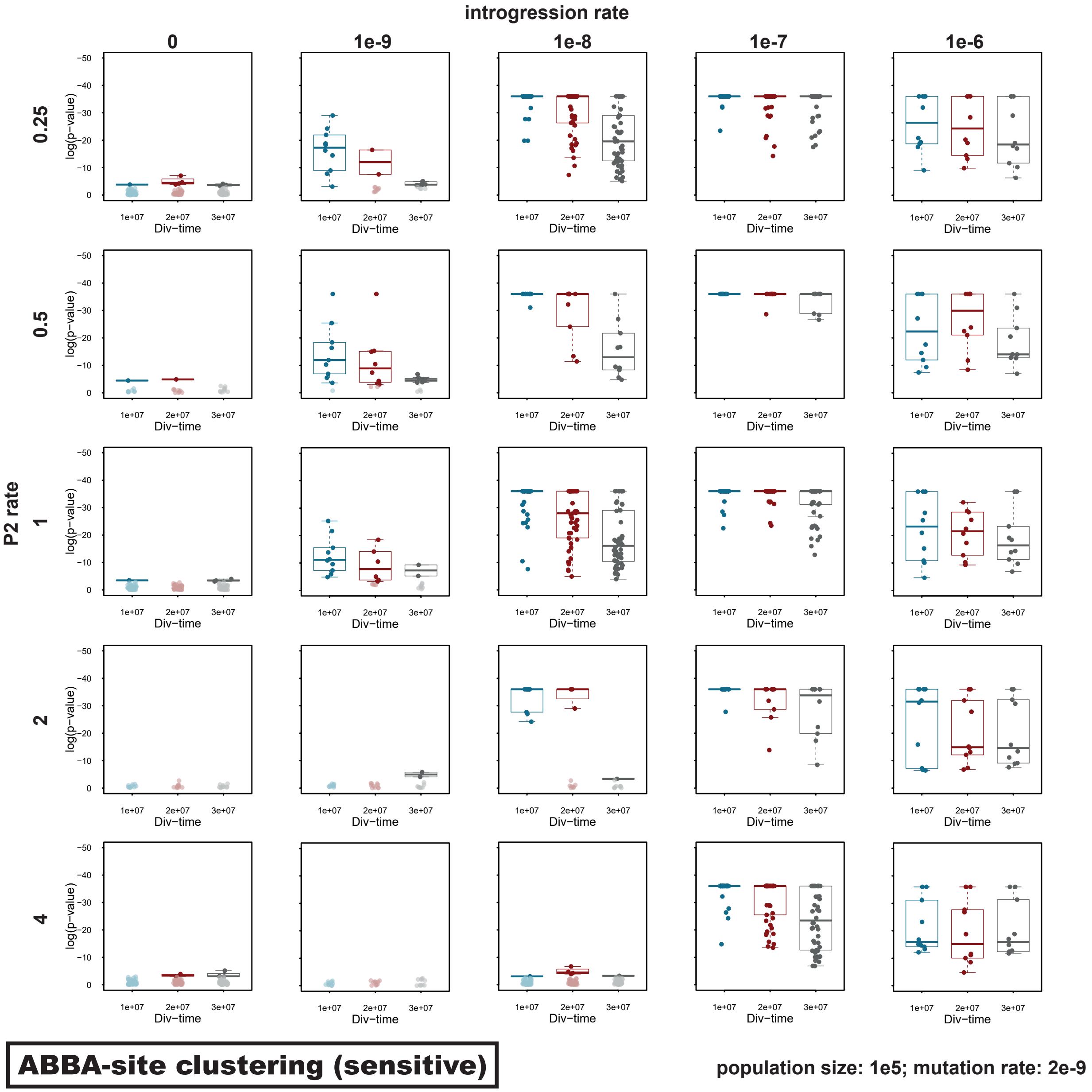

**Supplementary Figure S19:** Signals of introgression detected with the "sensitive" version of the 'ABBA'-site clustering test for datasets simulated with a population size $N_\mathrm{e} = 10^4$, a mutation rate $\mu = 1 \times 10^{-9}$, an introgression rate $m \in \{0, 10^{-9}, 10^{-8}, 10^{-7}, 10^{-6}\}$, a P2 branch rate $s \in \{0.25, 0.5, 1, 2, 4\}$ and a divergence time $t_{\mathrm{P1,P2}} \in \{1 \times 10^7, 2 \times 10^7, 3 \times 10^7\}$.

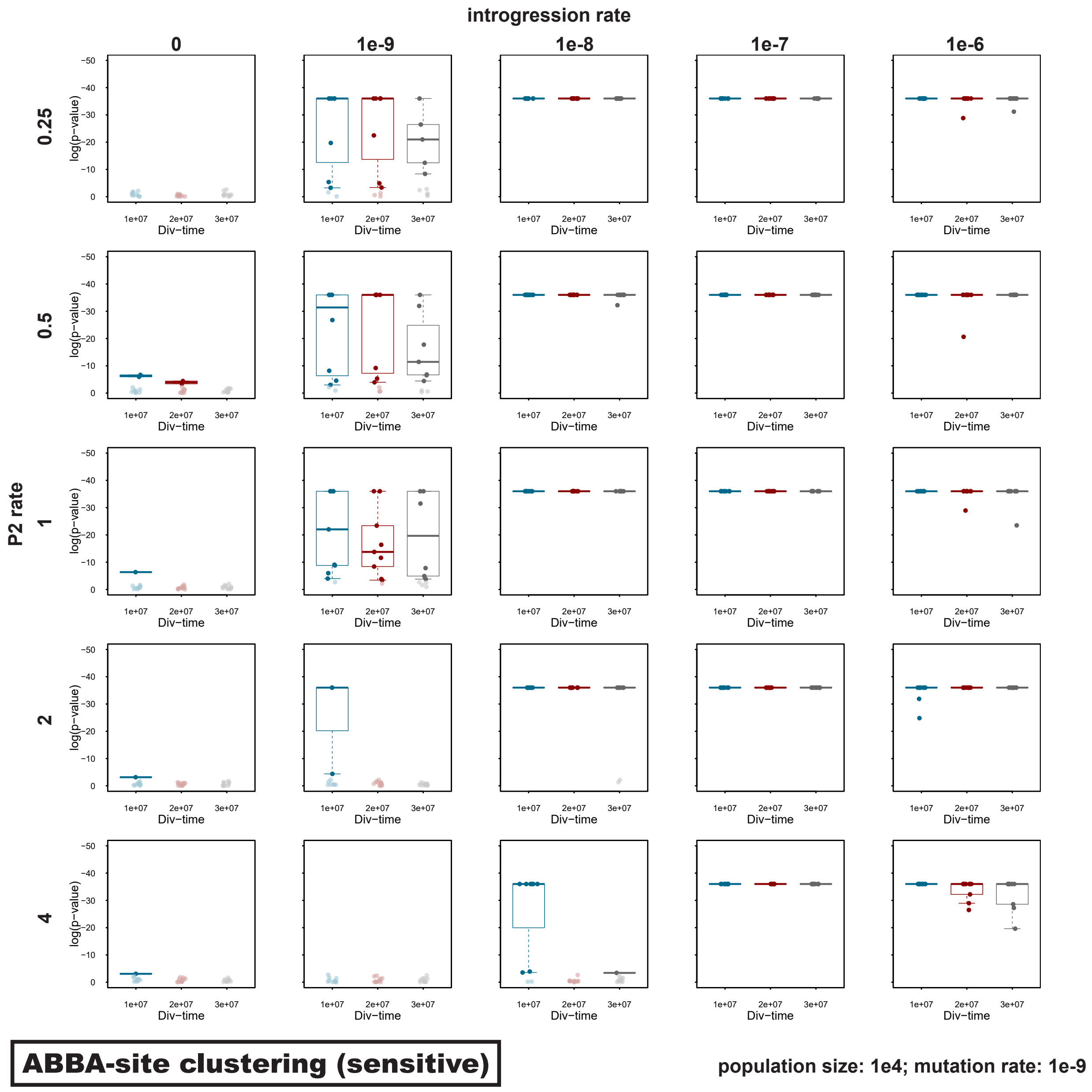

**Supplementary Figure S20:** Signals of introgression detected with the "sensitive" version of the 'ABBA'-site clustering test for datasets simulated with a population size $N_e = 10^4$, a mutation rate $\mu = 2 \times 10^{-9}$, an introgression rate $m \in \{0, 10^{-9}, 10^{-8}, 10^{-7}, 10^{-6}\}$, a P2 branch rate $s \in \{0.25, 0.5, 1, 2, 4\}$ and a divergence time $t_{\mathrm{P1,P2}} \in \{1 \times 10^7, 2 \times 10^7, 3 \times 10^7\}$.

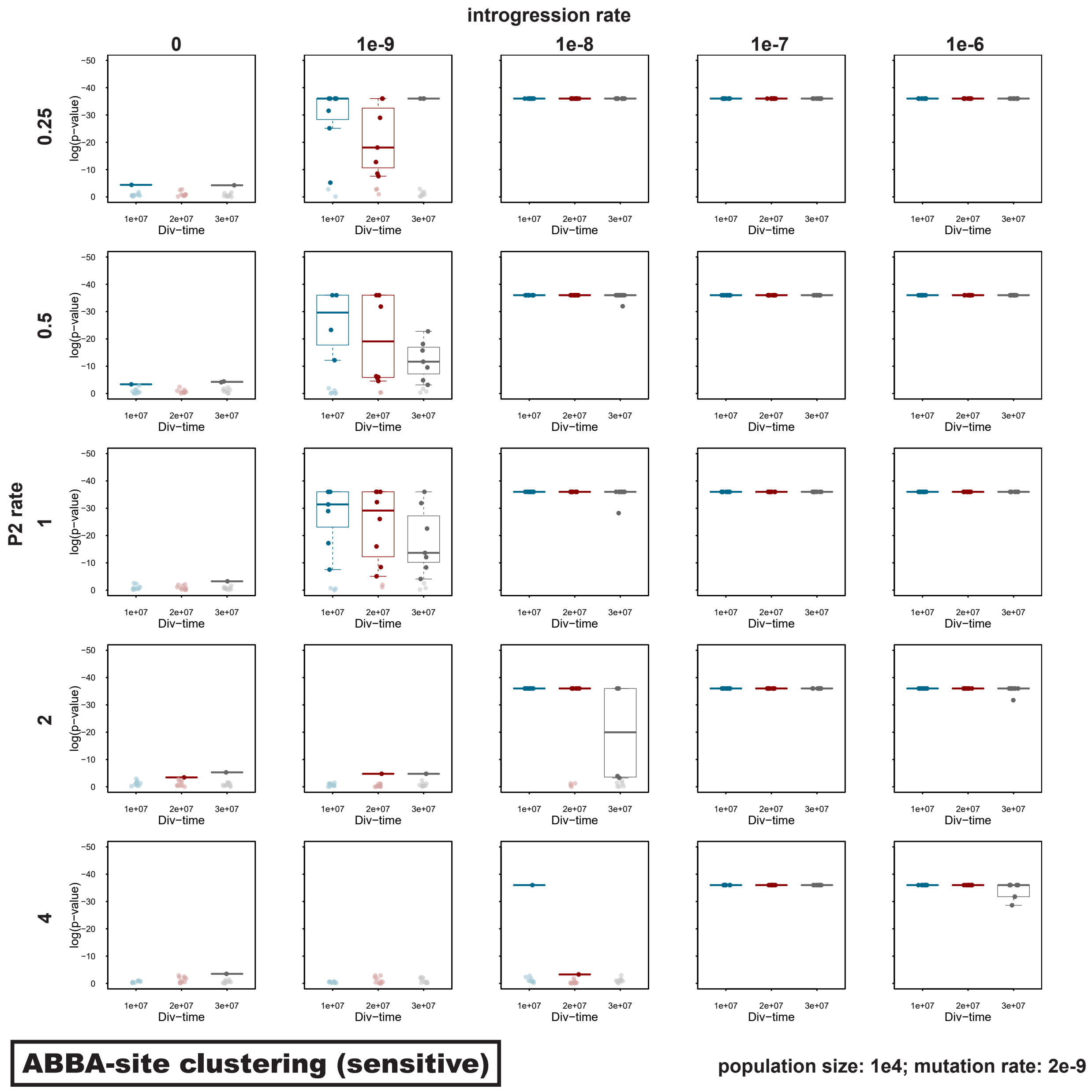

**Supplementary Figure S21:** Signals of introgression detected with the "robust" version of the 'ABBA'-site clustering test for datasets simulated with a population size $N_\mathrm{e} = 10^5$, a mutation rate $\mu = 1 \times 10^{-9}$, an introgression rate $m \in \{0, 10^{-9}, 10^{-8}, 10^{-7}, 10^{-6}\}$, a P2 branch rate $s \in \{0.25, 0.5, 1, 2, 4\}$ and a divergence time $t_{\mathrm{P1,P2}} \in \{1 \times 10^7, 2 \times 10^7, 3 \times 10^7\}$.

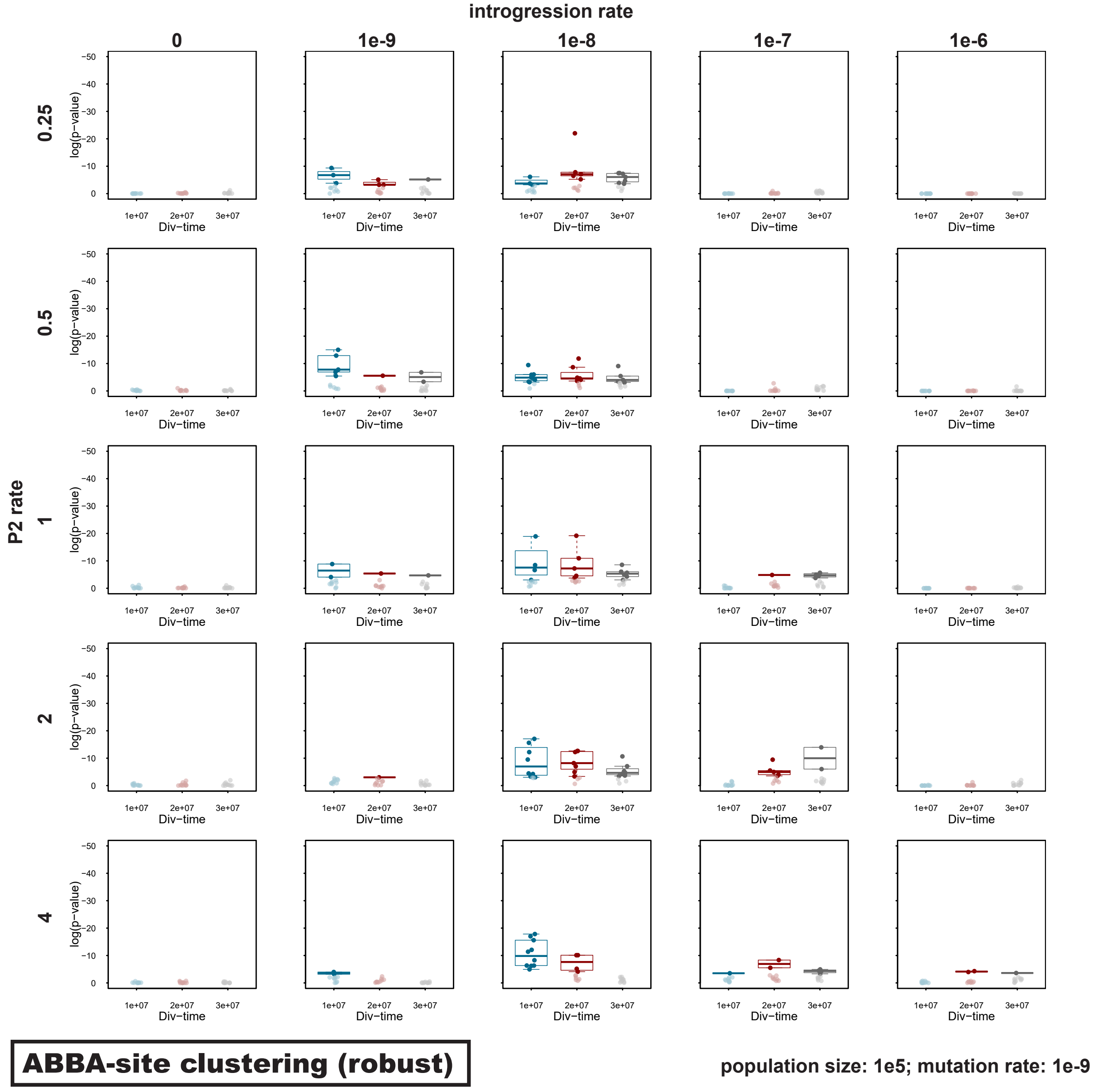

**Supplementary Figure S22:** Signals of introgression detected with the "robust" version of the 'ABBA'-site clustering test for datasets simulated with a population size $N_e = 10^5$, a mutation rate $\mu = 2 \times 10^{-9}$, an introgression rate $m \in \{0, 10^{-9}, 10^{-8}, 10^{-7}, 10^{-6}\}$, a P2 branch rate $s \in \{0.25, 0.5, 1, 2, 4\}$ and a divergence time $t_{P1,P2} \in \{1 \times 10^7, 2 \times 10^7, 3 \times 10^7\}$. For introgression rate $m \in \{0, 10^{-8}, 10^{-7}\}$ and a P2 branch rate $s \in \{0.25, 1, 4\}$, 50 replicates were simulated; for all other parameter combinations, we performed ten replicate simulations.

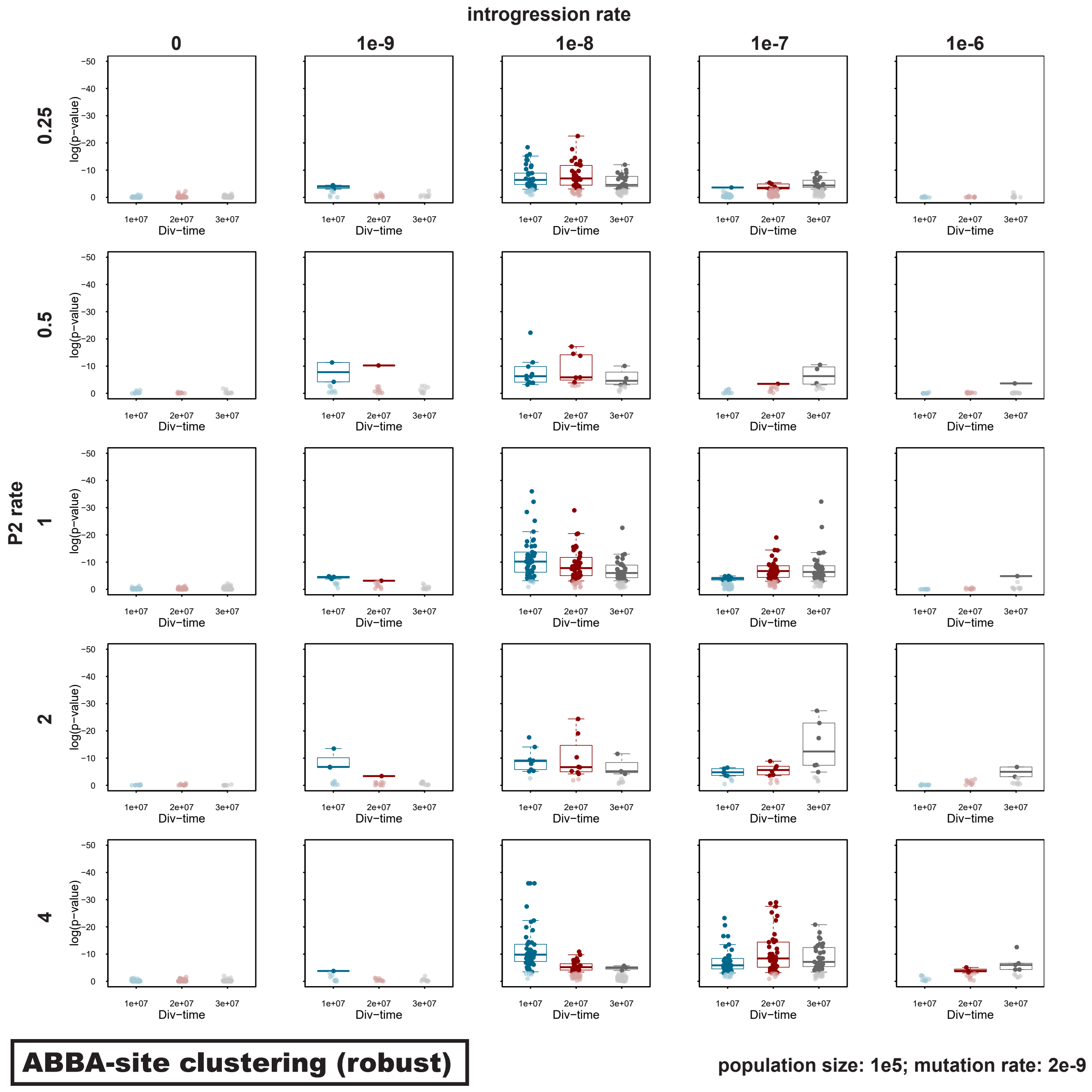

**Supplementary Figure S23:** Signals of introgression detected with the "robust" version of the 'ABBA'-site clustering test for datasets simulated with a population size $N_\mathrm{e} = 10^4$, a mutation rate $\mu = 1 \times 10^{-9}$, an introgression rate $m \in \{0, 10^{-9}, 10^{-8}, 10^{-7}, 10^{-6}\}$, a P2 branch rate $s \in \{0.25, 0.5, 1, 2, 4\}$ and a divergence time $t_{\mathrm{P1,P2}} \in \{1 \times 10^7, 2 \times 10^7, 3 \times 10^7\}$.

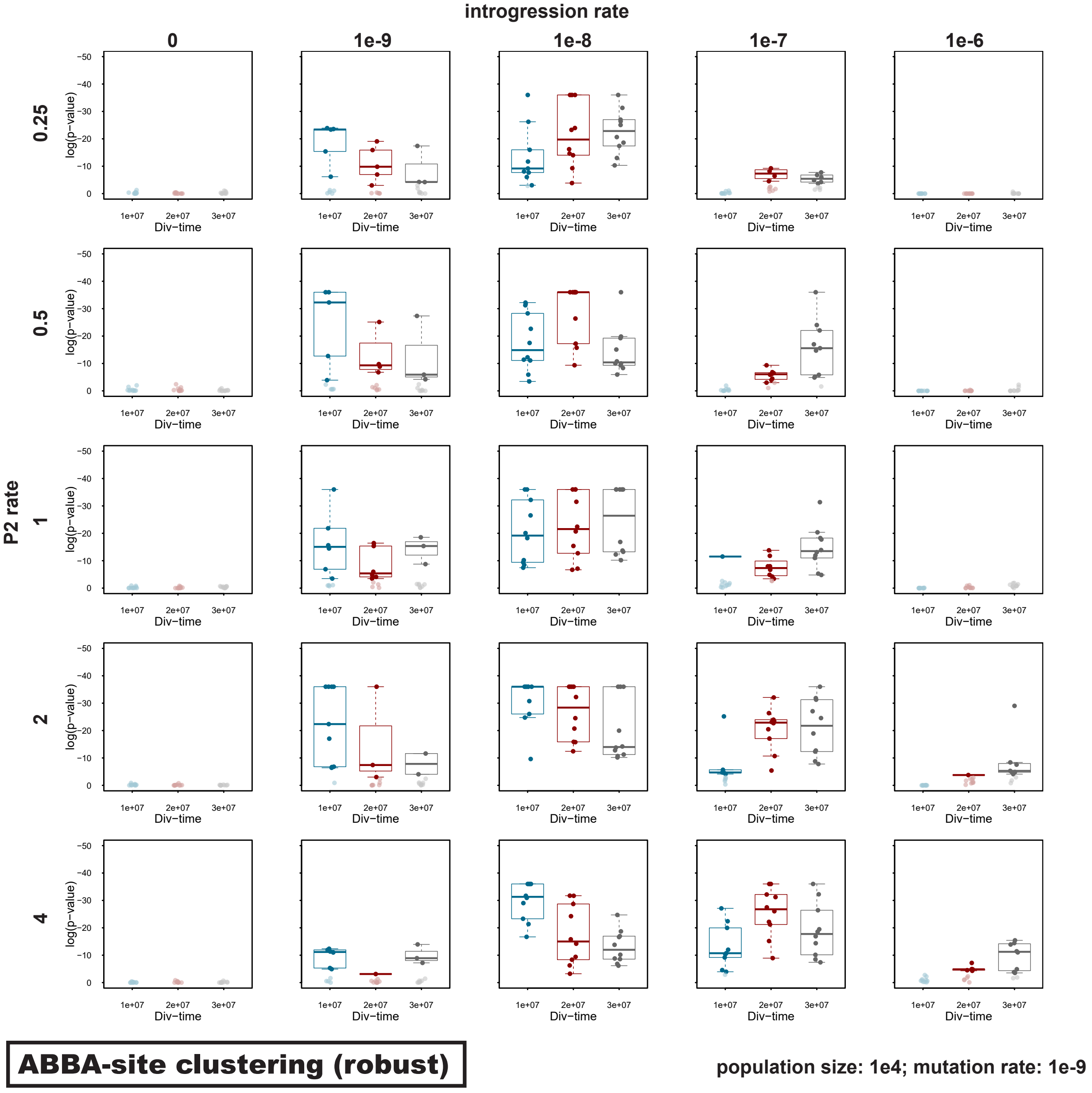

**Supplementary Figure S24:** Signals of introgression detected with the "robust" version of the 'ABBA'-site clustering test for datasets simulated with a population size $N_{\mathrm{e}} = 10^4$, a mutation rate $\mu = 2 \times 10^{-9}$, an introgression rate $m \in \{0, 10^{-9}, 10^{-8}, 10^{-7}, 10^{-6}\}$, a P2 branch rate $s \in \{0.25, 0.5, 1, 2, 4\}$ and a divergence time $t_{\mathrm{P1,P2}} \in \{1 \times 10^7, 2 \times 10^7, 3 \times 10^7\}$.

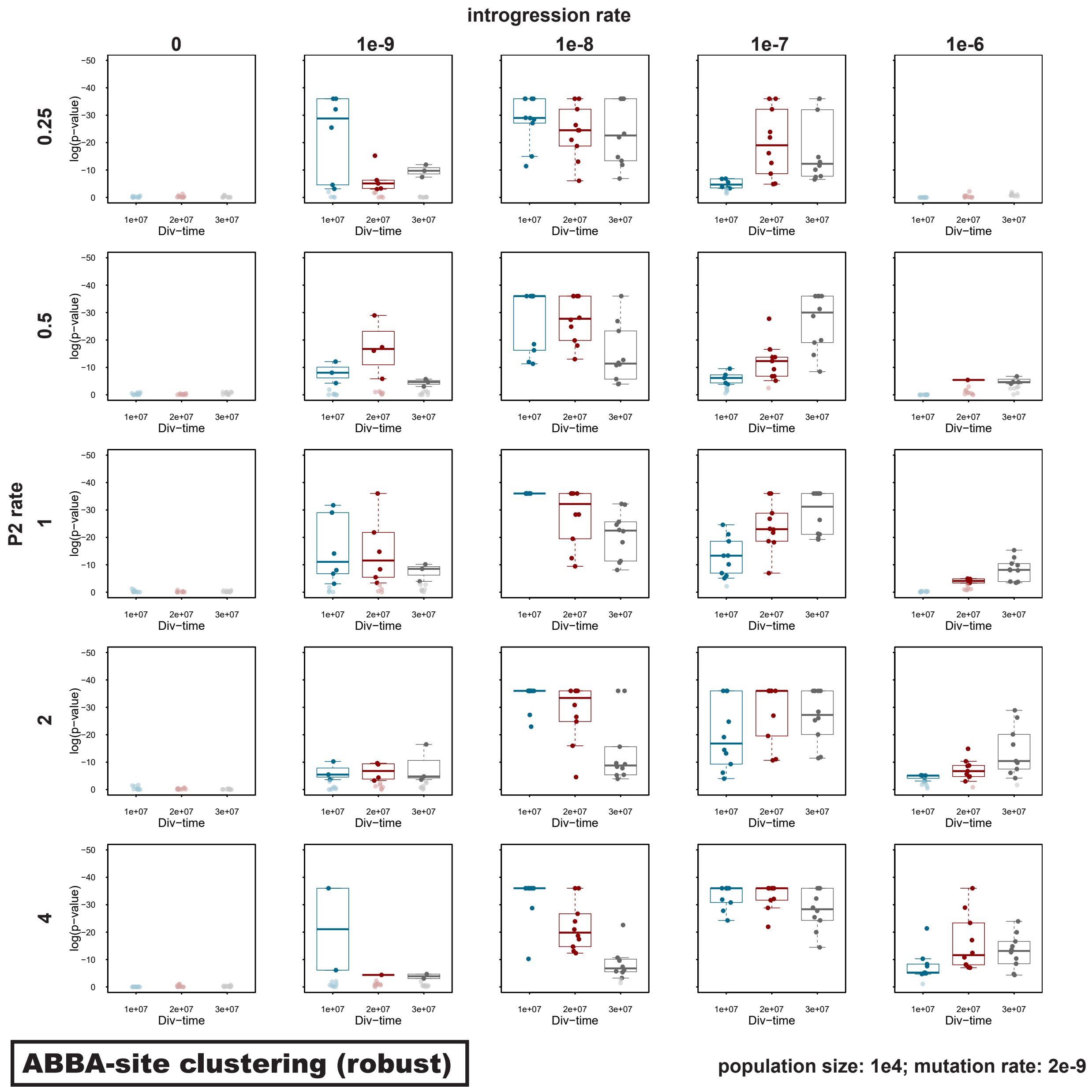

**Supplementary Figure S25:** Signals of introgression detected with the "sensitive" version of the 'ABBA'-site clustering test for datasets simulated with among-site recombination-rate variation (using a human recombination map; see Supplementary Note S2), a population size $N_e = 10^5$, a mutation rate $\mu = 2 \times 10^{-9}$, an introgression rate $m \in \{0, 10^{-8}, 10^{-7}\}$, a P2 branch rate $s \in \{0.25, 1, 4\}$ and a divergence time $t_{P1,P2} \in \{1 \times 10^7, 2 \times 10^7, 3 \times 10^7\}$.

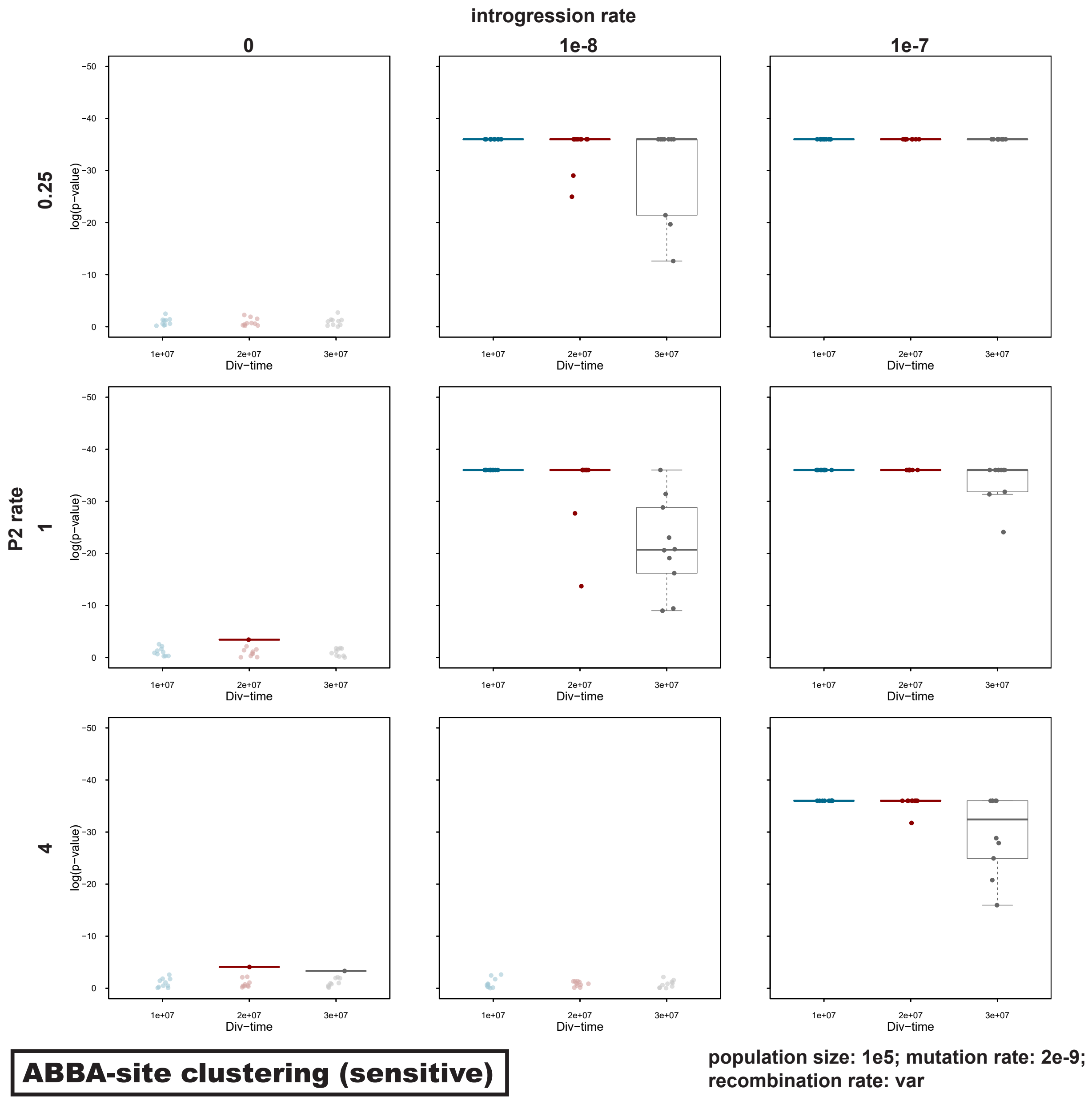

**Supplementary Figure S26:** Signals of introgression detected with the "robust" version of the 'ABBA'-site clustering test for datasets simulated with among-site recombination-rate variation (using a human recombination map; see Supplementary Note S2), a population size $N_e = 10^5$, a mutation rate $\mu = 2 \times 10^{-9}$, an introgression rate $m \in \{0, 10^{-8}, 10^{-7}\}$, a P2 branch rate $s \in \{0.25, 1, 4\}$ and a divergence time $t_{\mathrm{P1,P2}} \in \{1 \times 10^7, 2 \times 10^7, 3 \times 10^7\}$.

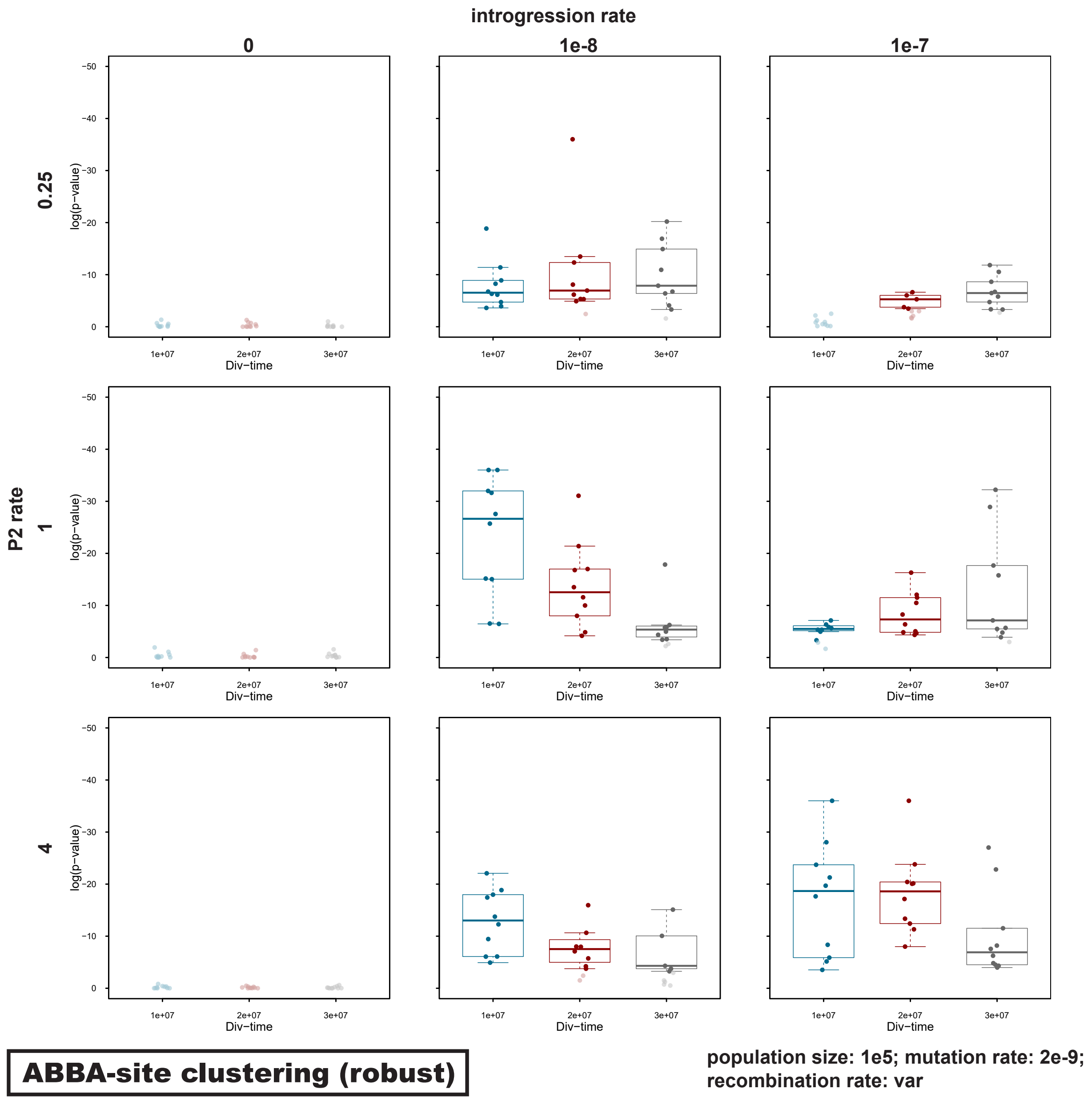

**Supplementary Figure S27:** Signals of introgression detected with the "sensitive" version of the 'ABBA'-site clustering test for datasets simulated with a moderate level of ILS (using a shortened internal branch with a length of 1 million generations; see Supplementary Note S4), a population size $N_{\mathrm{e}} \in \{10^4, 10^5\}$, a mutation rate $\mu \in \{1 \times 10^{-9}, 2 \times 10^{-9}\}$, an introgression rate $m = 0$, a P2 branch rate $s = 1$, and a divergence time $t_{\mathrm{P1,P2}} \in \{1 \times 10^7, 2 \times 10^7, 3 \times 10^7\}$.

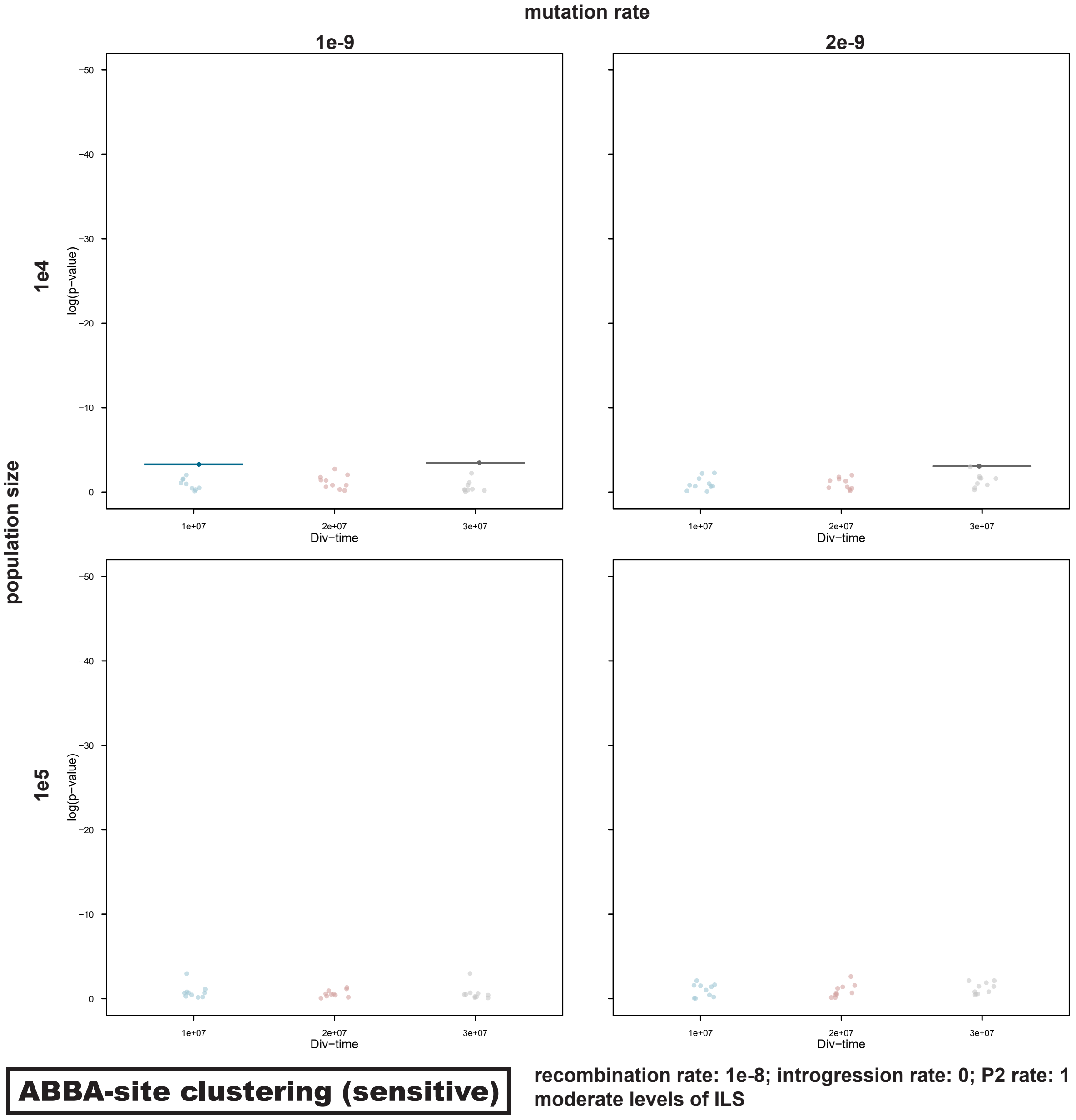

**Supplementary Figure S28:** Signals of introgression detected with the "sensitive" version of the 'ABBA'-site clustering test for datasets simulated with a high level of ILS (using a shortened internal branch with a length of 100,000 generations; see Supplementary Note S4), a population size $N_e \in \{10^4, 10^5\}$, a mutation rate $\mu \in \{1 \times 10^{-9}, 2 \times 10^{-9}\}$, an introgression rate $m = 0$, a P2 branch rate $s = 1$, and a divergence time $t_{P1,P2} \in \{1 \times 10^7, 2 \times 10^7, 3 \times 10^7\}$.

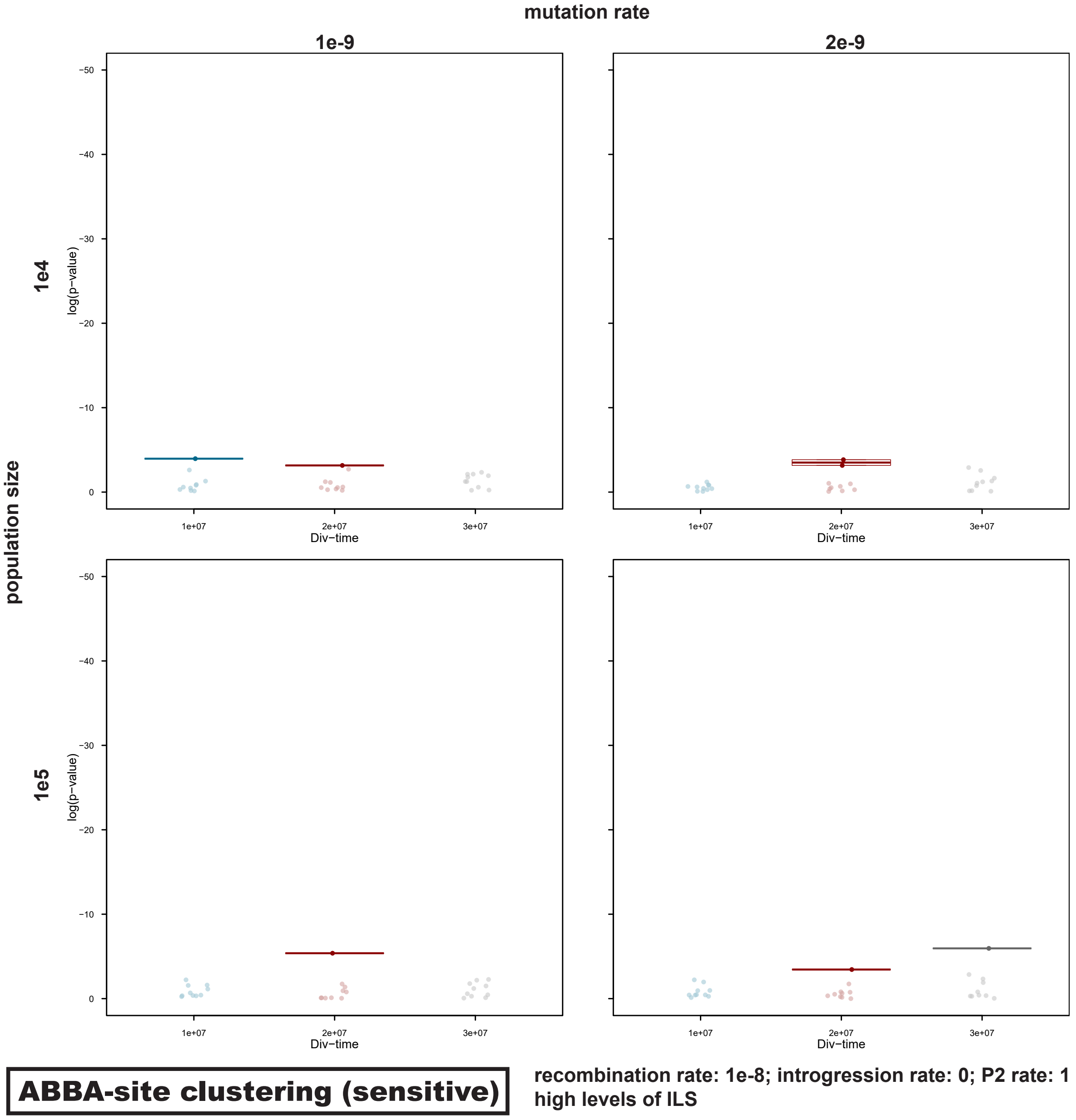

**Supplementary Figure S29:** Signals of introgression detected with the "sensitive" version of the 'ABBA'-site clustering test for datasets simulated with a very high level of ILS (using a shortened internal branch with a length of 10,000 generations; see Supplementary Note S4), a population size $N_e \in \{10^4, 10^5\}$, a mutation rate $\mu \in \{1 \times 10^{-9}, 2 \times 10^{-9}\}$, an introgression rate $m = 0$, a P2 branch rate $s = 1$, and a divergence time $t_{P1,P2} \in \{1 \times 10^7, 2 \times 10^7, 3 \times 10^7\}$.

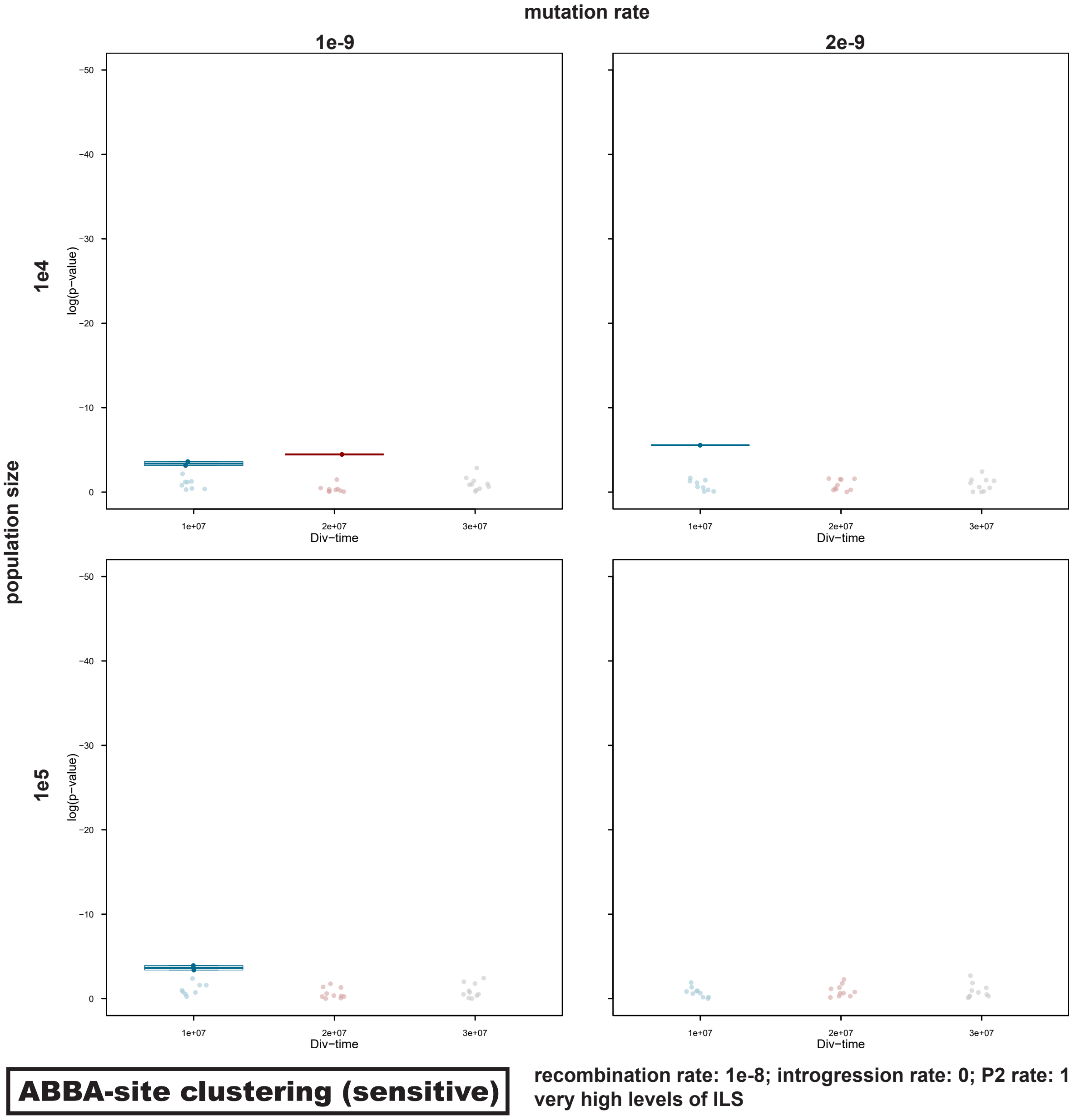

**Supplementary Figure S30:** Signals of introgression detected with the "robust" version of the 'ABBA'-site clustering test for datasets simulated with a moderate level of ILS (using a shortened internal branch with a length of 1 million generations; see Supplementary Note S4), a population size $N_\mathrm{e} \in \{10^4, 10^5\}$, a mutation rate $\mu \in \{1 \times 10^{-9}, 2 \times 10^{-9}\}$, an introgression rate $m = 0$, a P2 branch rate $s = 1$, and a divergence time $t_{\mathrm{P1,P2}} \in \{1 \times 10^7, 2 \times 10^7, 3 \times 10^7\}$.

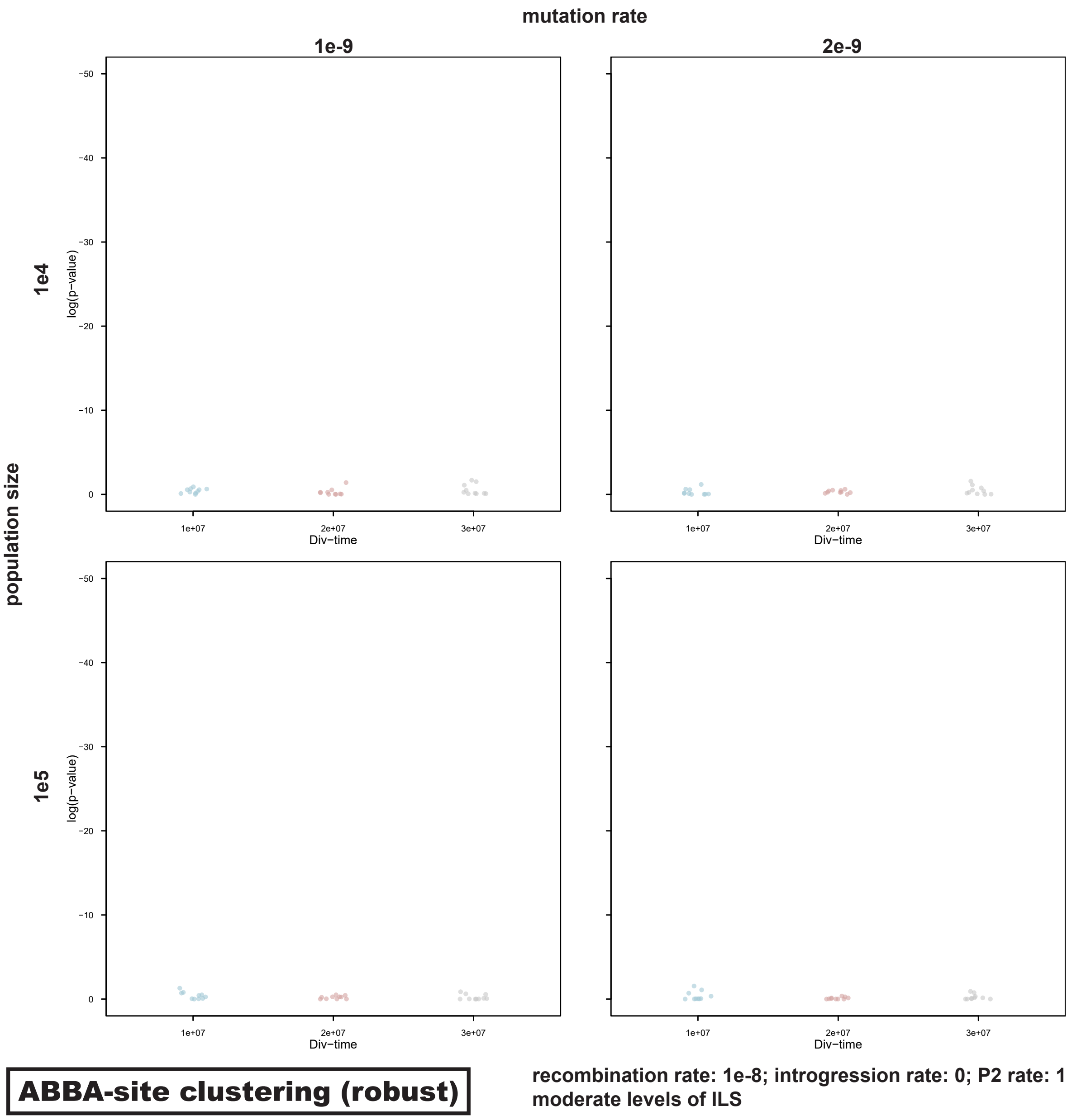

**Supplementary Figure S31:** Signals of introgression detected with the "robust" version of the 'ABBA'-site clustering test for datasets simulated with a high level of ILS (using a shortened internal branch with a length of 100,000 generations; see Supplementary Note S4), a population size $N_{\mathrm{e}} \in \{10^4, 10^5\}$, a mutation rate $\mu \in \{1 \times 10^{-9}, 2 \times 10^{-9}\}$, an introgression rate $m = 0$, a P2 branch rate $s = 1$, and a divergence time $t_{\mathrm{P1,P2}} \in \{1 \times 10^7, 2 \times 10^7, 3 \times 10^7\}$.

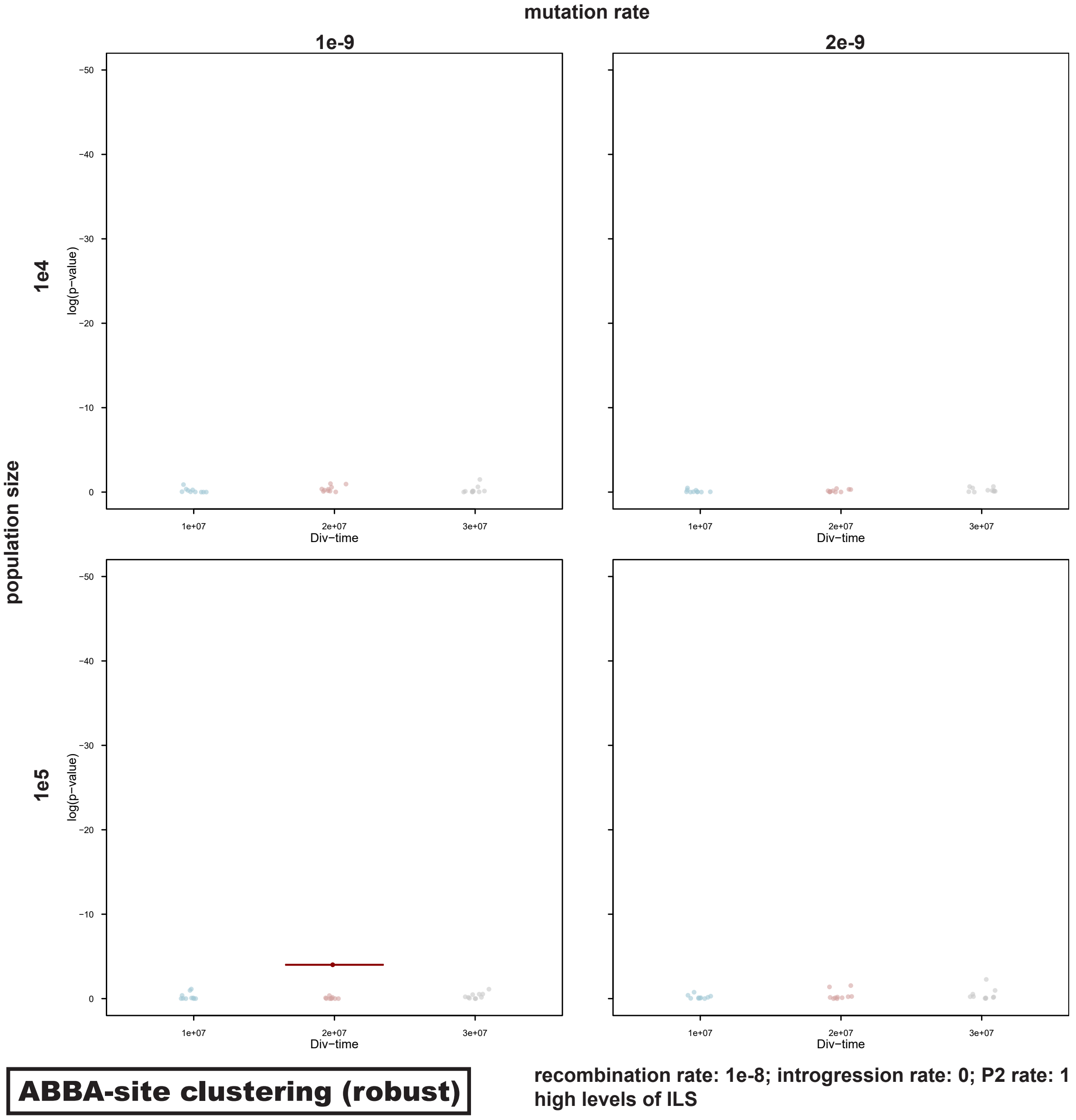

**Supplementary Figure S32:** Signals of introgression detected with the "robust" version of the 'ABBA'-site clustering test for datasets simulated with a very high level of ILS (using a shortened internal branch with a length of 10,000 generations; see Supplementary Note S4), a population size $N_{\mathrm{e}} \in \{10^4, 10^5\}$, a mutation rate $\mu \in \{1 \times 10^{-9}, 2 \times 10^{-9}\}$, an introgression rate $m = 0$, a P2 branch rate $s = 1$, and a divergence time $t_{\mathrm{P1,P2}} \in \{1 \times 10^7, 2 \times 10^7, 3 \times 10^7\}$.

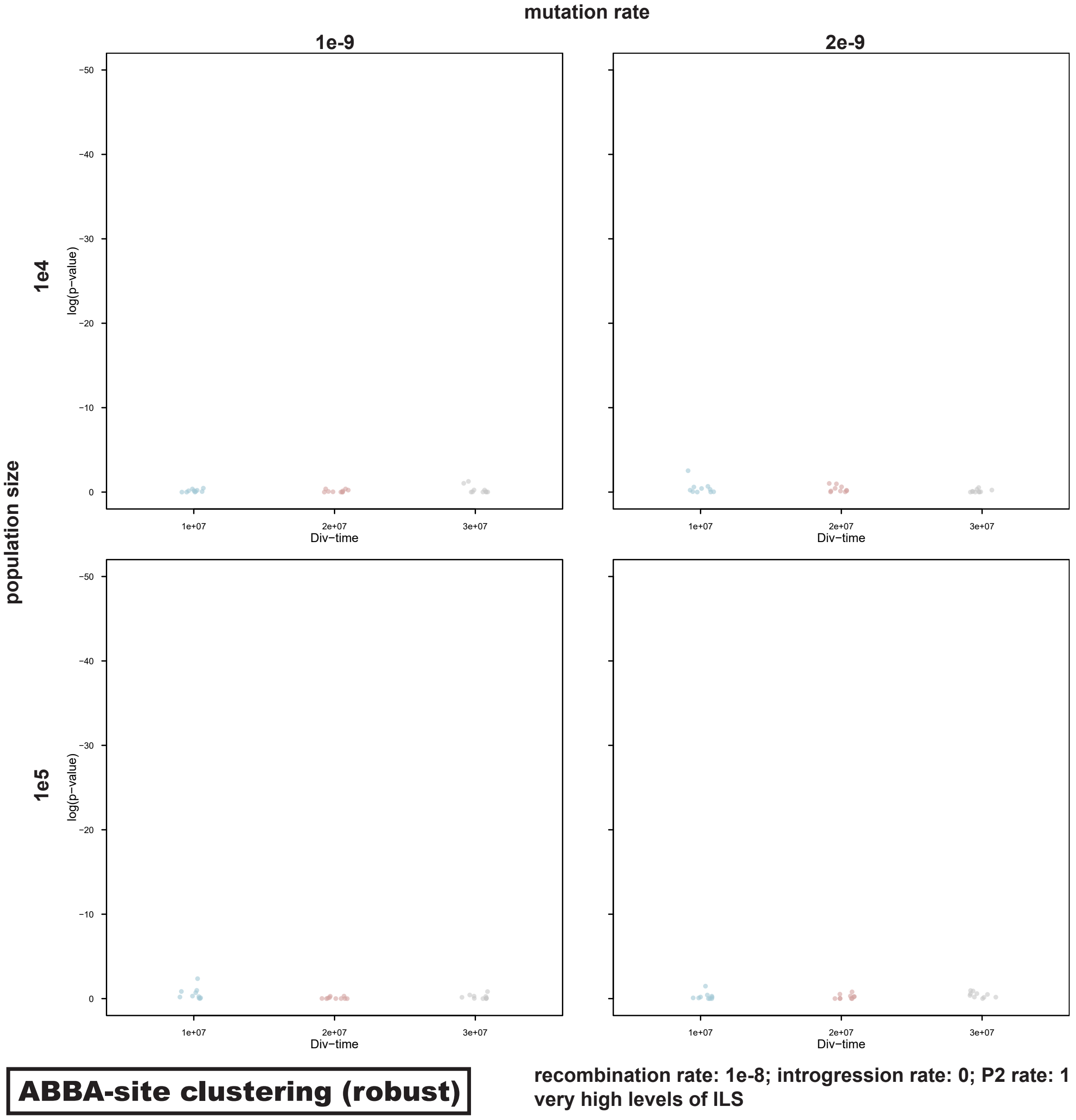

**Supplementary Figure S33:** Signals of introgression detected with the "sensitive" version of the 'ABBA'-site clustering test for datasets simulated with among-site mutation rate variation (using exponentially-distributed rates per 500-bp window; see Supplementary Note S3), a population size $N_e = 10^5$, a mutation rate $\mu = 2 \times 10^{-9}$, an introgression rate $m \in \{0, 10^{-8}, 10^{-7}\}$, a P2 branch rate $s \in \{0.25, 1, 4\}$, and a divergence time $t_{P1,P2} \in \{1 \times 10^7, 2 \times 10^7, 3 \times 10^7\}$.

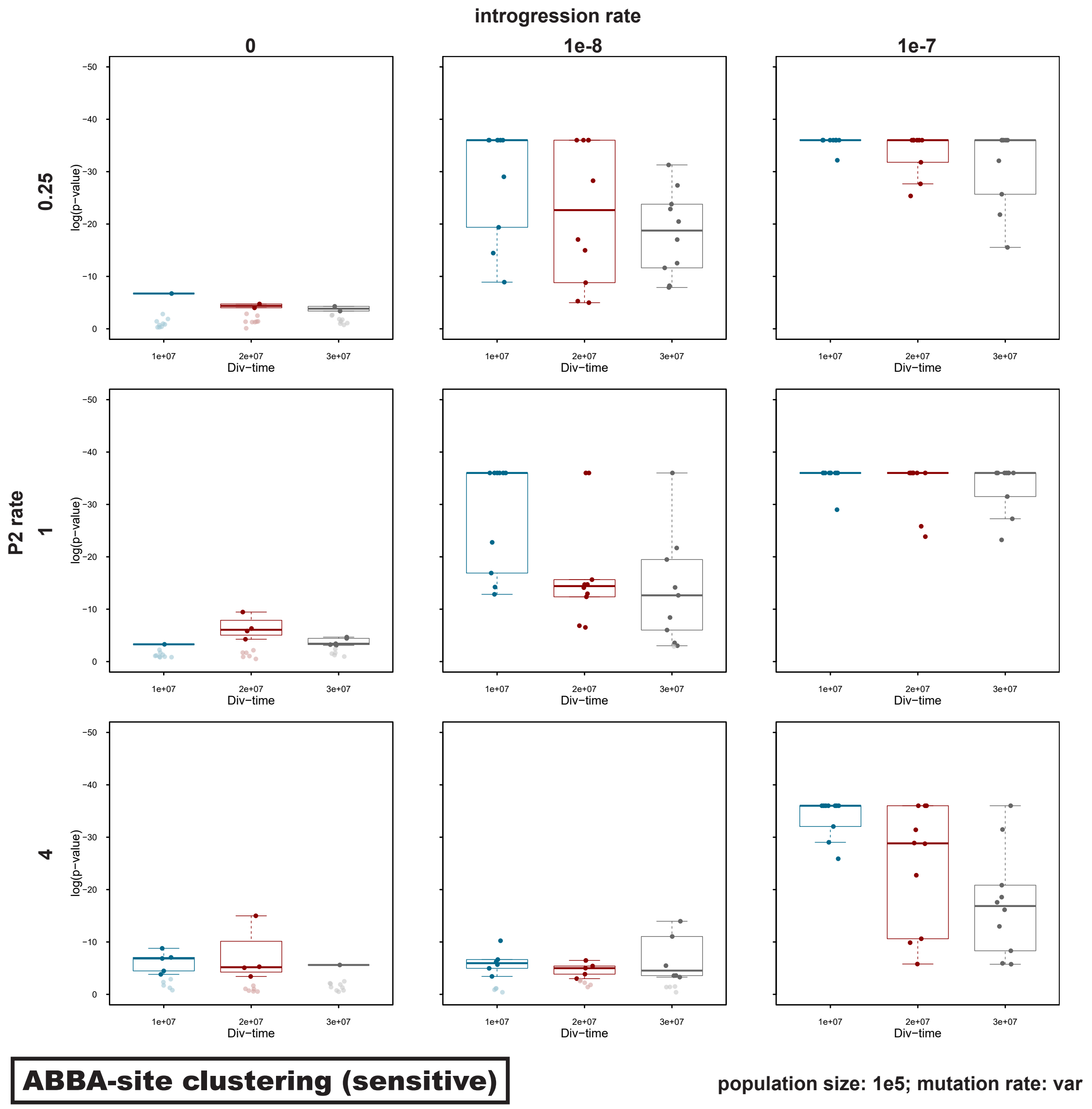

**Supplementary Figure S34:** Signals of introgression detected with the "robust" version of the 'ABBA'-site clustering test for datasets simulated with among-site mutation-rate variation (using exponentially-distributed rates per 500-bp window; see Supplementary Note S3), a population size $N_\mathrm{e} = 10^5$, a mutation rate $\mu = 2 \times 10^{-9}$, an introgression rate $m \in \{0, 10^{-8}, 10^{-7}\}$, a P2 branch rate $s \in \{0.25, 1, 4\}$, and a divergence time $t_{\mathrm{P1,P2}} \in \{1 \times 10^7, 2 \times 10^7, 3 \times 10^7\}$.

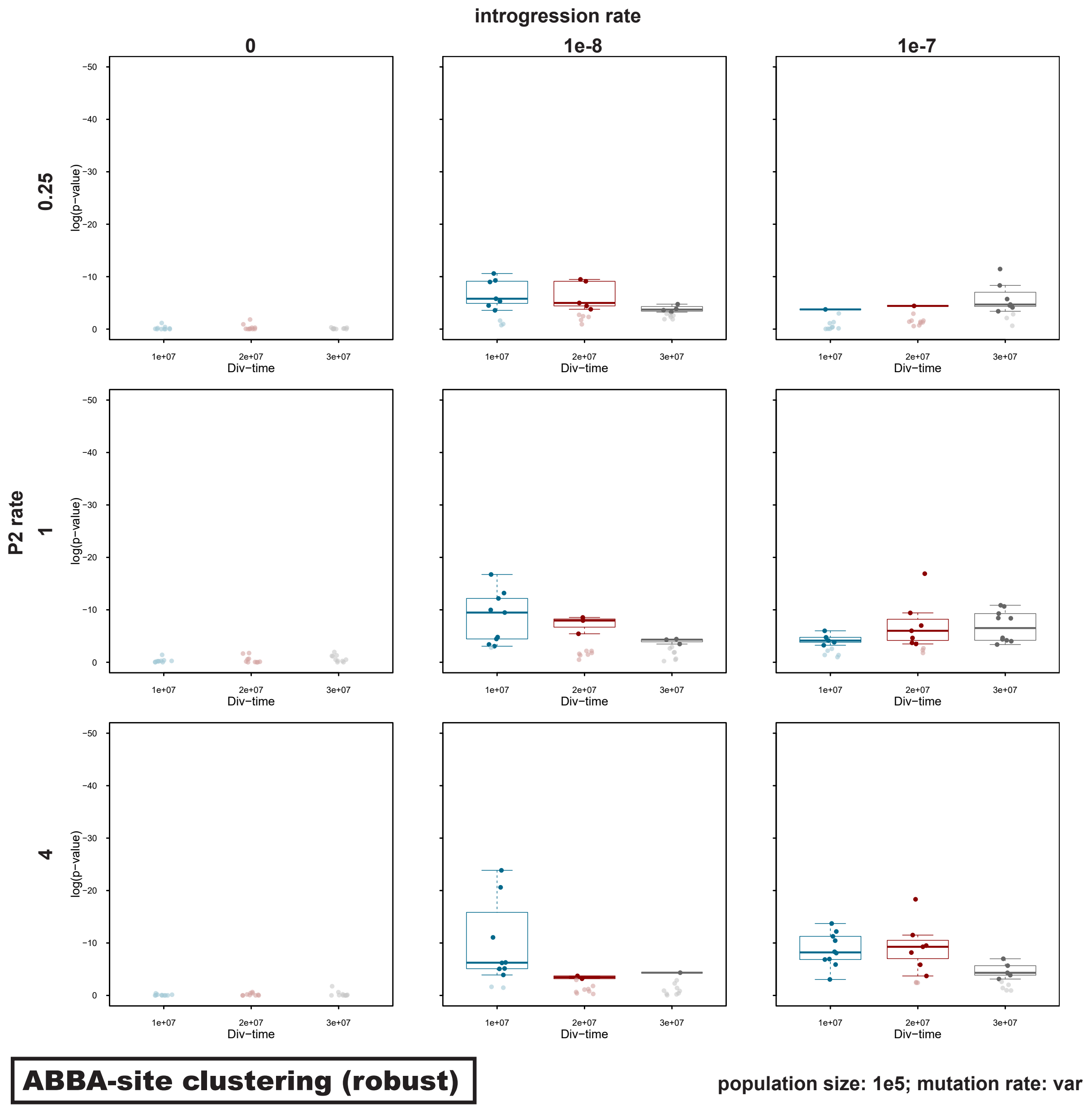

**Supplementary Figure S35:** Clustering of 'ABBA' sites in empirical data for Lake Tanganyika cichlid fishes, using the "sensitive" version of the 'ABBA'-site-clustering test. Results are shown per linkage group of the Nile tilapia reference assembly. With sorted "strong ABBA sites" on the horizontal axis, the black line indicates their position within a vector of all polymorphic sites on the vertical axis. A straight, diagonal line therefore illustrates a homogeneous distribution of these sites within this vector, while changes in the gradient illustrate clustering. Significant $p$-values are marked in bold.

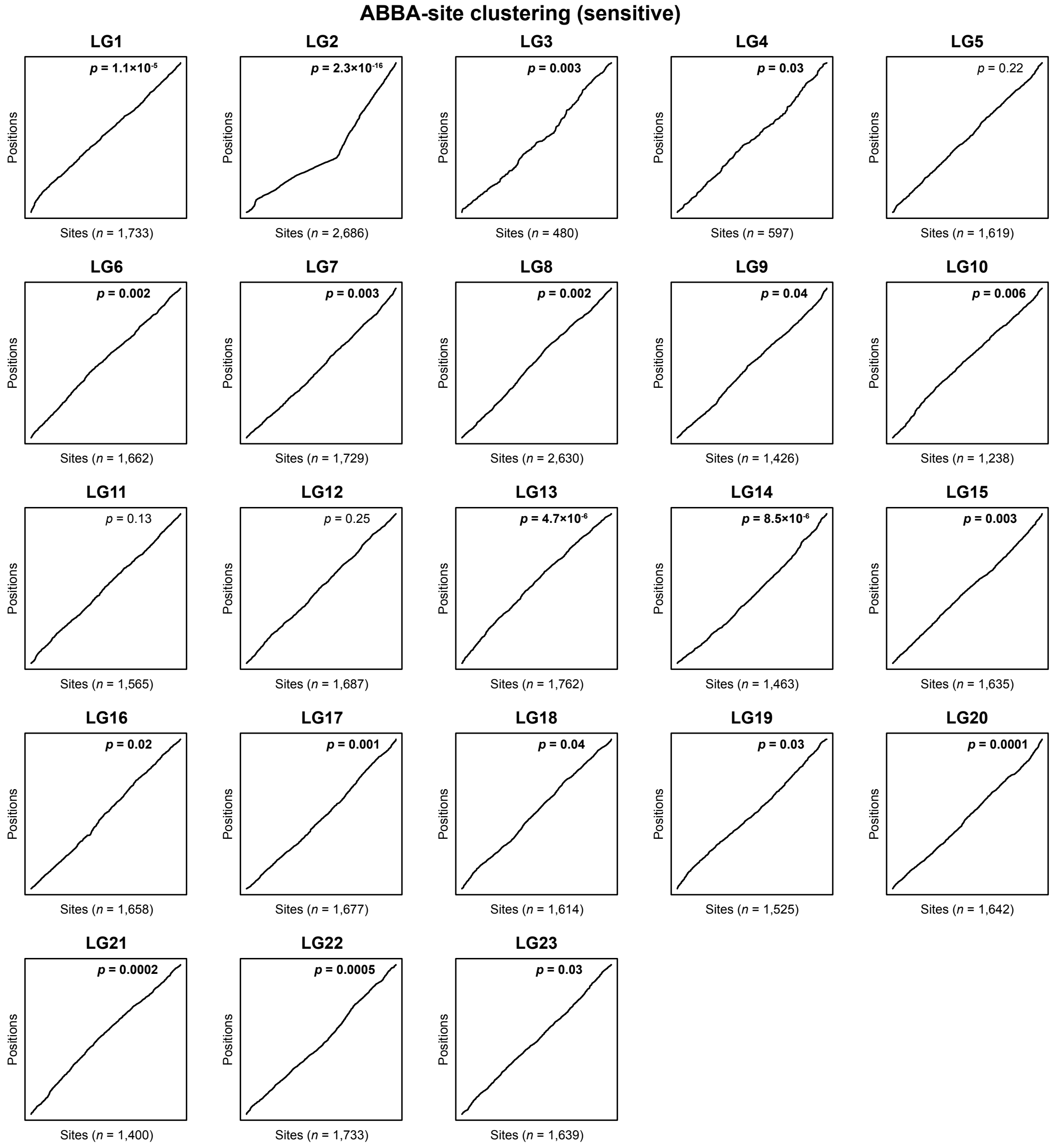

**Supplementary Figure S36:** Clustering of 'ABBA' sites in empirical data for Lake Tanganyika cichlid fishes, using the "robust" version of the 'ABBA'-site-clustering test. Results are shown per linkage group of the Nile tilapia reference assembly. With sorted "strong ABBA sites" on the horizontal axis, the black line indicates their position within a vector of "strong ABBA sites" and "strong BABA sites" on the vertical axis. A straight, diagonal line therefore illustrates a homogeneous distribution of these sites within this vector, while changes in the gradient illustrate clustering. A single significant $p$-value for LG2 is marked in bold.

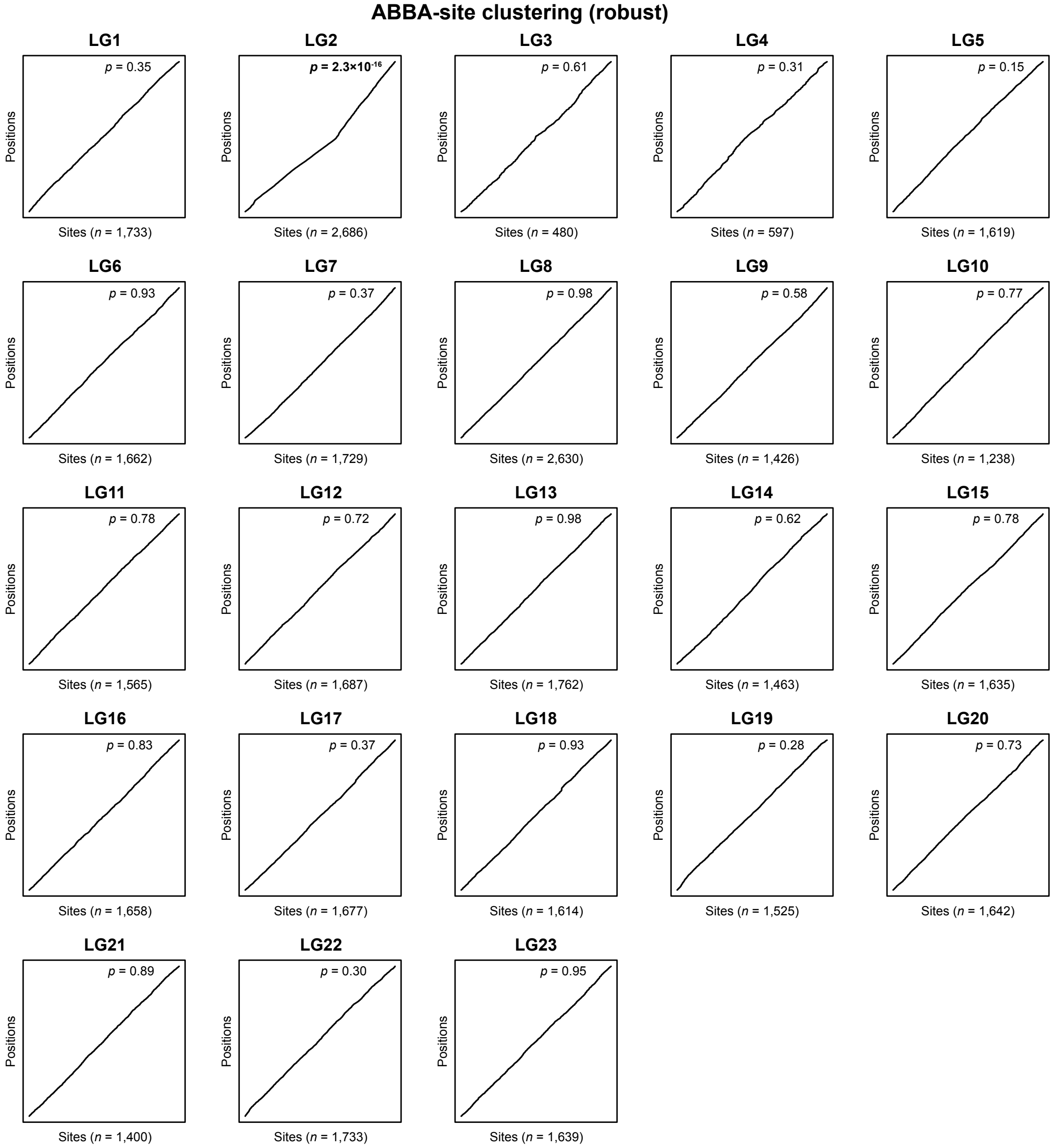

### Supplementary Tables

**Supplementary Table S1:** Signals of introgression detected with all applied methods, mean lengths of c-genes for all simulated datasets.
See separate file **Supplementary Table S1.xlsx**

**Supplementary Table S2:** Tree discordance with rate variation.
Maximum-likelihood trees were inferred with IQ-TREE for sets of 5,000, 2,000, or 1,000 alignments of 200, 500, or 1,000 bp, respectively. Here, results are shown only for datasets simulated without introgression ($m = 0$). Mean numbers of discordant trees were calculated across 10–50 analyses of replicate simulated datasets, indicating the numbers of trees in which not P1 and P2 appear as sister species, but P2 and P3 ($C_{\mathrm{P2,P3}}$) or P1 and P3 ($C_{\mathrm{P1,P3}}$). Where $C_{\mathrm{P2,P3}} > C_{\mathrm{P1,P3}}$ or vice versa, these numbers are given in bold.

| Divergence time ($t_{\mathrm{P1,P2}}$) | P2 rate ($s$) | Introgression rate ($m$) | Alignment length | # replicates | Mean values | |
|---|---|---|---|---|---|---|
| | | | | | $C_{\mathrm{P2,P3}}$ | $C_{\mathrm{P1,P3}}$ |
| 10 M | 0.25 | 0 | 200 | 10 | **64.9** | 55.2 |
| 10 M | 0.25 | 0 | 500 | 50 | 1.4 | 1.4 |
| 10 M | 0.25 | 0 | 1,000 | 10 | 0.0 | 0.0 |
| 20 M | 0.25 | 0 | 200 | 10 | **139.0** | 127.0 |
| 20 M | 0.25 | 0 | 500 | 50 | **4.2** | 4.0 |
| 20 M | 0.25 | 0 | 1,000 | 10 | 0.0 | 0.0 |
| 30 M | 0.25 | 0 | 200 | 10 | **217.7** | 203.9 |
| 30 M | 0.25 | 0 | 500 | 50 | **10.5** | 10.0 |
| 30 M | 0.25 | 0 | 1,000 | 10 | 0.1 | 0.1 |
| 10 M | 0.5 | 0 | 200 | 10 | **66.6** | 62.0 |
| 10 M | 0.5 | 0 | 500 | 50 | **2.1** | 1.7 |
| 10 M | 0.5 | 0 | 1,000 | 10 | 0.0 | 0.0 |
| 20 M | 0.5 | 0 | 200 | 10 | **144.3** | 138.0 |
| 20 M | 0.5 | 0 | 500 | 50 | **5.2** | 4.7 |
| 20 M | 0.5 | 0 | 1,000 | 10 | 0.0 | 0.0 |
| 30 M | 0.5 | 0 | 200 | 10 | **227.7** | 223.1 |
| 30 M | 0.5 | 0 | 500 | 50 | **13.0** | 10.6 |
| 30 M | 0.5 | 0 | 1,000 | 10 | 0.0 | **0.3** |
| 10 M | 2 | 0 | 200 | 10 | **92.5** | 89.9 |
| 10 M | 2 | 0 | 500 | 50 | 1.1 | **2.5** |
| 10 M | 2 | 0 | 1,000 | 10 | 0.0 | 0.0 |
| 20 M | 2 | 0 | 200 | 10 | 187.9 | **189.1** |
| 20 M | 2 | 0 | 500 | 50 | 6.6 | **6.9** |
| 20 M | 2 | 0 | 1,000 | 10 | 0.1 | 0.1 |
| 30 M | 2 | 0 | 200 | 10 | **314.8** | 307.1 |
| 30 M | 2 | 0 | 500 | 50 | 23.3 | **25.1** |
| 30 M | 2 | 0 | 1,000 | 10 | **1.2** | 0.7 |
| 10 M | 4 | 0 | 200 | 10 | 112.7 | **114.4** |
| 10 M | 4 | 0 | 500 | 50 | 2.7 | 2.7 |
| 10 M | 4 | 0 | 1,000 | 10 | 0.0 | 0.0 |
| 20 M | 4 | 0 | 200 | 10 | 236.3 | **265.1** |
| 20 M | 4 | 0 | 500 | 50 | 15.5 | **15.9** |
| 20 M | 4 | 0 | 1,000 | 10 | **0.6** | 0.2 |
| 30 M | 4 | 0 | 200 | 10 | 432.4 | **452.4** |
| 30 M | 4 | 0 | 500 | 50 | 47.0 | **52.9** |
| 30 M | 4 | 0 | 1,000 | 10 | **3.7** | 3.3 |

**Supplementary Table S3:** Tree discordance with introgression.
Maximum-likelihood trees were inferred with IQ-TREE for sets of 5,000, 2,000, or 1,000 alignments of 200, 500, or 1,000 bp, respectively. Here, results are shown only for datasets simulated with introgression (between P2 and P3) but without rate variation ($s = 1$). Mean numbers of discordant trees were calculated across 10–50 replicate simulated datasets, indicating the numbers of trees in which not P1 and P2 appear as sister species, but P2 and P3 ($C_{\mathrm{P2,P3}}$) or P1 and P3 ($C_{\mathrm{P1,P3}}$). Where $C_{\mathrm{P2,P3}} > C_{\mathrm{P1,P3}}$ or vice versa, these numbers are given in bold.

| Divergence time ($t_{\mathrm{P1,P2}}$) | P2 rate ($s$) | Introgression rate ($m$) | Alignment length | # replicates | Mean values $C_{\mathrm{P2,P3}}$ | Mean values $C_{\mathrm{P1,P3}}$ |
|---|---|---|---|---|---|---|
| 10 M | 1 | $10^{-9}$ | 200 | 10 | **95.2** | 73.2 |
| 10 M | 1 | $10^{-9}$ | 500 | 10 | **7.9** | 1.7 |
| 10 M | 1 | $10^{-9}$ | 1,000 | 10 | **2.0** | 0.1 |
| 20 M | 1 | $10^{-9}$ | 200 | 10 | **175.1** | 166.8 |
| 20 M | 1 | $10^{-9}$ | 500 | 10 | **13.6** | 6.2 |
| 20 M | 1 | $10^{-9}$ | 1,000 | 10 | **2.6** | 0.3 |
| 30 M | 1 | $10^{-9}$ | 200 | 10 | **269.1** | 245.2 |
| 30 M | 1 | $10^{-9}$ | 500 | 10 | **21.7** | 16.8 |
| 30 M | 1 | $10^{-9}$ | 1,000 | 10 | **3.3** | 0.2 |
| 10 M | 1 | $10^{-8}$ | 200 | 10 | **233.9** | 85.1 |
| 10 M | 1 | $10^{-8}$ | 500 | 50 | **71.7** | 5.5 |
| 10 M | 1 | $10^{-8}$ | 1,000 | 10 | **29.8** | 0.5 |
| 20 M | 1 | $10^{-8}$ | 200 | 10 | **319.7** | 181.9 |
| 20 M | 1 | $10^{-8}$ | 500 | 50 | **70.0** | 10.6 |
| 20 M | 1 | $10^{-8}$ | 1,000 | 10 | **34.8** | 1.3 |
| 30 M | 1 | $10^{-8}$ | 200 | 10 | **408.8** | 273.0 |
| 30 M | 1 | $10^{-8}$ | 500 | 50 | **81.8** | 22.8 |
| 30 M | 1 | $10^{-8}$ | 1,000 | 10 | **34.5** | 3.2 |
| 10 M | 1 | $10^{-7}$ | 200 | 10 | **1356.1** | 170.4 |
| 10 M | 1 | $10^{-7}$ | 500 | 50 | **570.6** | 42.6 |
| 10 M | 1 | $10^{-7}$ | 1,000 | 10 | **271.1** | 9.4 |
| 20 M | 1 | $10^{-7}$ | 200 | 10 | **1364.4** | 303.0 |
| 20 M | 1 | $10^{-7}$ | 500 | 50 | **556.3** | 57.0 |
| 20 M | 1 | $10^{-7}$ | 1,000 | 10 | **269.8** | 15.5 |
| 30 M | 1 | $10^{-7}$ | 200 | 10 | **1388.3** | 433.5 |
| 30 M | 1 | $10^{-7}$ | 500 | 50 | **554.1** | 79.7 |
| 30 M | 1 | $10^{-7}$ | 1,000 | 10 | **279.0** | 21.3 |
| 10 M | 1 | $10^{-6}$ | 200 | 10 | **3672.3** | 276.1 |
| 10 M | 1 | $10^{-6}$ | 500 | 10 | **1754.4** | 78.0 |
| 10 M | 1 | $10^{-6}$ | 1,000 | 10 | **938.9** | 19.1 |
| 20 M | 1 | $10^{-6}$ | 200 | 10 | **3549.4** | 489.6 |
| 20 M | 1 | $10^{-6}$ | 500 | 10 | **1654.2** | 116.7 |
| 20 M | 1 | $10^{-6}$ | 1,000 | 10 | **906.3** | 31.3 |
| 30 M | 1 | $10^{-6}$ | 200 | 10 | **3374.3** | 608.2 |
| 30 M | 1 | $10^{-6}$ | 500 | 10 | **1599.1** | 145.6 |
| 30 M | 1 | $10^{-6}$ | 1,000 | 10 | **890.9** | 40.9 |

**Supplementary Table S4:** Genetic distances.
Pairwise genetic distances between species ($d_{\mathrm{xy}}$) were calculated from genomic datasets simulated with a population size $N_{\mathrm{e}} = 10^5$, a mutation rate $\mu = 2 \times 10^{-9}$, and a recombination rate $r = 10^{-8}$. Genetic distances are provided for the species pairs P1 and P2, $d_{\mathrm{xy}}$(P1,P2); P1 and P3, $d_{\mathrm{xy}}$(P1,P3); and P2 and P3, $d_{\mathrm{xy}}$(P2,P3).

| | | | Genetic distances | | |
|---|---|---|---|---|---|
| Divergence time ($t_{\mathrm{P1,P2}}$) | Introgression rate ($m$) | P2 rate ($s$) | $d_{\mathrm{xy}}$(P1,P2) | $d_{\mathrm{xy}}$(P1,P3) | $d_{\mathrm{xy}}$(P2,P3) |
| 10 M | 0 | 0.25 | 0.0253 | 0.0765 | 0.0629 |
| 20 M | 0 | 0.25 | 0.0491 | 0.1112 | 0.0853 |
| 30 M | 0 | 0.25 | 0.0720 | 0.1444 | 0.1071 |
| 10 M | 0 | 1 | 0.0397 | 0.0765 | 0.0764 |
| 20 M | 0 | 1 | 0.0765 | 0.1112 | 0.1113 |
| 30 M | 0 | 1 | 0.1113 | 0.1443 | 0.1443 |
| 10 M | 0 | 4 | 0.0942 | 0.0765 | 0.1281 |
| 20 M | 0 | 4 | 0.1755 | 0.1114 | 0.2052 |
| 30 M | 0 | 4 | 0.2465 | 0.1442 | 0.2724 |
| 10 M | $10^{-7}$ | 0.25 | 0.0327 | 0.0695 | 0.0468 |
| 20 M | $10^{-7}$ | 0.25 | 0.0563 | 0.1047 | 0.0697 |
| 30 M | $10^{-7}$ | 0.25 | 0.0789 | 0.1377 | 0.0917 |
| 10 M | $10^{-7}$ | 1 | 0.0465 | 0.0684 | 0.0599 |
| 20 M | $10^{-7}$ | 1 | 0.0834 | 0.1046 | 0.0962 |
| 30 M | $10^{-7}$ | 1 | 0.1181 | 0.1374 | 0.1291 |
| 10 M | $10^{-7}$ | 4 | 0.1007 | 0.0688 | 0.1131 |
| 20 M | $10^{-7}$ | 4 | 0.1814 | 0.1043 | 0.1920 |
| 30 M | $10^{-7}$ | 4 | 0.2518 | 0.1377 | 0.2611 |
| 10 M | $10^{-8}$ | 0.25 | 0.0261 | 0.0755 | 0.0609 |
| 20 M | $10^{-8}$ | 0.25 | 0.0501 | 0.1103 | 0.0830 |
| 30 M | $10^{-8}$ | 0.25 | 0.0727 | 0.1435 | 0.1054 |
| 10 M | $10^{-8}$ | 1 | 0.0408 | 0.0757 | 0.0744 |
| 20 M | $10^{-8}$ | 1 | 0.0774 | 0.1103 | 0.1092 |
| 30 M | $10^{-8}$ | 1 | 0.1123 | 0.1435 | 0.1425 |
| 10 M | $10^{-8}$ | 4 | 0.0949 | 0.0757 | 0.1261 |
| 20 M | $10^{-8}$ | 4 | 0.1764 | 0.1104 | 0.2031 |
| 30 M | $10^{-8}$ | 4 | 0.2470 | 0.1436 | 0.2710 |

**Supplementary Table S5:** ABBA-site clustering in empirical data.
Results of the ABBA-site-clustering test applied to empirical data, along with the $D$-statistic and its $p$-value. Both the $p$-values of the "sensitive" and "robust" versions of the ABBA-site-clustering test are reported. All statistics were calculated with the program Dsuite (Malinsky et al., 2020). The empirical dataset used in these analyses, was a subset of the variant call dataset by Ronco et al. (2021) for the Lake Tanganyika cichlid assemblage, mapped to the tilapia (*Oreochromis niloticus*) reference assembly with accession GCF_001858045.1 (Conte et al., 2017). LG: linkage group; $C_{\text{ABBA}}$: number of "ABBA" sites; $C_{\text{BABA}}$: number of "BABA" sites; $C_{\text{ABBA}}, s$: number of "strong ABBA" sites used in the ABBA-site-clustering test.

| | | | $D$-statistic | | | ABBA-site clustering test | |
|---|---|---|---|---|---|---|---|
| LG | $C_{\text{ABBA}}$ | $C_{\text{BABA}}$ | $D$ | $p$ | $C_{\text{ABBA}}, s$ | $p$ ("sensitive" version) | $p$ ("robust" version) |
| 1 | 1,805.8 | 1,770.3 | 0.010 | 0.6862 | 1,733 | $\mathbf{1.1 \times 10^{-5}}$ | 0.3553 |
| 2 | 2,771.8 | 1,541.5 | 0.285 | $\mathbf{1.8 \times 10^{-5}}$ | 2,686 | $\mathbf{2.3 \times 10^{-16}}$ | $\mathbf{2.3 \times 10^{-16}}$ |
| 3 | 513.7 | 457.5 | 0.058 | 0.0754 | 480 | **0.0034** | 0.6083 |
| 4 | 669.2 | 640.2 | 0.022 | 0.4992 | 597 | **0.0317** | 0.3132 |
| 5 | 1,713.1 | 1,626.1 | 0.026 | 0.2612 | 1,619 | 0.2198 | 0.1487 |
| 6 | 1,755.2 | 1,680.8 | 0.022 | 0.2192 | 1,662 | **0.0019** | 0.9256 |
| 7 | 1,817.4 | 1,704.5 | 0.032 | **0.0472** | 1,729 | **0.0032** | 0.3697 |
| 8 | 2,726.9 | 2,685.0 | 0.008 | 0.5246 | 2,630 | **0.0022** | 0.9822 |
| 9 | 1,492.4 | 1,387.1 | 0.037 | 0.0829 | 1,426 | **0.0408** | 0.5820 |
| 10 | 1,306.2 | 1,178.3 | 0.051 | **0.0017** | 1,238 | **0.0064** | 0.7861 |
| 11 | 1,633.2 | 1,444.3 | 0.061 | **0.0002** | 1,565 | 0.1270 | 0.7805 |
| 12 | 1,744.6 | 1,674.5 | 0.021 | 0.2067 | 1,687 | 0.2533 | 0.7161 |
| 13 | 1,853.7 | 1,815.7 | 0.010 | 0.4950 | 1,762 | $\mathbf{4.7 \times 10^{-6}}$ | 0.9798 |
| 14 | 1,512.1 | 1,456.6 | 0.019 | 0.3991 | 1,463 | $\mathbf{8.5 \times 10^{-6}}$ | 0.6170 |
| 15 | 1,697.1 | 1,627.0 | 0.021 | 0.2541 | 1,635 | **0.0031** | 0.7825 |
| 16 | 1,722.2 | 1,679.5 | 0.013 | 0.3725 | 1,658 | **0.0286** | 0.8308 |
| 17 | 1,746.0 | 1,665.2 | 0.024 | 0.2820 | 1,677 | **0.0013** | 0.3679 |
| 18 | 1,702.1 | 1,688.5 | 0.004 | 0.8431 | 1,614 | **0.0411** | 0.9308 |
| 19 | 1,583.1 | 1,572.9 | 0.003 | 0.9055 | 1,525 | **0.0269** | 0.2753 |
| 20 | 1,704.0 | 1,679.8 | 0.007 | 0.7253 | 1,642 | **0.0002** | 0.7263 |
| 21 | 1,469.6 | 1,442.6 | 0.009 | 0.6425 | 1,400 | **0.0002** | 0.8945 |
| 22 | 1,818.1 | 1,813.9 | 0.001 | 0.9642 | 1,733 | **0.0005** | 0.3032 |
| 23 | 1,709.5 | 1,679.1 | 0.009 | 0.6521 | 1,639 | 0.3064 | 0.9455 |
| All | 38,467.0 | 35,910.9 | 0.012 | 0.3953 | 36,800 | $\mathbf{2.3 \times 10^{-16}}$ | $\mathbf{7.8 \times 10^{-9}}$ |
